# Development of Oxadiazolone Activity-Based Probes Targeting FphE for Specific Detection of *S. aureus* Infections

**DOI:** 10.1101/2023.12.11.571116

**Authors:** Jeyun Jo, Tulsi Upadhyay, Emily C. Woods, Ki Wan Park, Nichole J. Pedowitz, Joanna Jaworek-Korjakowska, Sijie Wang, Tulio A. Valdez, Matthias Fellner, Matthew Bogyo

**Author notes:** Correspondence (M.B.). These authors contributed equally (Jeyun Jo, Tulsi Upadhyay).

## Abstract

*Staphylococcus aureus* is a major human pathogen responsible for a wide range of systemic infections. Since its propensity to form biofilms *in vivo* poses formidable challenges for both detection and treatment, tools that can be used to specifically image *S. aureus* biofilms are highly valuable for clinical management. Here we describe the development of oxadiazolone-based activity-based probes to target the *S. aureus*-specific serine hydrolase FphE. Because this enzyme lacks homologs in other bacteria, it is an ideal target for selective imaging of *S. aureus* infections. Using X-ray crystallography, direct cell labeling and mouse models of infection we demonstrate that oxadiazolone-based probes enable specific labeling of *S. aureus* bacteria through the direct covalent modification of the FphE active site serine. These results demonstrate the utility of the oxadizolone electrophile for activity-based probes (ABPs) and validate FphE as a target for development of imaging contrast agents for the rapid detection of *S. aureus* infections.

## Introduction

*Staphylococcus aureus* is a ubiquitous commensal bacterium that colonizes approximately 30% of the human population.^1^ It plays a dual role as both a harmless inhabitant of human skin and mucosal epithelia and a major human pathogen contributing to a diverse spectrum of clinical manifestations.^2^ Clinical diseases caused by *S. aureus* range from uncomplicated superficial skin and soft tissue infections (SSTI) to severe infections such as infective endocarditis (IE) or life-threatening bacteremia.^3^ The management strategy for these syndromes varies depending on the infection site and diagnostic evaluation.^4–7^ The clinical presentation of *S. aureus* infections varies considerably,^8^ and the propensity of the bacteria to form biofilms further complicates both diagnosis of infections and monitoring of therapy response.^9, 10^ In addition, the alarming rise in antibiotic-resistant *S. aureus* strains including methicillin-resistant *Staphylococcus aureus* (MRSA) further necessitates the development of early and accurate diagnostic strategies for effective infection control.^11^ Such strategies need to be both highly sensitive for disease detection and highly specific to distinguish *S. aureus* from other microbial species that may cause similar clinical symptoms.

Serine hydrolases are one of the largest functional enzyme classes in most organisms, with highly diverse distribution across all three kingdoms of life.^12^ Enzymes in this family not only play pivotal roles in regulating diverse biological processes in mammals, but also often play key functional roles in pathogenic organisms, including bacteria and viruses.^13, 14^ Because serine hydrolases can play essential roles in normal cellular functions as well as in drug resistance and virulence, they have great potential as targets of therapeutics for a range of diseases.^14–17^ Serine hydrolases can be sub-divided into two groups based on overall function. One group is made up of proteases primarily involved in the hydrolysis of peptide bonds within proteins. The other is made up of metabolic hydrolases such as peptidases, amidases, lipases, esterases, and thioesterases that process amide, ester, or thioester bonds in small-molecular metabolites, peptides, or post-translational modifications on proteins.^12, 13^ While some members of metabolic serine hydrolases in human have been extensively characterized and validated as targets for clinically approved drugs,^14^ the potential of these enzymes as pharmacological or diagnostic targets in *S. aureus* has yet to be established.

Because serine hydrolases all use a highly nucleophilic active site serine, it is possible to use reactive electrophiles such as fluorophosphonates (FPs) to develop activity-based probes (ABPs) to label and track their activity in live cells and even whole organisms.^13^ FP-based probes enable global profiling of serine hydrolases across various organisms by a method known as activity-based protein profiling (ABPP).^18–25^ Using the broadly reactive fluorphosphonate probe, FP tetramethylrhodamine (FP-TMR), we recently identified a family of metabolic serine hydrolases in *S. aureus* that we named fluorophosphonate-binding hydrolases (Fphs; FphA–J).^26^ Among the Fphs, we found that the lipid esterase FphB is likely a virulence factor important for infection in the heart and liver *in vivo*.^26^ In addition, recent structural and biochemical studies of FphF and FphH have begun to shed light on possible functions of these enzymes.^27, 28^ The importance of FphB and FphH in mediating systemic *S. aureus* infections, makes them potential targets for development of therapeutic agents.^26, 28^ In order to develop therapeutic and diagnostic agents for *Staphylococcus* infections, it is optimal to select targets that are unique to this organism over other highly related bacterial species.^29^ In particular, FphE fits this criterion as it is the least conserved Fph protein across 50 species within the *Staphylococcus* genus.^30, 31^

While FP probes have proven to be particularly valuable due to their broad reactivity across diverse serine hydrolases, ABPs tailored to react with specific targets are often essential for applications such as cellular or *in vivo* imaging or profiling proteins with low expression levels that are challenging to detect with broad-spectrum probes.^32–40^ The development of target-selective ABPs often begins with selection of a suitable reactive electrophilic ‘warhead’ which forms a covalent bond to an active site nucleophile.^32, 41^ The electrophile itself can generate high selectivity towards specific nucleophiles and enzyme classes.^41^ Recently, a screen of inhibitors of the MRSA strain USA300 *S. aureus* identified compounds containing a 1,3,4-oxadiazole-2(3*H*)- one (oxadiazolone) electrophile,^42^ which has been used primarily to target lipid hydrolytic enzymes.^43–56^ Interestingly, when the most potent optimized hit from the MRSA screen was converted into an alkyne labeled probe, it labeled multiple Fph serine hydrolases as well as the highly conserved Fatty Acid Synthesis II (FASII) pathway enzyme FabH,^57^ and several other poorly characterized enzymes. These targets included enzymes that use both serine and cysteine as catalytic residues, suggesting that the oxadiazolone electrophile, while suitable for use as an ABP, has overall broad reactivity that needs to be tuned to generate target-specific probes.

Herein, we describe the design and synthesis of ABPs containing the oxadiazolone electrophile that have been optimized for selective targeting of FphE in *S. aureus*. Structural analysis of the co-crystal complex of FphE with one of our probes confirmed an irreversible covalent binding mechanism involving the active site serine. These structural studies also uncovered a unique type of homodimer interface with the catalytic triad formed by two monomeric units. Furthermore, we found that addition of a fluorescent label to our optimized oxadiazolone probe resulted in high selectivity for FphE in live *S. aureus* cells. This probe also enabled selective labeling of infections sites in a *S. aureus* mouse infection model. These results demonstrate the utility of the oxadiazolone electrophile for specific targeting of serine hydrolases in *S. aureus* and provides support for FphE as a target for *S. aureus*-specific imaging probes. We believe these probes have the potential to ultimately improve both diagnosis and management of *S. aureus* infections.

## Results and discussion

### Serine hydrolase FphE is unique to S. aureus

*Staphylococcus aureus* expresses a total of 10 Fph proteins with a range of homologs in other bacterial species.^30, 31^ In order to identify which Fph enzyme might be the best targets for probes that could uniquely identify *S. aureus,* we searched for the presence of Fph homologs from the USA300 MRSA strain in 30 different species using protein BLAST (Figure 1A and Table S1). We focused the BLAST search on the most common mortality-associated bacterial pathogens,^58^ because these would be the most likely species that would be relevant to distinguish in clinical infections. We identified homologs of Fph proteins mainly in Gram-positive bacteria, with the only outlier being *Listeria monocytogenes*. Of all the Fph enzymes only FphE is found in less than 50% of the 47 different species of the genus *Staphylococcus*, and notably, it is the only Fph absent in the representative nosocomial coagulase-negative staphylococci (CoNS), *S. epidermidis* and *S. haemolyticus*.^26, 30, 59, 60^ Thus, specific probes for FphE may offer opportunities for the development of imaging agents that can potentially discriminate infections caused by *S. aureus* from those caused by other pathogenic bacteria including CoNS, which has different and less aggressive clinical manifestations.^61, 62^

**Figure 1.**
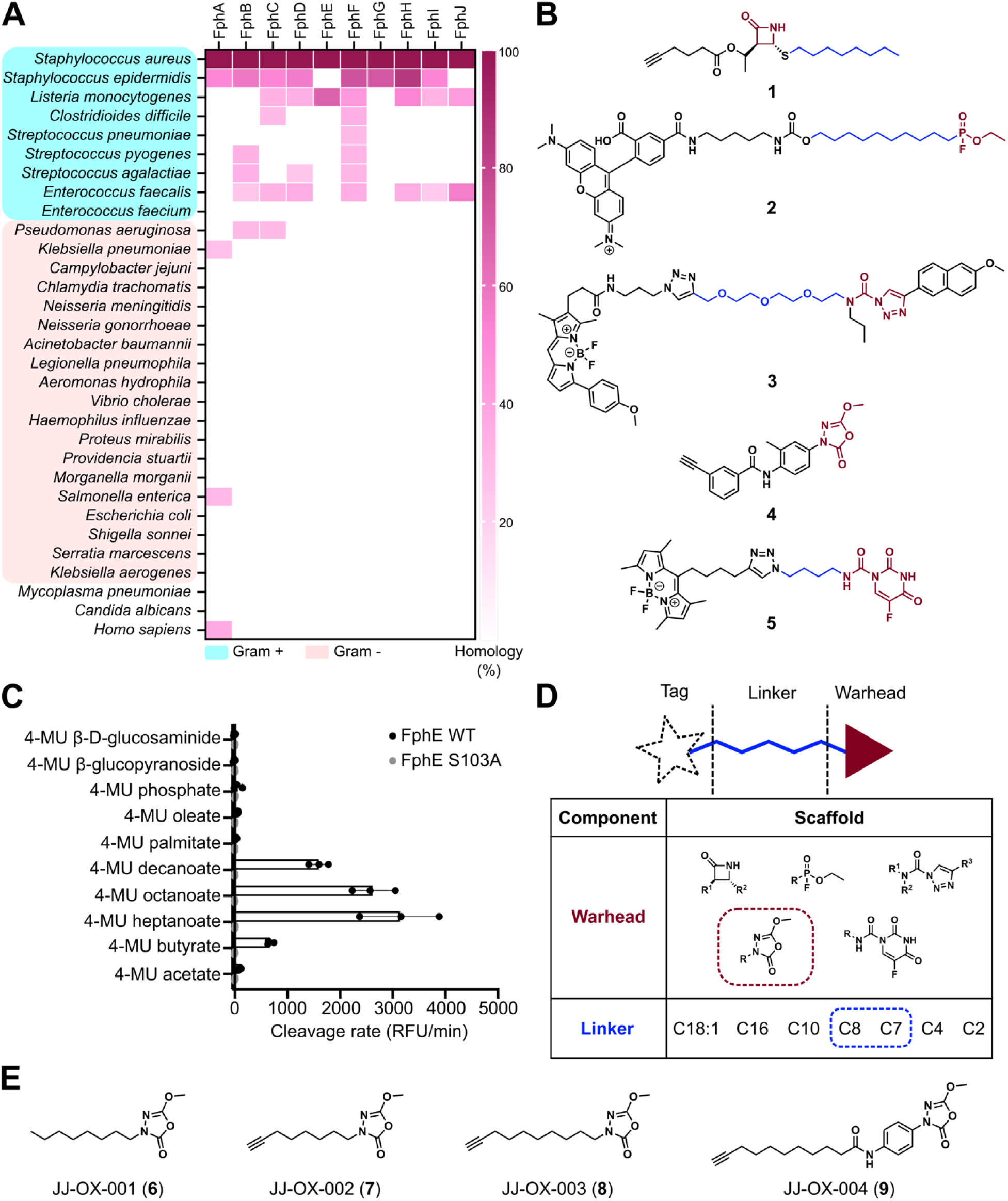
Rational design of activity-based probes targeting FphE. (A). Heat map indicating the conservation of Fph proteins among different clinically relevant microorganisms. The color gradient indicates the percentage homology with the *S. aureus* Fph, as determined by a threshold e-value of 1 x 10^-10^. (B) Structures of previously reported activity-based probes (ABPs) **1**–**5** capable of binding to FphE. Covalent warheads are highlighted in maroon, and chains directly attached to the warhead are highlighted in blue. (C) Cleavage rate (relative fluorescent unit (RFU)/minute) of 4-methylumbelliferone (4-MU) substrates by FphE wild-type (WT, black) and catalytic-dead mutant (S103A, gray). Each dot represents an individual experiment, and bars represent mean ± standard deviation (*n*LJ=LJ3). (D) Mix and match panel of alkyl chains and warheads to design FphE targeting ABPs. (E) Molecular structures of oxadiazolone-based covalent inhibitor (**6**) and ABPs (**7**–**9**). See the Supporting Information for detailed chemical synthesis.

### Rational design of FphE targeting activity-based probes

In an effort to develop selective activity-based probes (ABPs) that can serve as chemical tools to further characterize FphE, we needed to select an optimal electrophile. Among the various moieties capable of directly forming a covalent bond with the active site nucleophile of serine hydrolases, five classes including β-lactam (**1**), fluorophosphonate (**2**), triazole urea (**3**), oxadiazolone (**4**), and 1-carbamoyl-5-fluorouracil (**5**) have been successfully used in irreversible inhibitors or probes that label FphE (Figure 1B).^26, 42, 63–65^ In particular, the 1,3,4-oxadiazole-2(3*H*)-one (oxadiazolone) moiety in **4** has the potential to be tuned for selectivity for FphE. This class of electrophile was recently reported in a screen for compounds that kill MRSA bacteria. In addition, an optimized hit containing the oxadiazolone electrophiles was converted into an ABP that effectively labeled multiple Fph proteins in live MRSA bacteria.^42^ However, this ABP also labeled other targets including an aldehyde dehydrogenase and a fatty acid biosynthetic enzyme, both of which use a cysteine nucleophile. We therefore reasoned that the oxadiazolone could be used to make an effective APB for FphE if we could tune the selectivity of the probe by tailoring the linker to the preferences of FphE. We therefore determined the substrate preference of recombinantly expressed and purified FphE using a library of 4-methylumbelliferone (4-MU) substrates (Figure 1C). We found that wild-type FphE hydrolyzed saturated lipid esters ranging in chain length from C4 to C10, showing a particularly strong preference for the C7 and C8 chain lengths. This cleavage activity was lost upon mutation of serine-103 to alanine, suggesting that serine-103 is the catalytic active site residue.

We selected alkyl chain lengths of 7 or 8 and the oxadiazolone warhead to design our first set of potential FphE-specific probes (Figure 1D). This included the unlabeled inhibitor (**6**) and probes (**7**–**9**; Figure 1E). Compound **9** was designed to include the *N*-phenylamide moiety present in compound **4** between the lipid chain and warhead of **8**. Because the methyl group of the phenyl ring plays a crucial role in inducing *S. aureus* killing and the fact that FphE is a non-essential protein, we removed the methyl group in our probe **9**.^26, 42^

### JJ-OX-004 (**9**) as an irreversible covalent probe that selectively binds to the active serine in FphE

To assess the potential of oxadiazolones (**6**–**9**), we measured their inhibitory activity against recombinant FphE using 4-MU octanoate as a substrate (Figure 2A). Two alkyne probes, **8** and **9,** had sub-micromolar potency, with compound **9** displaying the most potent activity with an IC_50_ of 19.9 nM. Notably, there was a large difference in IC_50_ values between compounds **7** (> 50 µM) and **8** (187 nM), which only differ in their chain lengths by two carbons. This result contrasts with the analysis of the 4-MU substrate preference of FphE, which suggested some tolerance to carbon length (Figure 1B). This discrepancy implies that the inhibition of FphE by compound **8** may not be solely due to its binding with the catalytic active serine. Therefore, we assessed whether probes **8** and **9** labeled FphE through the active site serine (Ser103). Direct labeling of recombinant FphE confirmed that probe **8** binds much less effectively than probe **9** (Figure 2B). Interestingly, probe **8** also effectively labeled the catalytically dead mutant (S103A) suggesting that it may bind through interactions beyond the active site serine. Therefore, we chose to focus on probe **9** as it showed the highest potency and specific labeling that was dependent on the active site serine-103.

**Figure 2.**
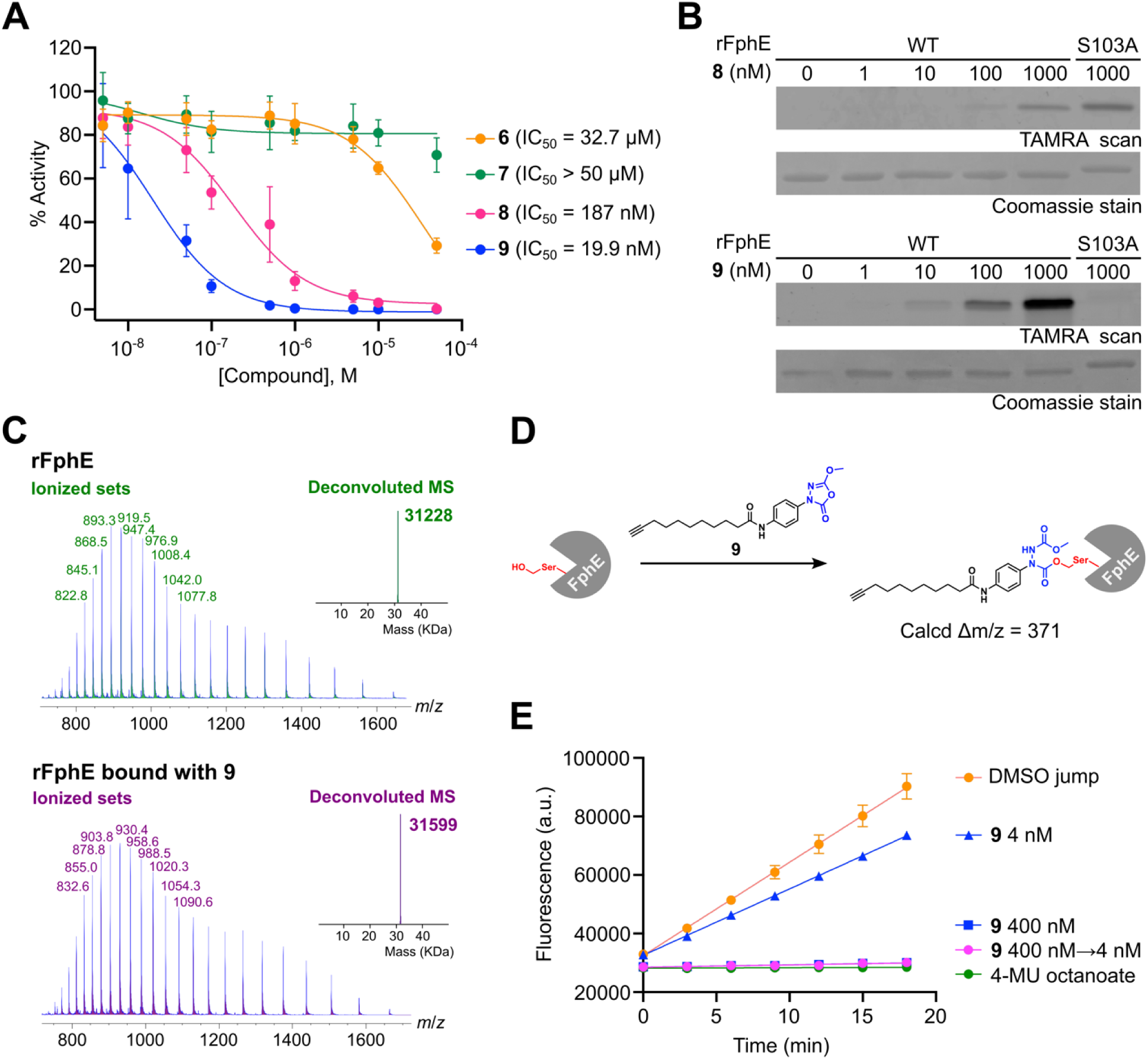
Selection and biochemical characterization of **9**. (A) IC_50_ determination of **6**–**9** against recombinant FphE (rFphE, 0.5 nM) measured by initial cleavage rate of 4-MU octanoat substrate (20 µM). Percent activity was calculated by comparing to a DMSO control, with point and bars representing mean ± standard deviation (*n*L=L6). (B) Concentration-dependent labeling of WT and S103A rFphE by clickable probes **8** or **9**. WT or S103A rFphE (300 ng) were incubated with each probe for one hour, labeled with TAMRA-azide by CuAAC reaction, and analyzed by SDS-PAGE/fluorescence scan. (C) Deconvoluted mass spectra of rFphE before (green) and after (purple) treatment of **9**. (D) Proposed mechanism by which **9** covalently binds to FphE. (E) Fluorescence progress curves for substrate hydrolysis activity of rFphE after jump dilution.

To confirm that probe **9** binds to and inhibits FphE through an irreversible covalent mechanism, we performed both a jump dilution assay and direct mass spectrometry-based analysis of the inhibited protein. We prepared the FphE-probe complex by incubating recombinant FphE with **9** to allow covalent modification of the protein. Mass spectrometry of the complex confirmed the observed increase in deconvoluted mass of 371 Da (Figure 2C), corresponding to the expected shift upon formation of the covalent bond between **9** and serine-103 via the proposed binding mechanism (Figure 2D).^42, 43, 51, 54, 55^ To further confirm the irreversibility of the modification, we used a jump dilution assay^66^ in which we incubated recombinant FphE with **9** at a concentration of 400 nM (20-fold of IC_50_) to ensure full occupation of the active site followed by 100-fold dilution into a fluorogenic substrate (4-MU octanoate) solution (Figure 2E). The progress curve demonstrated that FphE remains inhibited after dilution, confirming that it is an irreversible binder.

### Structural basis of covalent inhibitory mechanism of FphE by **9**

To gain further insights into FphE, we determined the protein crystal structures of unbound FphE (PDB ID 8T87 at 1.62 Å resolution) as well as FphE with **9** covalently bound (8T88 at 1.54 Å). FphE always formed dimeric crystals in a unique arrangement. The two dimer copies of FphE are assembled by N-terminal residues 1-136 of one chain and C-terminal residues 167-276 of the other chain with a long helix formed by residues 137-166 in an antiparallel helix bundle connecting the two “monomers” (Figure 3A). FphE is a member of the well-studied α/β hydrolase superfamily^67^ which normally contain a single core β-sheet connected by several α helices. Oligomers have been reported, however these are always formed by monomer surface interfaces, with FphF being an example.^27^ Alphafold^68^ also predicts a monomeric fold (Figure 3A) and we were unable to reproduce the observed dimer using Alphafold. The overall fold of the two copies of chain A and B of FphE in the determined dimer closely resemble the predicted monomer. However, the rearrangement of the helical area at residues 137-166 narrows access to the active site in the dimer, making the entrance more hydrophobic and completely restricting access from one side due to the nearby second copy of FphE. The two copies of FphE within the crystal structures closely resemble each other.

**Figure 3.**
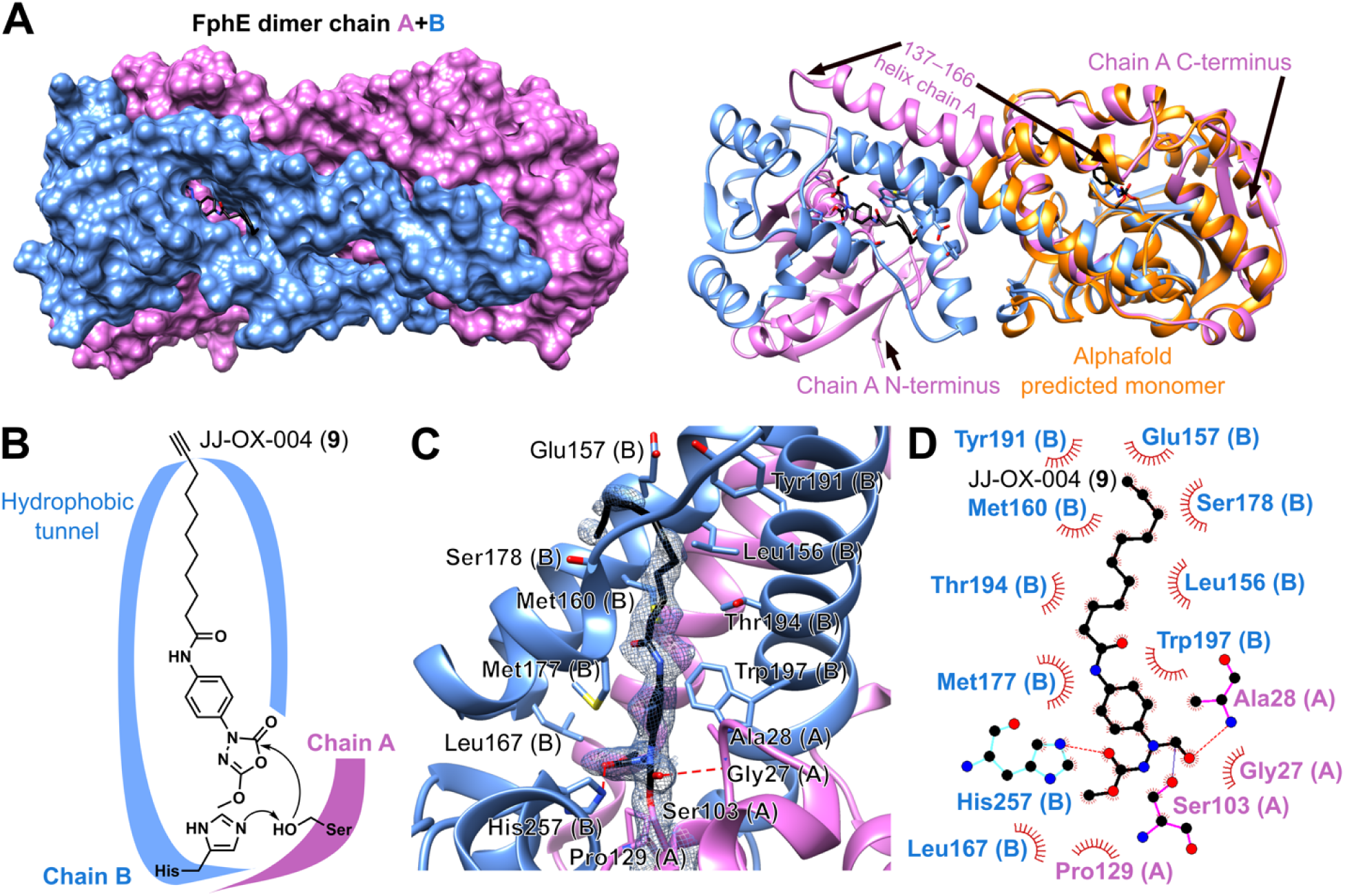
Structure of FphE in complex with **9** (JJ-OX-004). (A) Crystal structure of dimeric FphE (PDB ID 8T88 - **9** bound at 1.54 Å resolution; chain A colored in purple, chain B in blue) as surface representation or ribbon representation. Alphafold AF-Q2FDS6-F1 predicted monomeric form is shown as orange ribbons for comparison. (B) Schematic view of proposed covalent binding mode of **9** with FphE. (C) Close-up of oxadiazolone ABP **9** covalently bound to FphE-Ser103. The 2*F_o_−F_c_* electron density map for **9** and Ser103 of chain A is shown as a blue mesh at 0.5 σ. Hydrophobic tunnel residues are shown, as well as two hydrogen bond partners (red dashed lines). (D) LigPlot 2D illustration of interactions of **9** with specific residues of FphE.

The FphE serine hydrolase catalytic triad consists of Ser103, His257, and Glu127. Due to the unusual dimer arrangement, the triad is formed by Ser103 and Glu127 from chain A and His257 from chain B (Figure 3B). The active site is connected to the protein-surface via a hydrophobic tunnel (Figure 3B) formed by residues of chain B. The changes to the active site pocket and the tunnel upon binding of **9** are minimal, with only a couple of residues adopting slightly different side chain orientations compared to the unbound structure (Figure S2A). This suggests that the dimeric structure of FphE is a very good fit for **9** and binds the compound tightly without the need for major rearrangements, which supports the substrate specificity results.

This structure also confirms the proposed reaction mechanism between compound **9** and FphE (Figure 3B).^28, 42, 43, 51, 54, 55^ The electron density clearly shows the covalent bond between the terminal oxygen atom of active site Ser103 and the opened oxadiazolone ring (Figure 3C). The two oxygen atoms of the opened ring are stabilized in the structure by hydrogen bonds to the nitrogen ring atom of His257 and the backbone nitrogen of Ala28 (Figure 3C and 3D). The oxygen atom of **9** that forms a hydrogen bond with Ala28 occupies the space near the active site serine that has been described as the oxyanion hole ^27, 69^ for this protein family indicating that binding of **9** most likely resembles how oxadiazolones in general will bind and react with α/β hydrolase-fold enzymes. The opened oxidazolone ring of **9** is further stabilized by a small network of hydrogen bonds to several water molecules that connect it to other residues (Figure S2B). This hydrogen bonding network near Ser103 is also observed in the unbound structure (Figure S2C), with one water atom moving into the oxyanion hole. The hydrophobic tunnel is completely empty in the unbound structure, including any water molecules, suggesting that probe **9** likely mimics the yet to be identified biological substrate of FphE. The structure also gives an indication why the benzene ring in **9** improves potency. The ring atoms in **9** have hydrophobic interactions with FphE residues Ala28, Met177 and Trp197 (Figure 3C and 3D). The ring optimally bridges the space between Met17 and Trp197 with hydrogen-hydrogen distances as short as 2.3 Å (Figure S2D). In addition, the terminal methyl group of the oxidazolone also makes hydrophobic interactions with Trp197, as well as Leu167 (Figure S2D). Beyond the aromatic ring, the remaining lipid tail of **9** occupies the hydrophobic channel with several more residues (Leu156, Met160, Thr194) forming hydrophobic interactions with most atoms of the lipid chain (Figure 3C and D). Additional hydrophobic interactions with **9** are found for side chains of Glu157, Ser178 and Tyr191 at the surface of the protein. Electron density fit deteriorates for the atoms of **9** that are closer to the surface, indicating some flexibility/disorder in this region.

### Analysis of cellular localization of FphE by live S. aureus labeling with **9**

After confirming that **9** is a covalent probe for FphE through enzymatic and crystallographic analysis, we determined the labeling specificity of the probe in live *S. aureus* cells (strain USA300). We treated cells with **9** followed by cell lysis and then click labeling using TAMRA-azide with analysis by SDS-PAGE (Figure 4A, top row). We also analyzed the cellular localization of FphE by performing a protease protection assay in which only extracellular enzymes are digested by proteinase K, followed by activity-based probe profiling (ABPP; Figure 4A, bottom row).^70–72^ FphE was labeled by **9** in a dose-dependent manner, displaying high selectivity for the target protein at a concentration of 100 nM (Figure 4B). We treated purified recombinant FphE with PK (Figure 4C), which confirmed that it is sensitive to PK treatment. However, FphE remains intact even after 16 h of PK treatment in live *S. aureus*, suggesting it has an intracellular location. In contrast, the faint labeling of FphB (confirmed through transposon mutant of *fphB* (Figure 4D, lane 5 and 6)) was lost following PK treatment (Figure 4D), consistent with its reported localization on the cell surface.^26^

**Figure 4.**
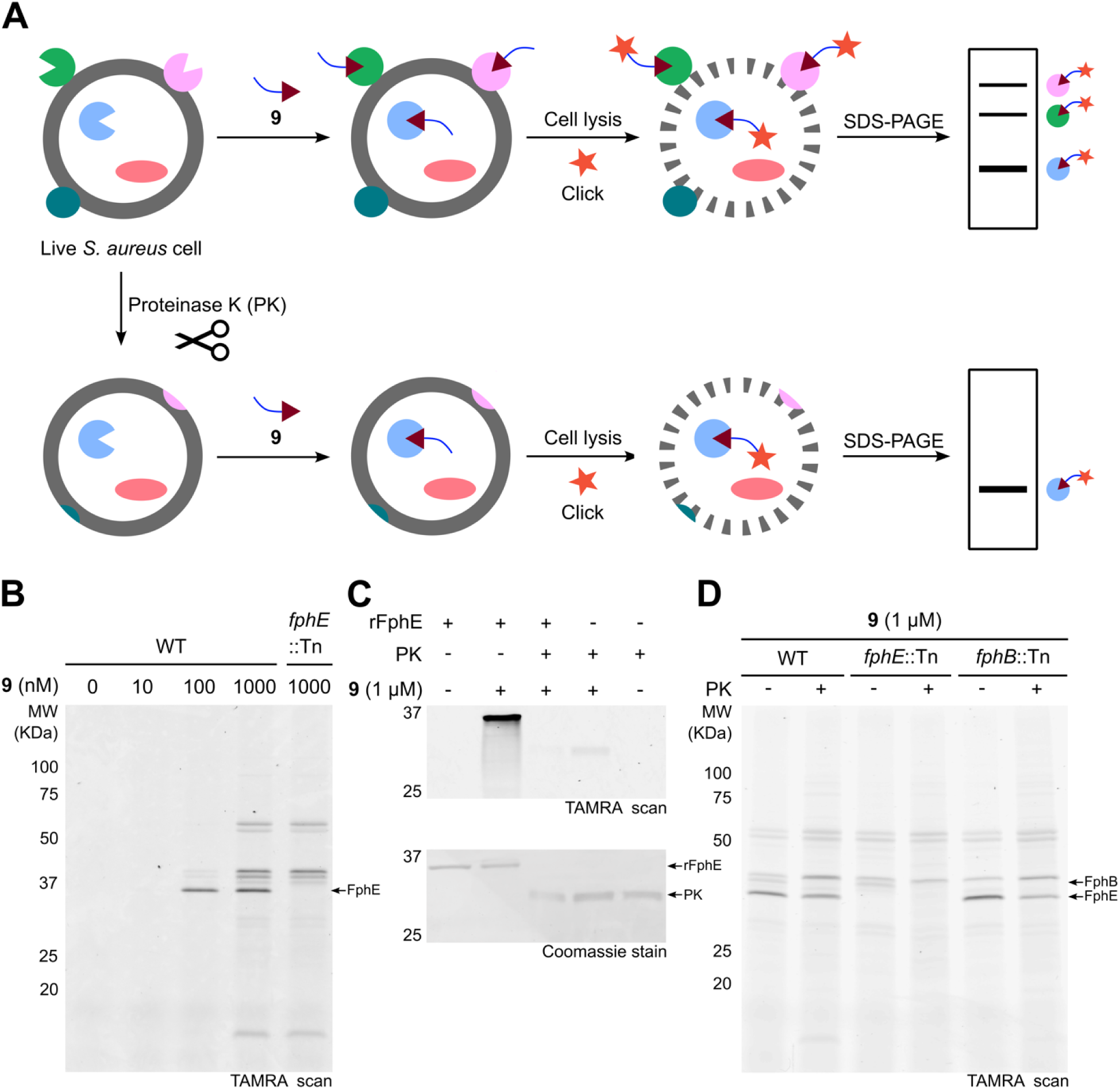
SDS-PAGE analysis of live *S. aureus* labeling and cellular localization analysis of FphE. (A) Schematic overview for assessment of localization of FphE with ABP **9**. Illustrated workflow of live *S. aureus* labeling to detect proteins covalently bound by **9** (top row) and protease protection assay to determine target protein localization (bottom row). (B) SDS-PAGE analysis of live *S. aureus* labeling with **9**. Wild-type (WT) and *fphE* transposon (Tn) mutant USA300 *S. aureus* strains were treated with the indicated concentrations of **9** at 37 °C for 2 h, then labeled with TAMRA-azide by CuAAC reaction. Afterwards, cells were lysed, and 10 µg of total proteins were loaded per lane. Proteins in cell lysates were quantified by BCA assay. Protein loading was assessed by Coomassie staining (Figure S1A). (C) Purified recombinant FphE (rFphE) was subjected to PK to verify that the enzyme is cleaved by the proteinase. (D) SDS-PAGE analysis of protease protection assay using WT, *fphE*::Tn, and *fphB*::Tn USA300 *S. aureus* strains. After 16h incubation with proteinase K (PK+) or without PK (PK−) at 37 °C, cells were treated with **9** at the same temperature for 2 h, then labeled with TAMRA-azide by CuAAC reaction. Total proteins (10 µg per lane) were used and protein loading was assessed by Coomassie staining (Figure S1B).

### JJ-OX-007 (**10**) as a selective imaging probe targeting FphE in live cells

We next synthesized JJ-OX-007 (**10**), a fluorescently labeled analog of ABP **9** in which the BODIPY dye is attached by click reaction prior to labeling (Figure 5A). This probe can be used for direct labeling and imaging studies without the need to do click chemistry. After confirming that probe **10** retained selective and covalent labeling of recombinant FphE (Figure S3 and Figure S4), we performed live cell labeling followed by gel-based ABPP (Figure 5B). The addition of the fluorophore in probe **10** increased the specificity of labeling of FphE compared to alkyne probe **9**. This result aligns with our previous observation that functionalizing the alkyne handle with BODIPY-TMR dye led to increased selectivity towards FphE.^64^

**Figure 5.**
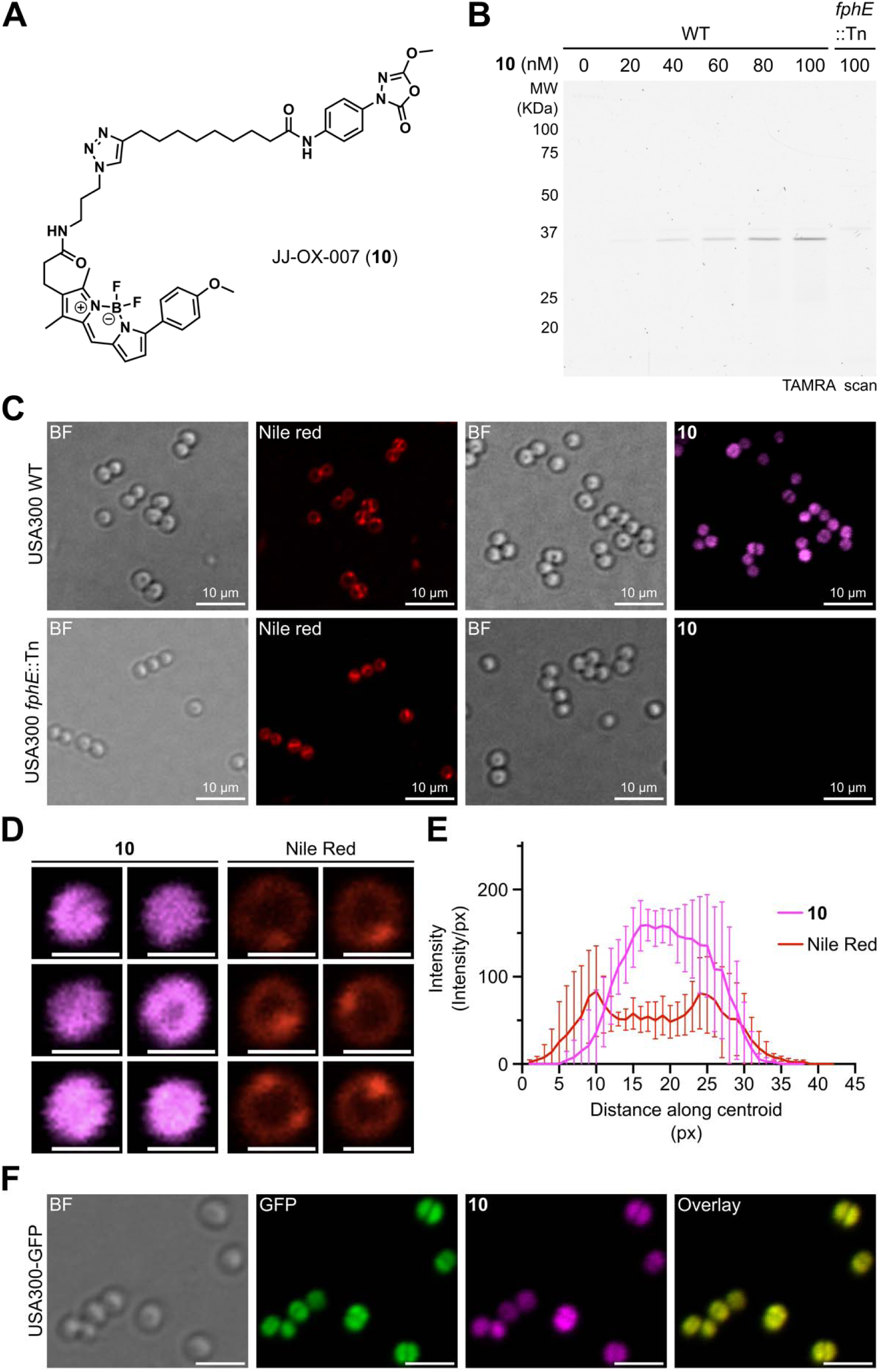
The probe JJ-OX-007 (**10**) selectively labels live *S. aureus* cells. (A) Chemical structure of fluorescent ABP **10**. Synthesis of this compound is described in the Supporting Information. (B) SDS-PAGE analysis of live *S. aureus* labeling with **10**. *S. aureus* USA300 wild-type (WT) and *fphE* transposon (*fphE*::Tn) mutant strains were treated with the indicated concentrations of **10** at 37 °C for 2 h. Cells were lysed, and 7 µg of total proteins were loaded per lane. Coomassie staining of the gel is shown in Figure S1C. (C) Confocal micrograph of *S. aureus* USA300 WT and *fphE*::Tn strains labelled with Nile Red (10 µg/mL) or **10 (**100 nM) during stationary phase. Fluorescence and brightfield (BF) images were obtained in the same measurement. (D) Representative single-cell confocal images of WT *S. aureus* USA300 that were treated by **10** (left) and Nile Red (right). Scale bar: 1 µm. (E) Quantified fluorescence intensity distribution across the cell. Microscopy images of cells were analyzed by implementing an agile script. The image was converted to binary format, cells were identified, and the major axis length along with the mean intensity of each cell was calculated. The mean intensity was plotted against the major axis length for each identified cell in the images. Bars represent mean ± standard deviation (*n*LJ=LJ6). (F) Confocal micrographs of *S. aureus* USA300-GFP cell labeled with **10** (100 nM) during stationary phase. Panel 1: BF; Panel 2: GFP; Panel 3: **10**; Panel 4: overlay of panels GFP and **10**. Scale bar: 5 μm.

Since FphE is a non-essential protein,^26, 42^ and probe **10** shows highly selective labeling of FphE, we anticipated it would not be toxic to *S. aureus*. This hypothesis was confirmed by an antimicrobial activity assay which revealed that probe **10** does not inhibit bacterial growth even at concentrations as high as 8 µg/mL (9.4 µM; Figure S5). This contrasts with the previously published oxadiazolone compound 3 which was shown to have potent MRSA killing activity.^42^ We believe this lack of toxicity of **10** is due to increased selectivity for FphE with loss of potency towards essential targets such as FabH. Because probe **10** is not cytotoxic, is suitable for use in live cell fluorescence imaging experiments. We first tested if the probe was selective enough to measure FphE activity in live cells using a plate reader-based assay. In fact, we were able to measure a statistically significant increase in the time-dependent fluorescence signal in the WT *S. aureus* cells compared to the *fphE* transposon mutant (*fphE*::Tn) using a standard plate reader (Figure S6).

We next assessed the applicability of **10** for imaging of FphE activity in live cells using confocal microscopy. We labeled WT and *fphE*::Tn *S. aureus* with the membrane dye Nile Red and probe **10** and then imaged the cells (Figure 5C). Since the two dyes have relatively close excitation wavelengths, we performed labeling studies using separate samples. In general, the fluorescent signal from Nile Red in the *fphE*::Tn strain exhibited a similar intensity and location to that of the wild-type cells (Figure 5C, second column). In contrast probe **10** produced a diffuse intracellular signal which was only observed in the WT *S. aureus* cells (Figure 5C, right most column). Interestingly, the labeling pattern of Nile Red in the WT *S. aureus* strain differs significantly from that of **10** (Figure 5C, upper panel), with the Nile Red signal enriched on the cell membrane and division septum and **10** showing broad and diffuse staining consistent with intracellular localization (Figure 5D and Figure 5E). To further confirm the subcellular localization of FphE, we used a strain of USA300 expressing an intracellular GFP reporter. The labeling pattern of probe **10** again showed central diffuse localization that overlapped with the GFP signal (Figure 5F).

### FphE as a target for specific imaging of S. aureus

Given the high degree of selectivity of probe **10** for FphE, which is a target that is only expressed in *S. aureus*, we wanted to determine if the probe could be used to selectively label *S. aureus* cells in the context of an *in vivo* infection. We first determined if probe **10** could specifically label *S. aureus* in the presence of other microbial species including pathogenic Gram-positive and Gram-negative bacteria, as well as the representative pathogenic yeast *Candida albicans*. Labeling of each of the microbes with probe **10** and the general serine protease probe FP-TMR followed by analysis of total cell lysates by SDS-PAGE (Figure 6A) confirmed that while multiple serine hydrolases were labeled by the FP-TMR probe in all tested microorganisms (Figure 6A, left gel), probe **10** predominantly labeled FphE in *S. aureus* (Figure 6A, right gel). To further confirm the specificity of probe **10** in live cells, we directly labeled a mixed culture of the microbes used for the gel labeling studies (including a GFP-expressing USA300 *S. aureus* strain in place of the corresponding unlabeled strain) and then performed confocal microscopy. This analysis revealed that within the mixed population of bacteria, only *S. aureus* cells were selectively labeled by probe **10** as confirmed by its colocalization only with the GFP positive *S. aureus* cells (Figure 6B). These results suggest that our probe should enable specific detection of *S. aureus* infections, with discrimination over most other common human pathogenic bacteria and yeast.

**Figure 6.**
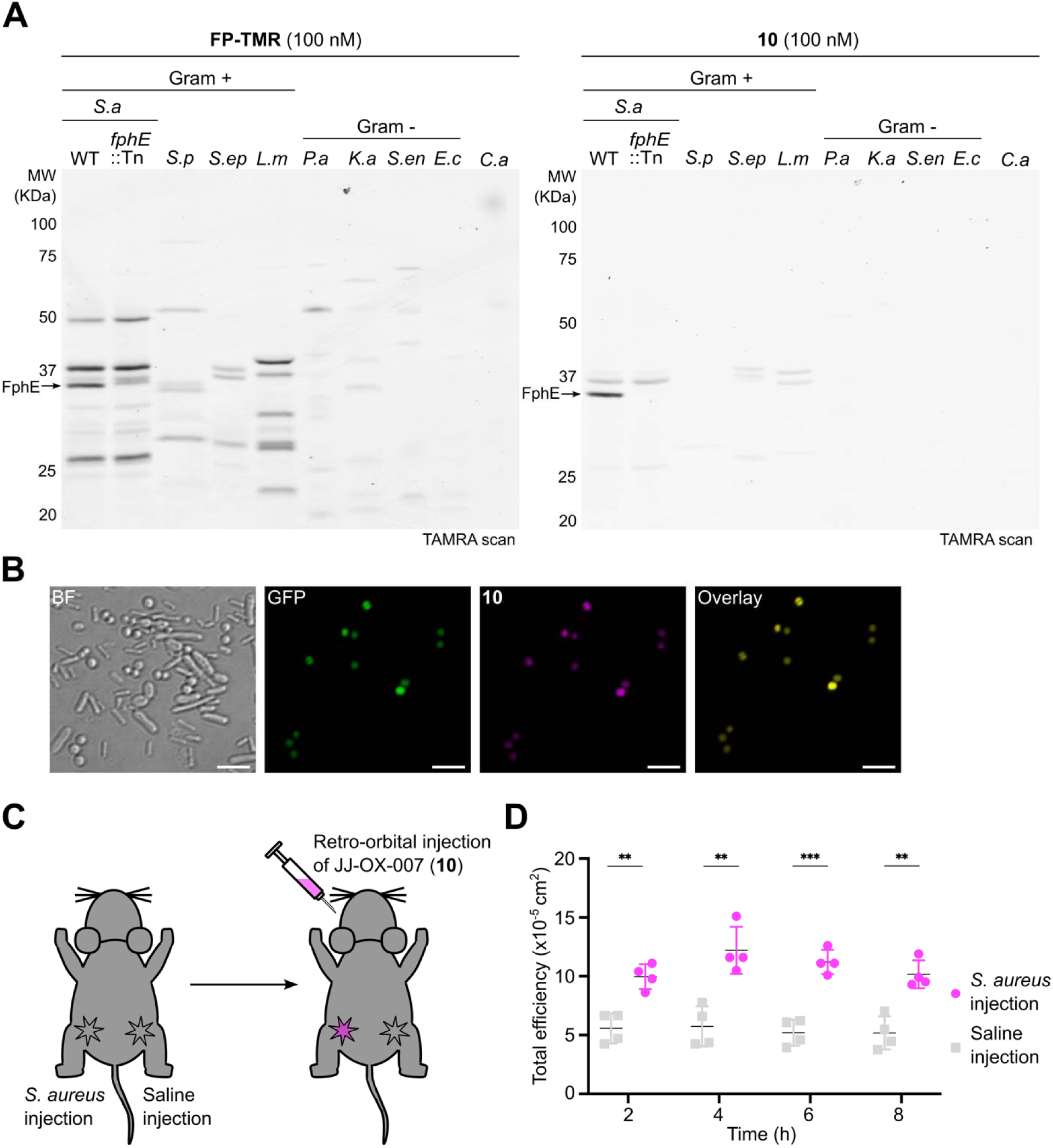
FphE as a target for *S. aureus* imaging. (A) SDS-PAGE analysis of indicated Gram-positive and Gram-negative bacterial pathogens labeled with fluorophosphonate tetramethylrhodamine (FP-TMR; left gel) or JJ-OX-007 (**10**; right gel). *S. aureus* (*S*.*a*) USA300 WT and transposon mutant (*fphE*::Tn), *S.p*: *Streptococcus pyogenes*, *S.ep*: *S. epidermidis*, *L.m*: *Listeria monocytogenes*, *P.a*: *Pseudomonas aeruginosa*, *K.a*: *Klebsiella aerogenes*, *S.en*: *Salmonella enterica*, *E.c*: *E. coli*, *C.a*: *Candida albicans*. Total proteins (5 µg per lane) were used, and protein loading was assessed by Coomassie staining (Figure S1D). (B) Confocal micrographs of mixed samples of *S. aureus* USA300-GFP with the listed Gram-positive and Gram-negative bacterial pathogens labeled by **10**. Scale bar: 10 μm. (C) Schematic representation of imaging experiment using probe **10** to visualize *S. aureus* infection site in mice. (D) Total efficiency of fluorescence detected in the infected thigh (magenta) or un-infected (saline injected) thigh (gray) over time in hours post-injection of **10** (*n* = 4). Probe was injected retro-orbitally 2 hours after the inoculation. Bars represent mean ± standard deviation. *P* values were determined using unpaired two-tailed Student’s *t*-tests. \*\**P* < 0.01, \*\*\**P* < 0.001.

To confirm that our probe has the potential to be used in human imaging studies, we tested for possible off targets of the probe in human cells. We therefore conducted labeling studies in intact and cell lysates derived from human embryonic kidney 293T (HEK293T) cell (Figure S7). Only one protein in the HEK293T lysate proteome was labeled by **10**, however, this labeling was dramatically reduced in the presence of recombinant FphE (Figure S7A). Furthermore, we observed no off target labeling when probe **10** was used with intact HEK293T cells (Figure S7B). Taken together, these results suggest that **10** is highly selective for FphE over other possible off targets in human cells.

Finally, we wanted to determine if probe **10** could be used to image *S. aureus* infections *in vivo* using optical imaging methods. We used a model of infection where mice were subcutaneously injected with *S. aureus* in the left hind thigh and normal saline was administered into the opposite thigh. We injected probe **10** retro-orbitally 1.5 hours after initial infection and then imaged the live mice using a fluorescence imaging system (Figure 6C). These results showed selective localization of probe **10** only at the infection sites compared to the saline control (Figure S8). Quantification of the total fluorescence (Figure 6D) and its ratio between the infected site and the non-infected site (Table S5) showed that probe **10** exhibited significantly higher signal at the injection site with the highest fluorescence ratio of 2.31-fold at 4 hours post-injection. This contrast over the saline control remained relatively consistent at approximately 2-fold from 2 to 8 hours, demonstrating the potential value of this probe for *S. aureus*-specific imaging.

## Conclusion

In summary, our study has successfully characterized FphE as a highly specific serine hydrolase expressed in *S. aureus*, setting it apart from other pathogenic microorganisms or human cells. Our rational design of FphE-targeting ABPs led to the development of JJ-OX-004 (**9**), an oxadiazolone-based irreversible covalent alkyne probe of FphE. Further structural analysis confirmed the binding mechanism of oxadiazolone warhead and the covalent modification of FphE by **9**. This probe also provided evidence through ABPP combined with a protease protection assay that FphE is an intracellular serine hydrolase. The subsequent development of the fluorescent probe JJ-OX-007 (**10**) enhanced selectivity for FphE in live *S. aureus* cells, providing a potential new imaging probe for *S. aureus*-specific detection. Our *in vivo* studies demonstrated the rapid and selective localization of probe **10** at the sites of *S. aureus* infections. Overall, our findings suggest that the oxidazolone electrophile can be chemically tuned to generate covalent probes that specifically target serine hydrolases *in vivo*. Taken together, these results suggest that FphE is a promising target for the development of *S. aureus*-specific imaging probes, and probe **10** has great potential for clinical applications for both early diagnosis and subsequent therapy monitoring.

## Supporting information

Blast results

PDB deposition validation 8T87

PDB deposition validation 8T88

## Associated content

### Supporting Information

Supplementary tables and figures, materials and methods for biological evaluation, detailed synthesis procedures and compound characterization, LC-MS, and NMR spectra (PDF)

Fph BLASTp results (XLSX)

### Accession Codes

The datasets generated and analyzed in this study are available in the worldwide Protein Data Bank (PDB) under PDB ID 8T87 (FphE unbound) and 8T88 (FphE bound with **9**). The authors will release the atomic coordinates and experimental data upon article publication.

## Acknowledgements

The authors thank Dr. Laura J. Keller at Genentech for helpful discussions. We thank Dr. Nathaniel I. Martin, Dr. Mario van der Stelt, and members of their laboratories at Leiden University for sharing compound 3. We also would like to thank Deborah Yung and Dr. Daniel Pletzer at the University of Otago for help obtaining the *S. aureus* USA300 genomic template. This work was funded by National Institutes of Health grants R01-EB026332 (to M.B.) and by the Stanford Clinician Scientist Training Program with the training grant R25DC020174 (to K.W.P). We gratefully acknowledge the funding support by the program “Excellence initiative— research university” for the AGH University in Krakow as well as the ARTIQ project ARTIQ/0004/2021 (to J.J.-K.). This work was supported in part by the New Zealand Marsden Fund Council from government funding, managed by Royal Society Te Apārangi and a Sir Charles Hercus Health Research Fellowship (to M.F.). This research was undertaken in part using the MX1 and MX2 beamlines at the Australian Synchrotron, part of ANSTO, and made use of the ACRF detector.

