## Supporting Information for "Development of Oxadiazolone Activity-Based Probes Targeting FphE for Specific Detection of *S. aureus* Infections"

|  | **Table of Contents** | **Page** |
| --- | --- | --- |
| **1.** | **Supplementary Tables and Figures** | **S4** |
|  | **Table S1**: The homology percentage data used for Figure 1A | S4 |
|  | **Table S2**: FphE unbound and ligand data collection and processing | S5 |
|  | **Table S3**: FphE unbound and ligand structure solution and refinement | S6 |
|  | **Table S4:** List of microbial strains used in this study | S7 |
|  | **Table S5:** Quantification of fluorescence signals in Figure S10 | S8 |
|  | **Figure S1**: Coomassie staining of SDS-PAGE gels shown in Figure 4B, Figure 4D, Figure 5B, and Figure 6A. | S9 |
|  | **Figure S2**: Comparison of bound and unbound FphE | S10 |
|  | **Figure S3**: Concentration-dependent labeling of WT and S103A FphE by JJ-OX-007 (**10**) | S11 |
|  | **Figure S4**: FphE - **10** crystallographic investigation | S12 |
|  | **Figure S5**: Growth inhibition activity test of **10** on *S. aureus* USA300 | S13 |
|  | **Figure S6**: Time-dependent *S. aureus* labeling with **10** | S14 |
|  | **Figure S7**: SDS-PAGE analysis of HEK293T lysate or intact cell labelling with **10** | S15 |
|  | **Figure S8**: *In vivo* fluorescent imaging of mouse subcutaneous wound infection model | S16 |
|  | **Figure S9**: Uncropped gels | S17 |
|  | **Figure S10**: Uncropped microscopy images | S18 |
| **2.** | **Materials and Methods** | **S19** |
|  | FphE cloning | S19 |
|  | Protein expression and purification | S19 |
|  | Substrate screening and enzyme inhibition assay | S20 |
|  | Gel labeling analysis of purified recombinant proteins | S21 |
|  | Mass spectrometry validation of FphE labeling with JJ-OX-004 (**9**) | S21 |
|  | Jump dilution assay | S22 |
|  | FphE crystallization | S22 |
|  | FphE data collection, processing, refinement, deposition, and analysis | S22 |
|  | Bacterial culture and bacterial growth inhibition assay | S23 |
|  | SDS-PAGE analysis of live *S. aureus* labeling | S23 |
|  | Protease protection assay | S24 |
|  | Plate-reader based live *S. aureus* labeling assay | S24 |
|  | Confocal microscopy | S25 |
|  | Labeling of different bacterial strains | S25 |
|  | Quantitative analysis of microscopy images | S26 |
|  | Mouse subcutaneous infection model | S27 |
|  | HEK293T cell labeling | S27 |
| **3.** | **Chemical Synthesis** | **S29** |
|  | General methods | S29 |
|  | **Scheme S1**: Synthesis of JJ-OX-001 (**6**) | S30 |
|  | **Scheme S2**: Synthesis of JJ-OX-002 (**7**) and JJ-OX-003 (**8**) | S30 |
|  | **Scheme S3**: Synthesis of JJ-OX-004 (**9**) and JJ-OX-007 (**10**) | S31 |
|  | Synthesis of JJ-OX-001 (**6**) | S32 |
|  | Synthesis of JJ-OX-002 (**7**) and JJ-OX-003 (**8**) | S34 |
|  | Synthesis of JJ-OX-004 (**9**) and JJ-OX-007 (**10**) | S37 |
| **4.** | **LC traces of synthesized probes 7–10** | **S40** |
| **5.** | **^1^H NMR Spectra** | **S44** |
| **6.** | **^13^C NMR Spectra** | **S50** |
| **7.** | **References** | **S56** |

**1. Supplementary Tables and Figures**

**Table S1.** The homology percentage data used for Figure 1A

All BLASTp results for Fph proteins are provided in a separate Excel file.

|  | FphA | FphB | FphC | FphD | FphE | FphF | FphG | FphH | FphI | FphJ |
| --- | --- | --- | --- | --- | --- | --- | --- | --- | --- | --- |
| *Staphylococcus aureus* | 100 | 100 | 100 | 100 | 100 | 100 | 100 | 100 | 100 | 100 |
| *Staphylococcus epidermidis* | 51 | 57.9 | 51.6 | 56.1 | 0 | 74.1 | 71.8 | 85 | 51.5 | 0 |
| *Listeria monocytogenes* | 0 | 0 | 33.1 | 34.7 | 67.6 | 43.7 | 0 | 52 | 32.7 | 42.8 |
| *Clostridioides difficile* | 0 | 0 | 30.2 | 0 | 0 | 34.7 | 0 | 0 | 0 | 0 |
| *Streptococcus pneumoniae* | 0 | 0 | 0 | 0 | 0 | 30.2 | 0 | 0 | 0 | 0 |
| *Streptococcus pyogenes* | 0 | 33.3 | 0 | 0 | 0 | 31.7 | 0 | 0 | 0 | 0 |
| *Streptococcus agalactiae* | 0 | 33.7 | 0 | 24.6 | 0 | 32.1 | 0 | 0 | 0 | 0 |
| *Enterococcus faecalis* | 0 | 25.2 | 32.6 | 35.5 | 0 | 35.1 | 0 | 35.6 | 23.6 | 54.9 |
| *Enterococcus faecium* | 0 | 0 | 0 | 0 | 0 | 0 | 0 | 0 | 0 | 0 |
| *Pseudomonas aeruginosa* | 0 | 30.1 | 29.9 | 0 | 0 | 0 | 0 | 0 | 0 | 0 |
| *Klebsiella pneumoniae* | 27.7 | 0 | 0 | 0 | 0 | 0 | 0 | 0 | 0 | 0 |
| *Campylobacter jejuni* | 0 | 0 | 0 | 0 | 0 | 0 | 0 | 0 | 0 | 0 |
| *Chlamydia trachomatis* | 0 | 0 | 0 | 0 | 0 | 0 | 0 | 0 | 0 | 0 |
| *Neisseria meningitidis* | 0 | 0 | 0 | 0 | 0 | 0 | 0 | 0 | 0 | 0 |
| *Neisseria gonorrhoeae* | 0 | 0 | 0 | 0 | 0 | 0 | 0 | 0 | 0 | 0 |
| *Acinetobacter baumannii* | 0 | 0 | 0 | 0 | 0 | 0 | 0 | 0 | 0 | 0 |
| *Legionella pneumophila* | 0 | 0 | 0 | 0 | 0 | 0 | 0 | 0 | 0 | 0 |
| *Aeromonas hydrophila* | 0 | 0 | 0 | 0 | 0 | 0 | 0 | 0 | 0 | 0 |
| *Vibrio cholerae* | 0 | 0 | 0 | 0 | 0 | 0 | 0 | 0 | 0 | 0 |
| *Haemophilus influenzae* | 0 | 0 | 0 | 0 | 0 | 0 | 0 | 0 | 0 | 0 |
| *Proteus mirabilis* | 0 | 0 | 0 | 0 | 0 | 0 | 0 | 0 | 0 | 0 |
| *Providencia stuartii* | 0 | 0 | 0 | 0 | 0 | 0 | 0 | 0 | 0 | 0 |
| *Morganella morganii* | 0 | 0 | 0 | 0 | 0 | 0 | 0 | 0 | 0 | 0 |
| *Salmonella enterica* | 31.2 | 0 | 0 | 0 | 0 | 0 | 0 | 0 | 0 | 0 |
| *Escherichia coli* | 0 | 0 | 0 | 0 | 0 | 0 | 0 | 0 | 0 | 0 |
| *Shigella sonnei* | 0 | 0 | 0 | 0 | 0 | 0 | 0 | 0 | 0 | 0 |
| *Serratia marcescens* | 0 | 0 | 0 | 0 | 0 | 0 | 0 | 0 | 0 | 0 |
| *Klebsiella aerogenes* | 0 | 0 | 0 | 0 | 0 | 0 | 0 | 0 | 0 | 0 |
| *Mycoplasma pneumoniae* | 0 | 0 | 0 | 0 | 0 | 0 | 0 | 0 | 0 | 0 |
| *Candida albicans* | 0 | 0 | 0 | 0 | 0 | 0 | 0 | 0 | 0 | 0 |
| *Homo sapiens* | 36.3 | 0 | 0 | 0 | 0 | 0 | 0 | 0 | 0 | 0 |

**Table S2.** FphE unbound and ligand-bound data collection and processing

Values for the outer shell are given in parentheses.

|  | Unbound | **9** (JJ-OX-004) bound |
| --- | --- | --- |
| PDB ID | 8T87 | 8T88 |
| Diffraction source | Australian synchrotron MX2 | Australian synchrotron MX1 |
| Wavelength (Å) | 0.954 | 0.954 |
| Detector | DECTRIS EIGER X 16M | DECTRIS EIGER X 9M |
| Space group | P 1 2_1_ 1 | P 1 2_1_ 1 |
| a, b, c (Å) | 46.8 74.1 71.4 | 46.7 74.3 73.6 |
| α, β, γ (°) | 90.0, 91.3, 90.0 | 90.0, 91.7, 90.0 |
| Resolution range (Å) | 46.78 – 1.62  (1.65 – 1.62) | 39.99 – 1.54  (1.56 – 1.54) |
| Total No. of reflections | 359349 (15655) | 390181 (18405) |
| No. of unique reflections | 60409 (2673) | 73321 (3543) |
| Completeness (%) | 98.4 (88.6) | 98.0 (95.1) |
| Redundancy | 5.9 (5.9) | 5.3 (5.2) |
| 〈 *I*/σ(*I*)〉 | 12.9 (1.4) | 17.5 (1.3) |
| CC_1/2_ | 0.999 (0.597) | 1.000 (0.598) |
| *R*_merge._ | 0.054 (1.048) | 0.033 (0.938) |
| *R*_p.i.m._ | 0.036 (0.706) | 0.024 (0.689) |

**Table S3.** FphE unbound and ligand-bound structure solution and refinement

Values for the outer shell are given in parentheses.

|  | Unbound | **9** (JJ-OX-004) bound |
| --- | --- | --- |
| PDB ID | 8G48 | 8G49 |
| Resolution range (Å) | 38.73 – 1.62 (1.65 – 1.62) | 39.99 – 1.54 (1.56 – 1.54) |
| Final R-work | 0.187 (0.390) | 0.157 (0.307) |
| Final R-free | 0.219 (0.432) | 0.186 (0.316) |
| Protein residues | 554 | 554 |
| Ligands JJ004 | 0 | 2 |
| Magnesium atoms | 6 | 2 |
| Water molecules | 219 | 456 |
| R.m.s. deviations |  |  |
| Bonds (Å) | 0.011 | 0.010 |
| Angles (°) | 1.135 | 1.077 |
| Average *B* factors (Å^2^) | 44.3 | 31.5 |
| Ligands | - | 46.8 |
| Magnesium atoms | 41.2 | 24.8 |
| Water molecules | 39.6 | 34.6 |
| Ramachandran plot |  |  |
| Most favored (%) | 98.4 | 98.6 |
| Outlier (%) | 0 | 0 |

**Table S4.** List of microbial strains used in this study

| **Strain** | **Description** | **Reference/Source** |
| --- | --- | --- |
| *S. aureus* USA 300 | Wild-type USA300 Los Angeles County  (LAC) clone; multilocus sequence type  8, SCCmec type IV cured of antibiotic  resistance plasmid | ^1^ |
| *S. aureus* USA 300 *fphE*::Tn | Transposon insertion mutant in  SAUSA300_2518; Ery^R^, Linc^R^ | Nebraska Transposon Mutant  Library |
| *S. aureus* USA 300 *fphB*::Tn | Transposon insertion mutant in  SAUSA300_2473; Ery^R^, Linc^R^ | Nebraska Transposon Mutant  Library |
| *S. aureus* USA 300 LAC + pCM29 (GFP) | Wild-type USA300 LAC strain with GFP plasmid pCM29; Cm^R^ | Manuel Amieva, Stanford University |
| *Staphylococcus epidermidis* ATCC 12228 | Wild-type strain | Elizabeth Joyce, UCSF |
| *Streptococcus pyogenes* ATCC 12384 | Wild-type strain | Niaz Banaei, Stanford University |
| *Listeria monocytogenes* | Wild-type strain | Michael Howitt, Stanford University |
| *Pseudomonas aeruginosa* ATCC 27853 | Wild-type strain | Niaz Banaei, Stanford University |
| *Klebsiella aerogenes* ATCC 13048 | Wild-type strain | Niaz Banaei, Stanford University |
| *Salmonella enterica* LT2 | Wild-type strain | Denise Monack, Stanford University |
| *Escherichia coli* BW25113 | Wild-type strain | Coli Genetic Stock Center |
| *Candida albicans* ATCC 10231 | Wild-type strain | Niaz Banaei, Stanford University |

**Table S5.** Quantification of fluorescence signals in **Figure S10**

Values are mean ± standard deviation (*n* = 4).

| Time (h) | Total efficiency^a^  (infected, ×10^-5^ cm^2^) | Total efficiency  (non-infected, ×10^-5^ cm^2^) | Total efficiency ratio^b^  (infected/non-infected) |
| --- | --- | --- | --- |
| 2 | 9.97 ± 1.06 | 5.57 ± 1.27 | 1.87 ± 0.50 |
| 4 | 12.20 ± 2.00 | 5.74 ± 1.72 | 2.31 ± 0.91 |
| 6 | 11.23 ± 1.03 | 5.21 ± 1.12 | 2.26 ± 0.71 |
| 8 | 10.16 ± 1.19 | 5.18 ± 1.42 | 2.11 ± 0.79 |

^a^Mean signal efficiency × area of the ROI. Signal efficiency is defined as a ratio of detected emission radiance over LED excitation radiance.

^b^Values are presented as the mean ± standard deviation of the total efficiency ratio for each mouse.

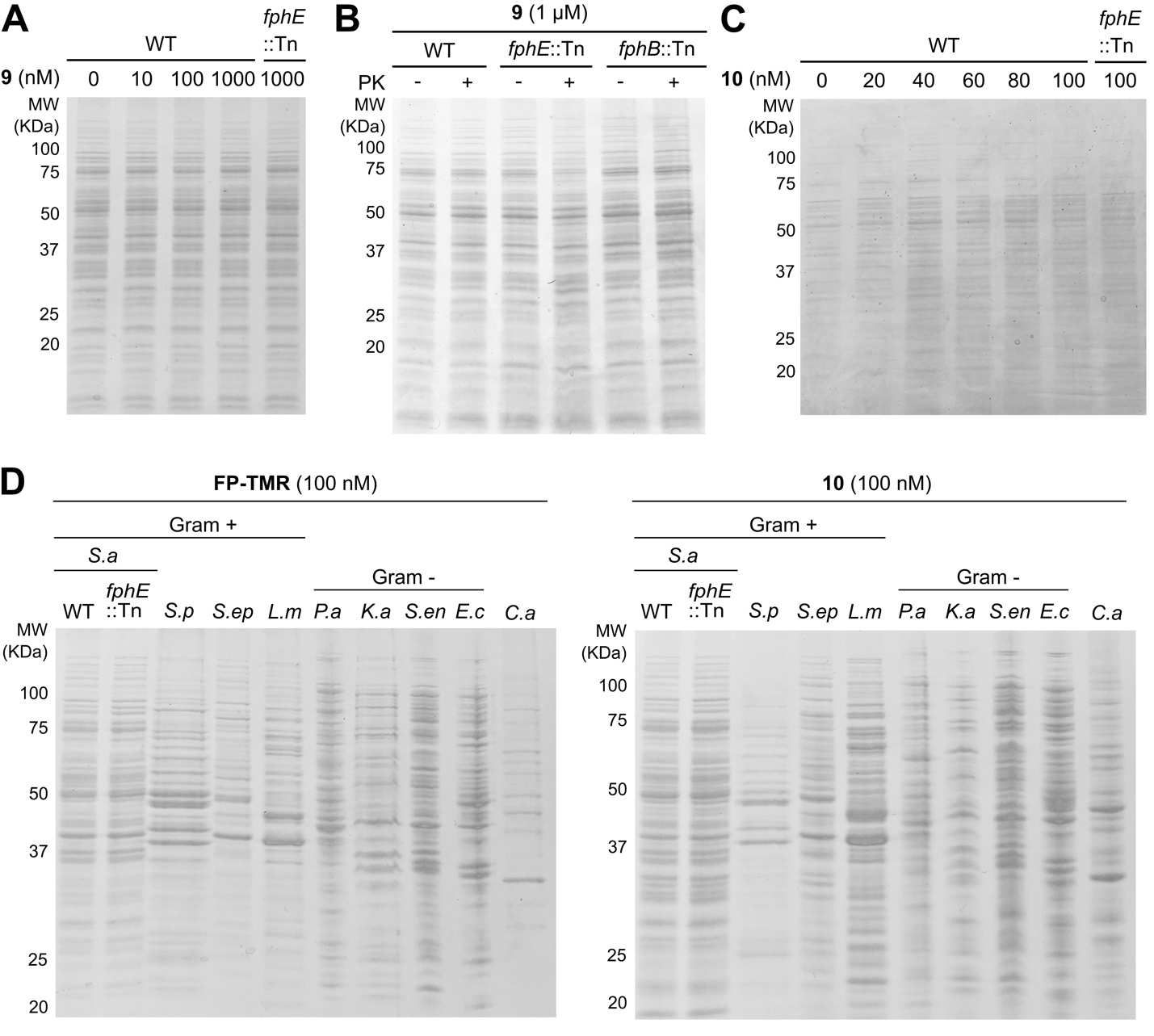
**Figure S1.** Coomassie staining of SDS-PAGE gels shown in (A) Figure 4B, (B) Figure 4D, (C) Figure 5B, and (D) Figure 6A.

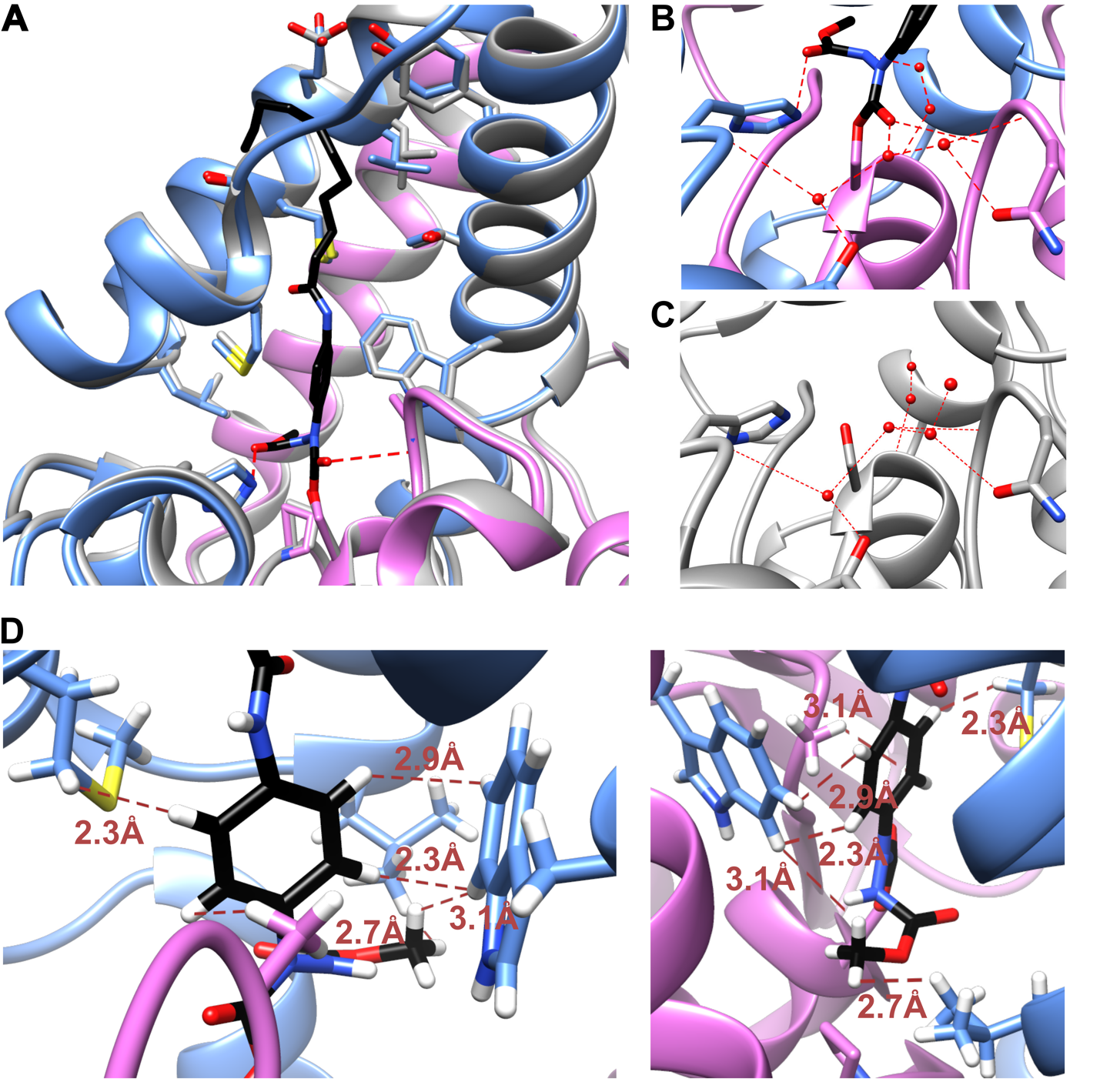
**Figure S2.** Comparison of bound and unbound FphE. (A) Cα alignment of **9** bound (PDB ID 8T88; dimer copies in magenta and blue) with unbound structure (8T89; all in grey). (B) Hydrogen bonding network in **9** bound structure. (C) Hydrogen bonding network in unbound structure. (D) Hydrophobic network around the ring of **9**, two different orientations.

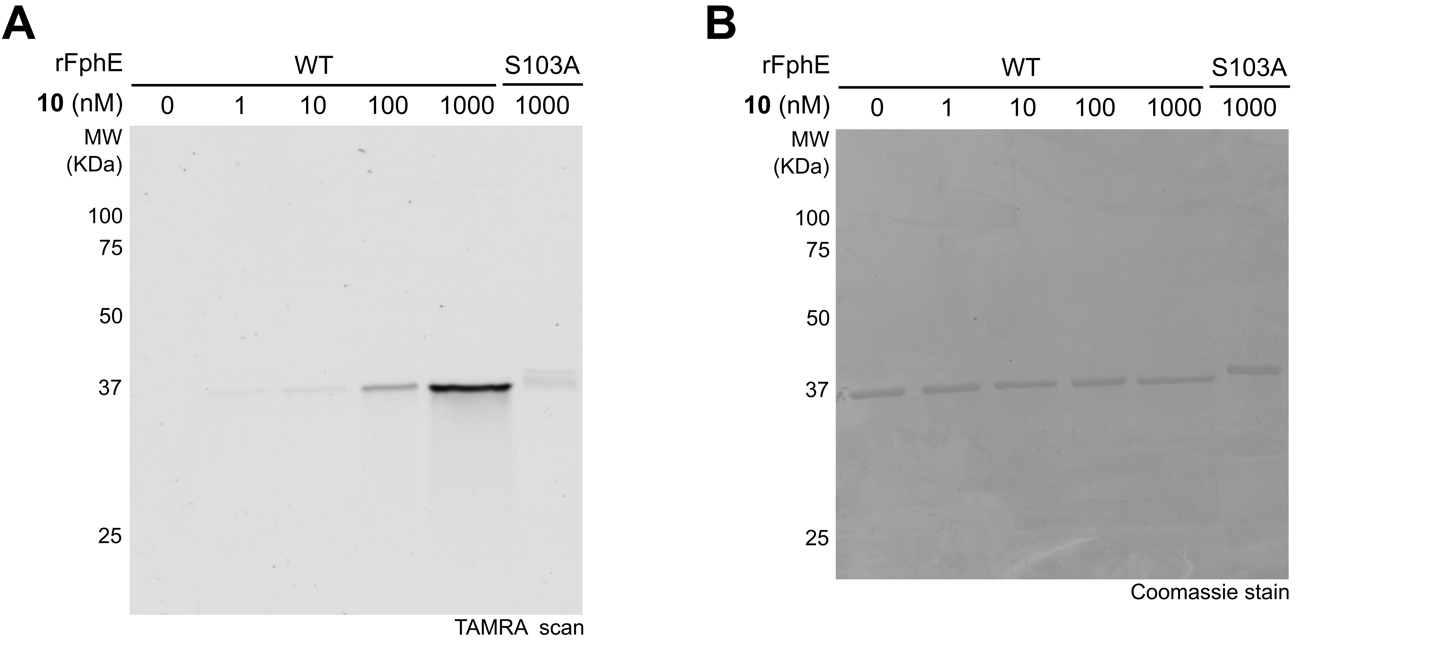
**Figure S3.** Concentration-dependent labeling of WT and S103A FphE by JJ-OX-007 (**10**). FphE were incubated with the probe for 1 h and then analyzed by (A) SDS-PAGE/fluorescence scan. (B) Protein loading was assessed by Coomassie staining.

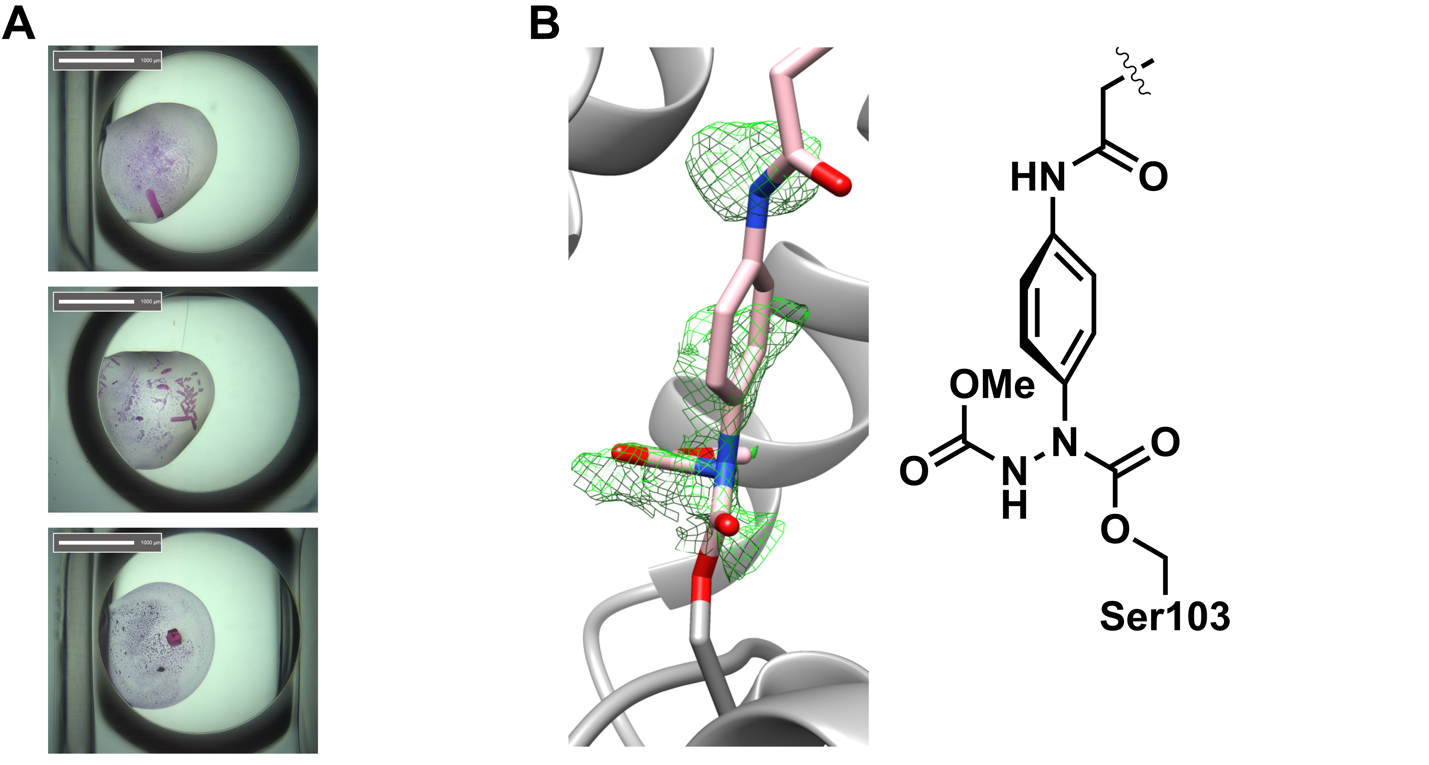
**Figure S4.** FphE -**10** crystallographic investigation. (A) Examples of probe accumulation in FphE crystals turning them pink. (B) Best electron density evidence of **10** binding to FphE crystals (left). The *F_o_−F_c_* electron density map for the ligand region is shown after a first round of refinement without placing a ligand in refinement. The model shown is the FphE-**9** bound structure (8T88) overlaid on the dataset in the presence of **10**. A simplified representation of the covalently modified chemical structure is presented (right).

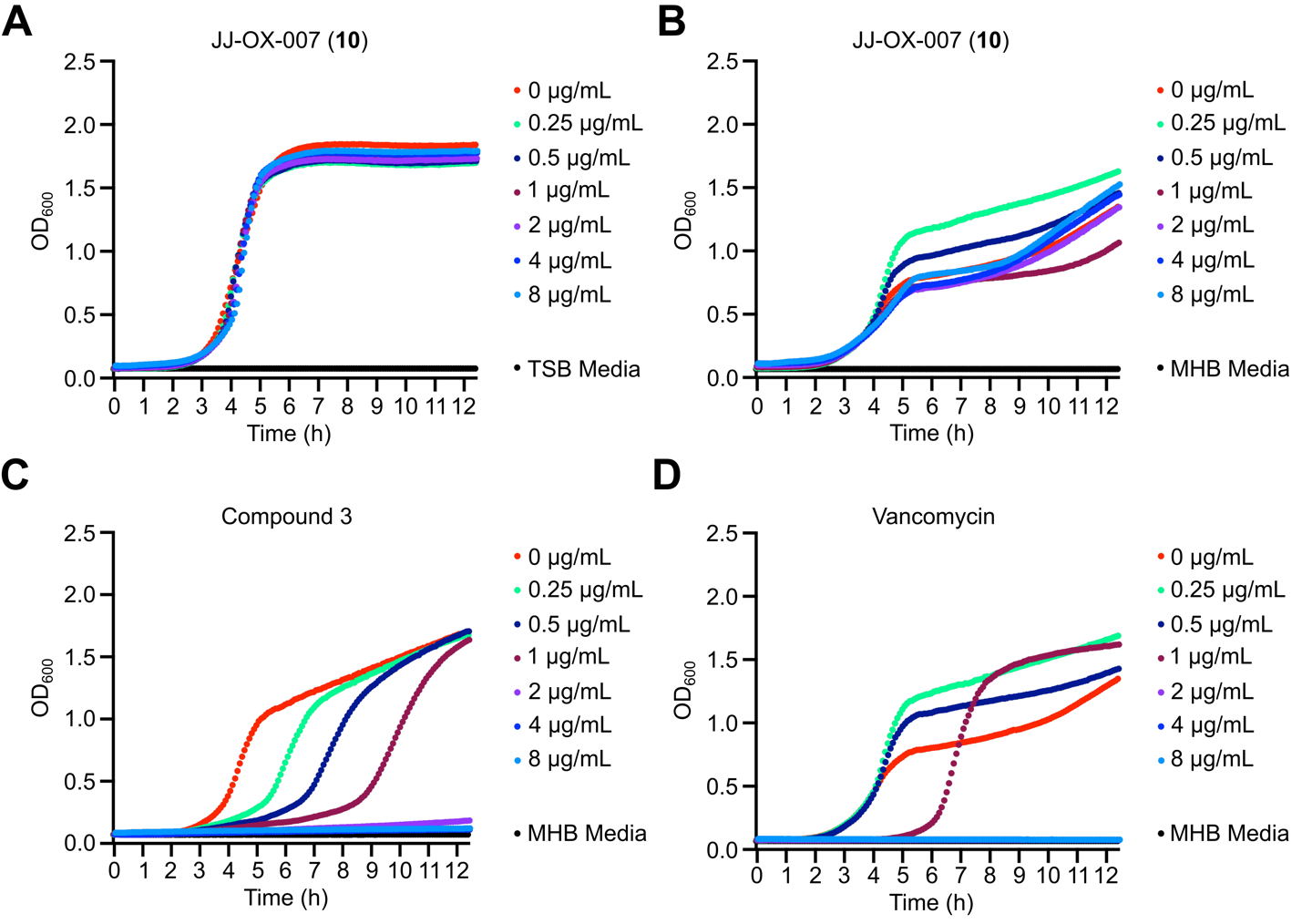
**Figure S5.** Growth inhibition assay of **10** on *S. aureus* USA300. Bacterial growth curves in the presence of **10** under at various concentrations in (A) tryptic soy broth (TSB) media and (B) Mueller-Hinton broth (MHB) media. Bacterial growth curves under various concentrations of (C) compound 3^2^ and (D) Vancomycin in MHB media.

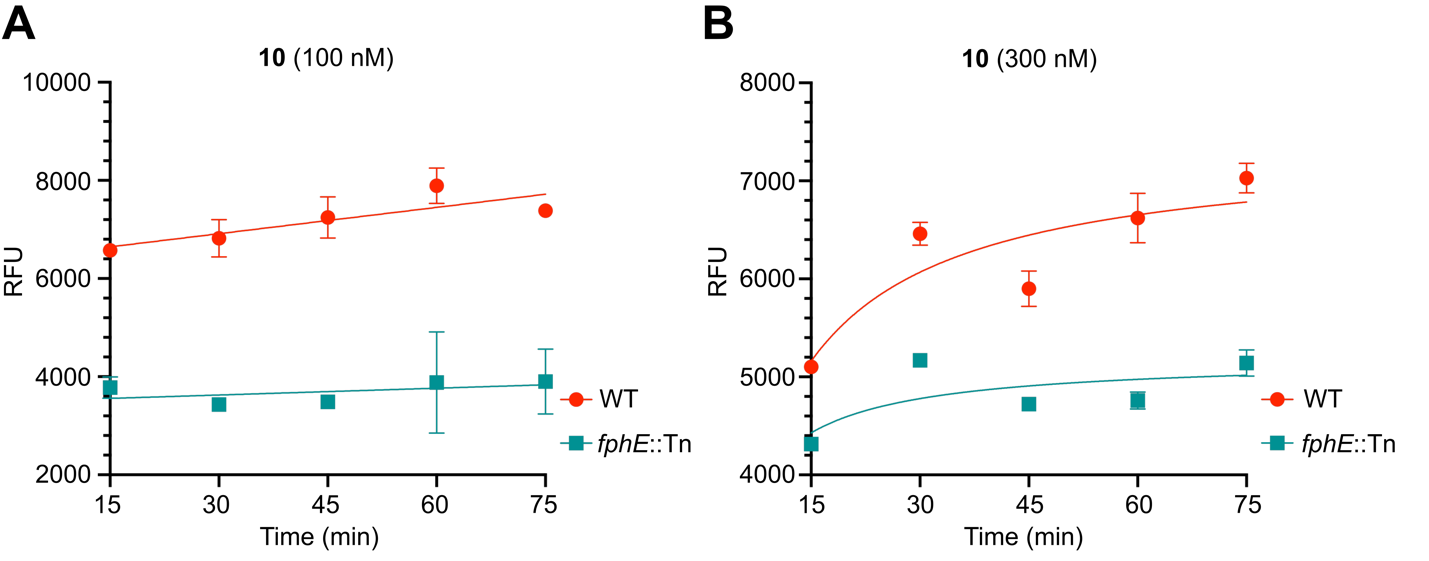
**Figure S6.** Time-dependent *S. aureus* labeling with **10**. *S. aureus* USA300 wild-type (WT) and *fphE* transposon mutant (*fphE*::Tn) strains were treated with (A) 100 nM or (B) 300 nM of **10** at 37 °C. Labeling of intact cells were monitored every 15 minutes for each strain in 384-well plate after washing cells three times with 1X PBS. Fluorescence was detected using 552 nm excitation and 578 nm emission filter. Bars represent mean ± standard deviation (*n* = 3).

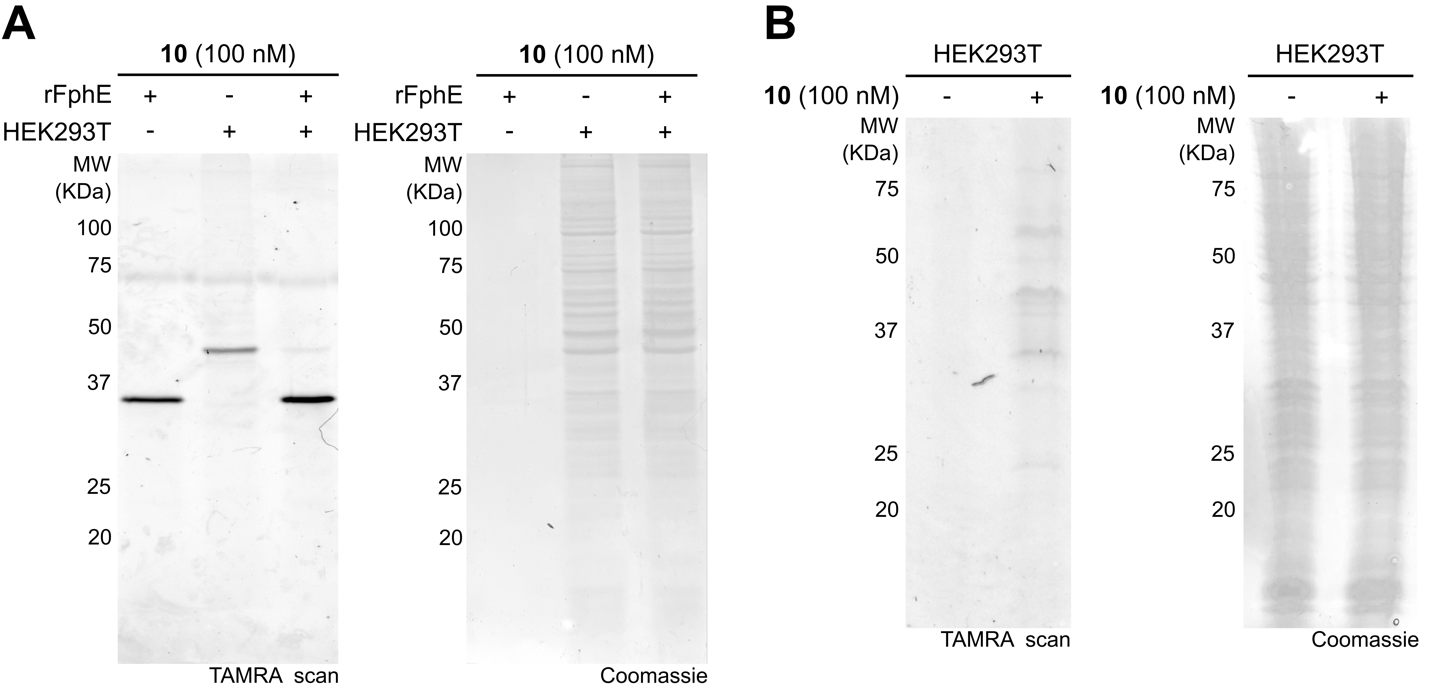
**Figure S7.** SDS-PAGE analysis of HEK293T lysate or intact cell labelling with **10**. (A) Representative in-gel HEK293T lysate labeling with **10** (Left). HEK293T cell lyaste (WT) with or without recombinant FphE (rFphE, 30 ng) were treated with the indicated concentrations of **10** at 37 °C for 2 h. After then, 7 µg of total proteins were loaded per lane. Coomassie staining of the gel is shown (right). (B) Labeling profile of intact HEK293T cell. After 2 h preincubation with **10**, cells were lysed and analyzed by SDS-PAGE (left). 7 µg of total proteins were loaded per lane, and protein loading was assessed by Coomassie staining (right).

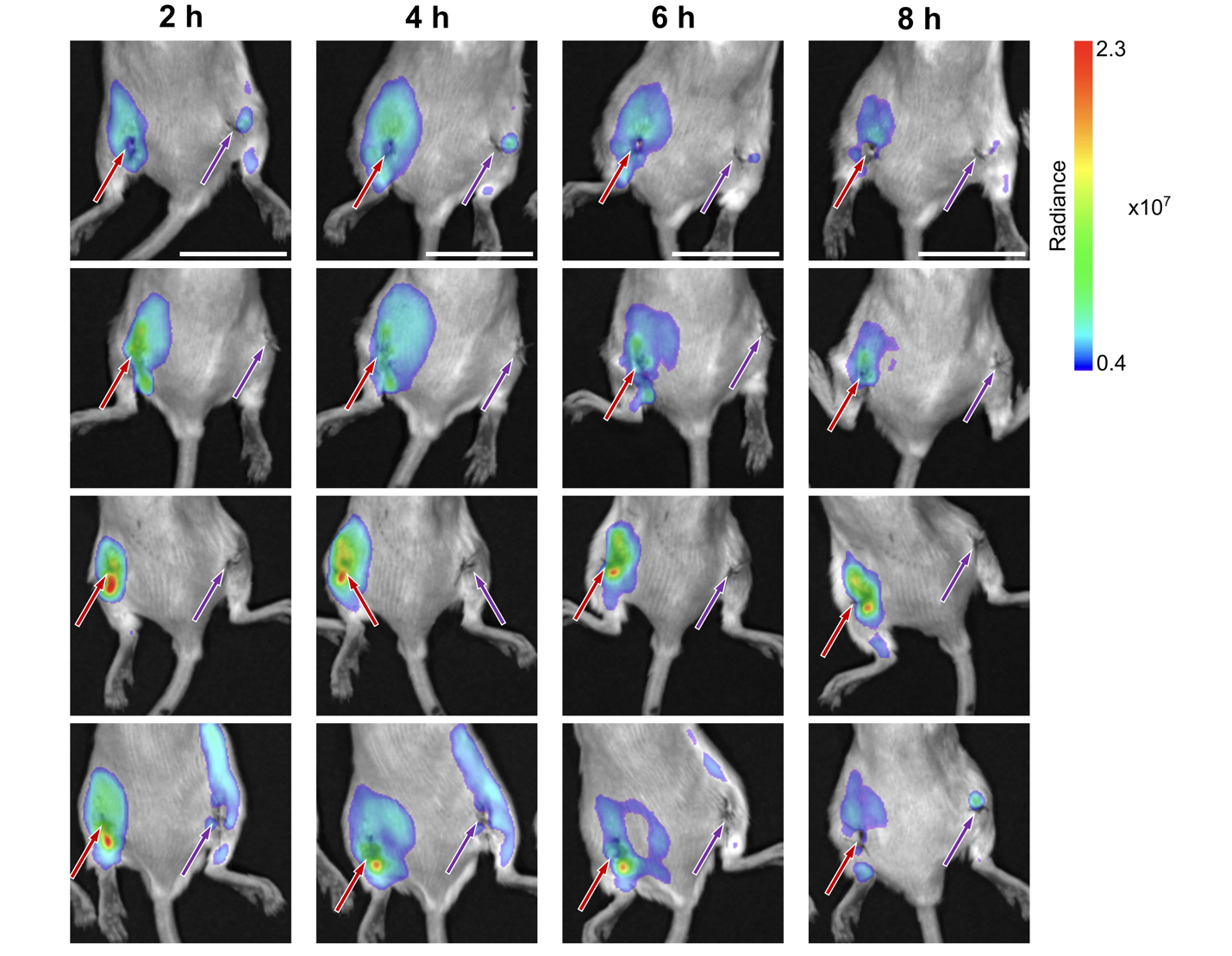
**Figure S8.** *In vivo* fluorescent imaging of mouse subcutaneous wound infection model. Approximately 5 x 10^7^ CFU *S. aureus* USA300 was injected subcutaneously at the site of an incision over the left thigh (red arrow), and an equivalent volume of sterile normal saline was injected at the site of an incision over the right thigh (blue arrow). After 1.5 hours, 100 µL of **10** (100 µM, 30% PEG-400 in 1X PBS) was injected retro-orbitally. Fluorescence was detected using 535 nm excitation and 610 nm emission filter at 2, 4, 6, and 8 hours after injection of the probe. Scale bar: 2 cm.

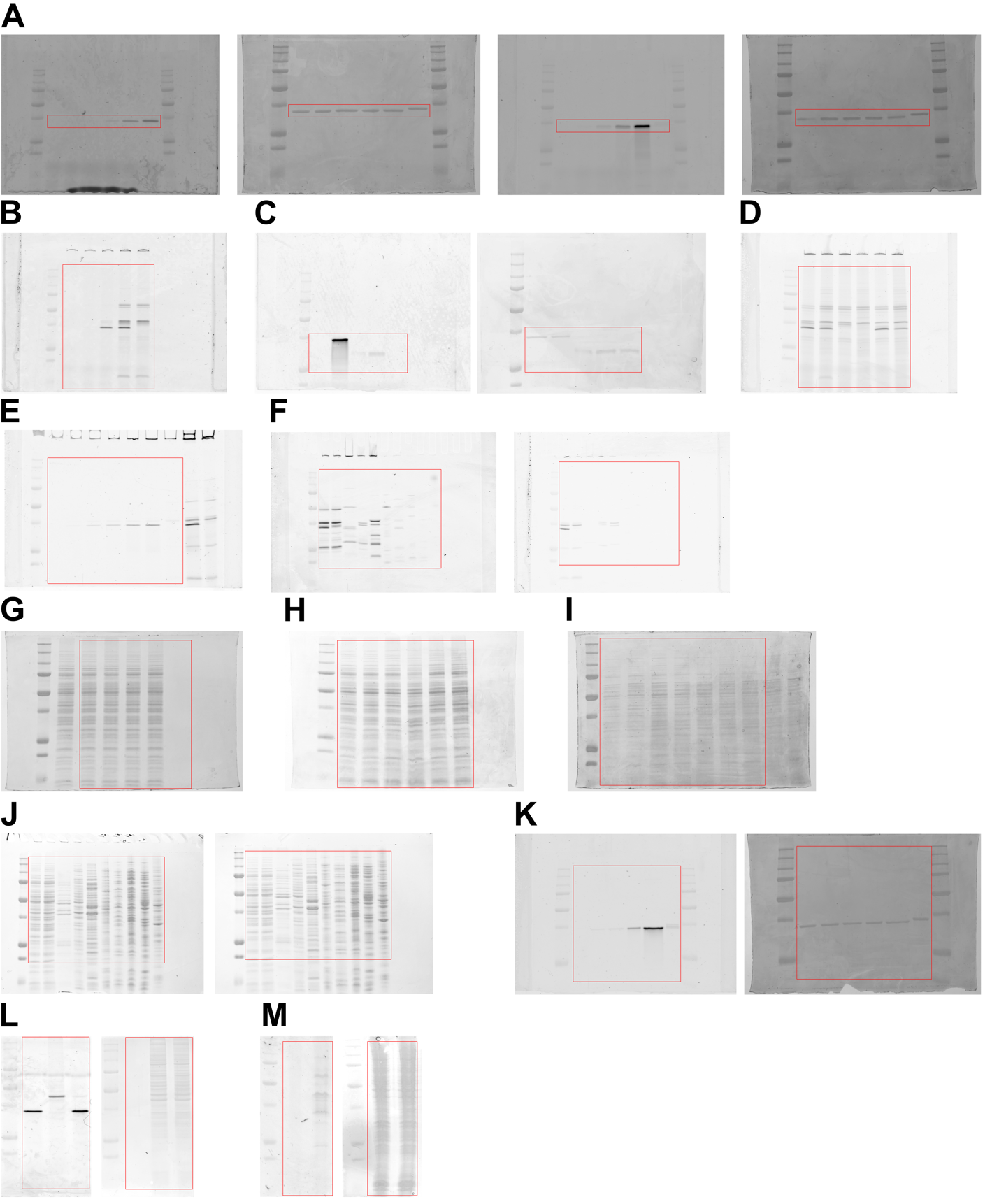
**Figure S9.** Uncropped gels for (A) Figure 2B, (B) Figure 4B, (C) Figure 4C, (D) Figure 4D, (E) Figure 5B, (F) Figure 6A, (G) Figure S1A, (H) Figure S1B, (I) Figure S1C, (J) Figure S1D, (K) Figure S3, (L) Figure S7A, and (M) Figure S7B. Each cropped region is highlighted by a red box.

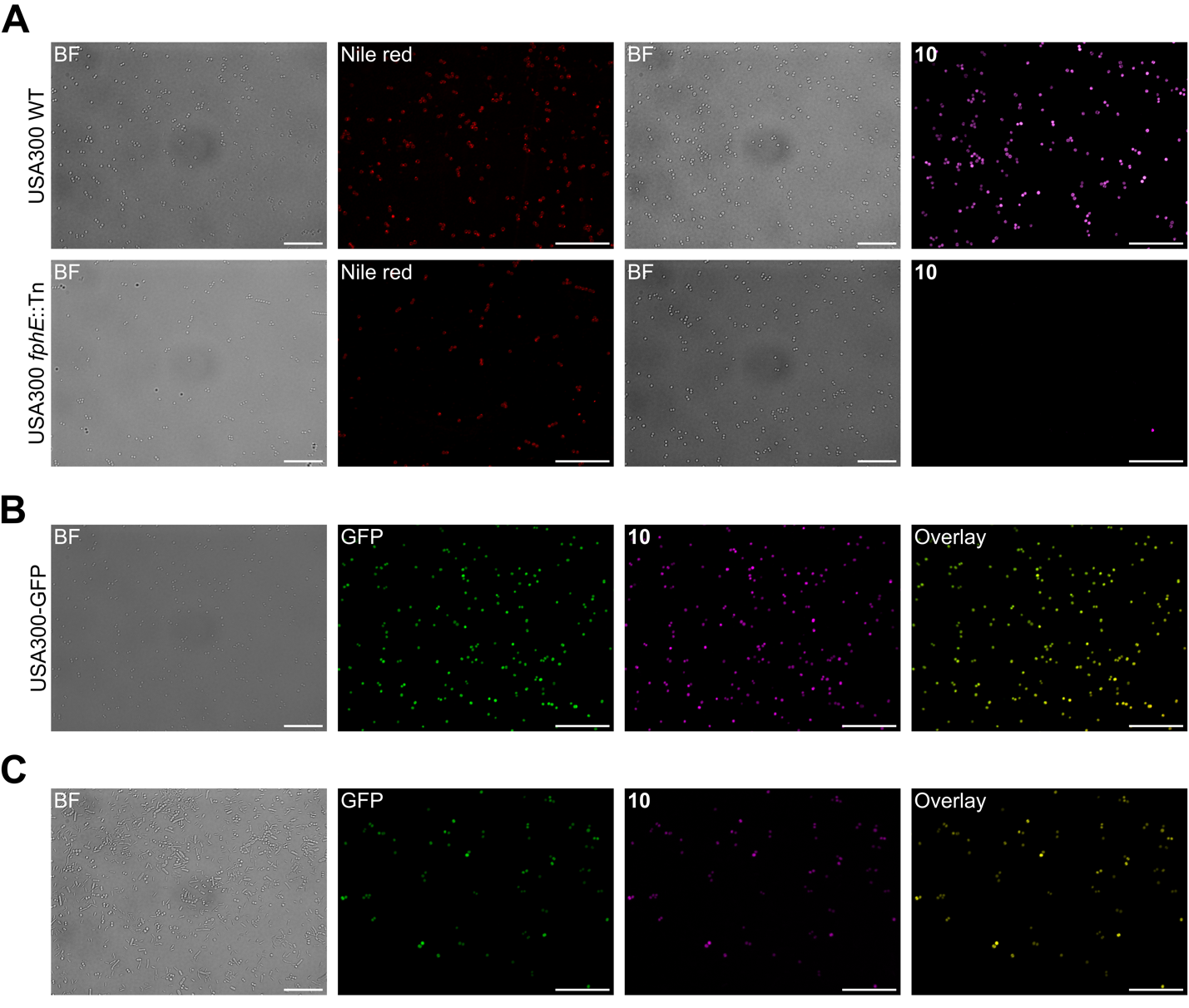
**Figure S10.** Uncropped microscopy images for (A) Figure 5C, (B) Figure 5F, and (C) Figure 6B. Scale bar: 20 µm.

**2. Materials and Methods**

**FphE cloning**

The full length *fphE* (currently annotated as Uncharacterized hydrolase Q2FDS6 Y2518_STAA3 in UniProt,^3^ gene loci SAUSA300_2518) was amplified from the *S. aureus* USA300 genome using primers CAG GGA CCC GGT ATG GAA ACT TTA GAA TTA CA and CGA GGA GAA GCC CGG TTA ACC CCA CAT ATT TAA TAA TA that introduced overhangs for ligation-independent cloning.^4^ The PCR product was gel-purified cloned into modified pET28a-LIC vectors incorporating an N-terminal His_6_-tag and a 3C protease cleavage site.

Ser103Ala mutant FphE (S103A) was generated by site directed mutagenesis using primer TAT ATA TTA GGT TCA GCG TCA GGT TCA ATC GTT GCG ATG C and confirmed by sequencing.

**Protein expression and purification**

Chemically competent BL21 (DE3) *E. coli* was transformed by the pet28a-(pETNKI-his-3C-LIC-kan) FphE or mutant FphE (S103A) expression vector. An overnight culture of the transformed bacteria in LB (+ Kanamycin 50 μg/mL) selection medium was diluted 1:100 into 2 L of selection medium at 37 °C, 220 rpm. The culture was induced for protein expression at OD_600_ 0.6–0.7 by adding 0.2 mM isopropyl β-D-1-thiogalactopyranoside (IPTG) and grown overnight at 18 °C with shaking (180 rpm).  For purification, cells were harvested by centrifugation and bacterial pellets was stored at −80 °C. The following day, each pellet was thawed on ice, resuspended in 30 mL lysis buffer (20 mM Tris pH 8.0, 300 mM NaCl, 0.05 mg/mL lysozyme), and lysed by sonication (3 times, 1 min each: pulse ON 4 sec. pulse OFF 1 sec, 50% amplitude, Branson Sonifier; Branson, Danbury, CT). Lysates were centrifuged at 15,000 rpm in an Avanti JA-17 rotor (Beckman Coulter Life Sciences, Indianapolis, IN) for 45 min, and the supernatant transferred to Ni-NTA resin pre-equilibrate with equilibration buffer (50 mM Tris pH 8.0, 300 mM NaCl, 5 mM imidazole, 10% glycerol) and incubated at 4 °C, rotating for 60 min, then purified by gravity flow. The resin was washed three times with washing buffer (50 mM Tris pH 8.0, 300 mM NaCl, 5 mM imidazole, 10% glycerol) and the His_6_-tagged protein was eluted in 50 mM Tris pH 8.0, 300 mM NaCl, 300 mM imidazole, 10% glycerol buffer. Eluates were analyzed by SDS-PAGE (Coomassie stain). To remove the His_6_-tag from the wild-type (WT) protein, the elutes were incubated with 1 μg 3C-protease for 100 μg protein in presence of 2 mM DTT overnight at 4°C. The complete cleavage was confirmed by SDS-PAGE. The 3C-protease-cleaved rFphE was then purified by size exclusion chromatography using S200 10/300 column (EMD Millipore, Billerica, MA) with the elution buffer (20 mM HEPES pH 7.0 and 100 mM NaCl). For crystallography, rFphE underwent a two-steps purification, first by anion exchange (RESOURCE^TM^ Q, 10 mM HEPES pH 7.5, gradient 10 to 1,000 mM NaCl) and second by size exclusion chromatography (10 mM HEPES pH 7.5, 100 mM NaCl) using a Superdex 75 Increase column (GE Life Sciences, Pittsburgh, PA). Finally, purified rFphE was either used directly for crystal drops or flash-frozen for long-term storage at −80 °C. The concentrations of purified WT and mutant (S103A) rFphE were determined by Bradford assay.

**Substrate screening and enzyme inhibition assay**

The enzyme activity of the purified enzymes was tested using series of 4-methylumbelliferyl (4-MU)-based fluorogenic substrates as described earlier.^5-7^ Briefly, 0.5 nM of WT or mutant (S103A) rFphE were added 50 μM solution of each substrate mixed in 1X phosphate buffered saline (PBS; Corning, Manasas, VA) with 0.02% TritonX-100 (Fisher Scientific, Fairlawn, NJ). Fluorescence (λ_ex_ = 365 nm and λ_em_ = 455 nm) was then read at 30 °C in 1-min intervals for 60 min using a Cytation 3 Multi-Mode Reader (BioTek, Winooski, VT). Turnover rates in the linear phase of the reaction (10–20 min) were calculated using Graphpad Prism9 as RFU/min. Rates were normalized by subtracting background hydrolysis rates measured for each substrate in reaction buffer in the absence of protein.

The inhibitory activity of chemical probes was determined by incubating WT rFphE (0.5 nM) with different concentrations of the inhibitor and quantifying residual activity using the 20 μM fluorogenic substrate 4-methylumbelliferyl (4-MU) octanoate. Inhibitors at various concentrations were preincubated with rFphE at room temperature (rt) for 1 h, and rFphE activity was measured by monitoring the change in fluorescence intensity after adding 4-MU octanoate for 1 h using a Cytation 3 Multi-Mode Reader (λ_ex_ = 365 nm and λ_em_ = 455 nm). Calculations were performed using GraphPad prism 9 software. Sigmoidal curves were fitted to the data using the following dose-response equation:

$$y=A1+(A2-A1)/(1+\left( \frac{x}{IC50} \right))$$

(x, inhibitor concentration; y, percent activity of the reaction; A1, 100%, y value at the top plateau; A2, 0%, y value at the bottom plateau; used standard slope of −1.0, Hill coefficient). IC_50_ values were derived from the fitted curve.

**Gel labeling analysis of purified recombinant proteins**

Purified WT (2.0 µM) and S103A rFphE (2.0 µM) were incubated with the indicated concentrations of each chemical probe for 1 h at 37 ºC. For samples treated with alkyne probes, 20 µL of resulting mixtures were added 2.16 µL of freshly prepared click mix (0.5 µL of 50 mM CuSO_4_ in H_2_O, 1.16 µL of 100 mM BTTAA in DMSO, and 0.5 µL of N_3_-TAMRA in DMSO) and 1.16 µL of 300 mM sodium ascorbate solution in H_2_O. After incubation for 30 min at 37 ºC, 20 µL of 4X SDS loading buffer was added and samples were boiled at 100 ºC for 5 min. The samples treated with fluorescent probe was directly subjected by 4X SDS loading buffer without click reaction and boiled at the same condition with alkyne probes. Then, samples were analyzed by SDS-PAGE (12%) running at 120 V in an electrophoresis chamber under the ambient temperature. Fluorescence gels were visualized using a GE Typhoon FLA 9000 (GE Healthcare, Pittsburgh, PA) followed by staining using Coomassie.

**Mass spectrometry validation of FphE labeling with JJ-OX-004 (9)**

2 µM WT rFphE was incubated with 200 µM of **9** (from a 10 mM stock in DMSO) in 100 µL 1X PBS (pH=7.4) with 2% (v/v) DMSO, at room temperature for labeling. After 4 h, 10 µL of the labeled sample was directly used and analyzed on an Agilent 1200 HPLC equipped with an Agilent Zorbax SB-C3 column (1.8 µm, 2.1 x 150 mm, pore size 300 Å) coupled to an Agilent 6125B Single Quad Mass Spectrometer (Agilent Technologies, Santa Clara, CA). For LC-MS conditions, 95% solvent A (0.1% formic acid in LC-MS grade water), 5% solvent B (0.1% formic acid in LC-MS grade acetonitrile) at 0.6 mL/min was used for 2 min to remove excess of salts into waste before a linear 6-minute gradient from 80% solvent A (20% solvent B) to 20% solvent A (80% solvent B), followed by a one-minute 95% solvent A (5% solvent B). Protein labeling efficiency was quantified by the UV. The acquired mass of protein was deconvoluted using the deconvolution software in Agilent Bioanalysis Software package.

**Jump dilution assay**

The assay was performed by incubating 400 nM of compound **9** with 50 nM WT rFphE in assay buffer (0.02% TritonX-100 in 1X PBS) for 1 h at ambient temperature. A 0.5 μL portion of each sample was then diluted (100-fold, final concentration of 0.5 nM rFphE) into a 50 μL solution containing 20 μM of 4-MU octanoate substrate. Enzyme activity was recorded by monitoring the change in fluorescence intensity for 1 h using a Cytation 3 Multi-Mode Reader (λ_ex_ = 365 nm and λ_em_ = 455 nm). As a control, compound **9** were also tested in 4 nM and 400 nM concentrations with 0.5 nM rFphE and 20 μM of 4-MU octanoate. The total volume/well and final concentrations of enzyme and substrates were kept constant between the controls and jump dilution samples. Fluorescent intensities were plotted against time using GraphPad Prism 9.

**FphE crystallization**

FphE was broad screened for crystallization using commercially available screens and hits were further optimized manually. For the unbound FphE structure 0.2 µL of 15 mg/mL FphE (9 mM HEPES pH 7.5, 87 mM NaCl, 13% DMSO) were mixed with 0.2 µL of reservoir solution. Sitting drop reservoir contained 25 µL of 0.18 M magnesium chloride, 0.1 M Tris pH 7.5, 22.5% w/v polyethylene glycol monomethyl ether 2000. The crystal was frozen in a solution of ~25% glycerol, 75% reservoir. For compound **9** (JJ-OX-004) bound, 13 µL of 19 mg/mL FphE (10 mM HEPES pH 7.5, 100 mM NaCl) were mixed with 5 µL compound **9** (10 mM in DMSO) and incubated at 18 °C overnight. 0.15 µL FphE-compound **9** solution was mixed with 0.2 µL of reservoir solution. Sitting drop reservoir contained 25 µL of 0.18 M magnesium chloride, 0.1 M Tris pH 8.5, 22.5% w/v polyethylene glycol monomethyl ether 2000. The crystal was frozen in a solution of ~25% glycerol, 75% reservoir.

**FphE data collection, processing, refinement, deposition, and analysis**

X-ray diffraction data were collected at the Australian synchrotron MX1^8^ and MX2^9^ beamline. Datasets were processed with XDS,^10^ merging and scaling were performed using AIMLESS.^11^ Phases were initially solved with Phenix Phaser molecular replacement^12^ using a model form Alphafold^13^ via Uniprot^3^ for FphE *S. aureus* strain USA300 AF-Q2FDS6-F1. Model building and refinement were conducted in COOT^14^ and Phenix.^15^ The final structure was deposited to the worldwide protein databank^16^ PDB ID 8T87 and 8T88. Statistics for the datasets are listed in Table S2 and S3. Structure figures, analysis and alignments were created with UCSF Chimera^17^ and LigPlot.^18^

**Bacterial culture and bacterial growth inhibition assay**

The bacterial strain *Staphylococcus aureus* USA300 were cultured on tryptic soy agar (TSA) plates by overnight incubation at 37 °C. A single colony was transferred to tryptic soy broth (TSB; Sigma-Aldrich, St. Louis, MO). The primary culture was grown to exponential phase (OD_600_ = 0.2) at 37 °C and suspension was diluted 1:100. An inoculum of 100 μL was introduced to a treatment plate containing two-fold serial dilutions of tested compounds in TSB or Mueller-Hinton broth 2 (MHB2, cation adjusted; Millipore Sigma, Bedford, MA) media (100 μL of treatment/well) to give a final total volume of 200 μL/well and a final inoculum density of ~5 × 10^5^ CFU/mL. The completed assay plate was incubated at 37°C and the growth data were recorded, with a Cytation 3 Multi-Mode Reader.

**SDS-PAGE analysis of live *S. aureus* labeling**

Wild-type (WT) and *fphE* transposon mutant (*fphE*::Tn) USA300 *S. aureus* cells were cultured on TSA plates by overnight incubation at 37 °C. Three colonies from each strain were picked and transferred to TSB media (10 mL). The culture was grown to stationary phase shaking at 37 °C. Bacterial cultures were spun down at 5,000 g for 10 min and the supernatant was removed. The pellets were washed twice in 1X PBS and resuspended in the same buffer (5 mL). The sample was aliquoted into 400 µL portions. Cells were preincubated with the indicated concentration of alkyne probe **9** or fluorescent probe **10** for 2 h at 37 ºC. After washing twice with 1X PBS, cells were resuspended in 1X PBS with (for **9** treated samples) or without 0.02% TritonX-100 and lysed by bead-beating at 4 ºC. For compound **9** treated samples, 20 µL of lysed cells in buffer were added 2.16 µL of freshly prepared click mix (0.5 µL of 50 mM CuSO_4_ in H_2_O, 1.16 µL of 100 mM BTTAA in DMSO, and 0.5 µL of N_3_-TAMRA in DMSO) and 1.16 µL of 300 mM sodium ascorbate solution in H_2_O. After incubation for 30 min at 37 ºC, 20 µL of 4X SDS loading buffer was added and samples were boiled at 100 ºC for 5 min. The prepared samples labeled by **10** were directly added by 4X SDS loading buffer without click reaction and boiled at 100 ºC for 5 min. The denatured samples were allowed to cool to ambient temperature and analyzed by SDS-PAGE (12%) running at 120 V in an electrophoresis chamber. Protein concentrations for each sample were quantified by using the BCA assay kit (Pierce/Thermo Scientific, Rockford, IL), and the indicated amounts of total protein were loaded into each well. Fluorescence gels were visualized using a GE Typhoon FLA 9000, followed by staining using Coomassie.

**Protease protection assay**

WT, *fphB*::Tn, and *fphE*::Tn USA300 *S. aureus* strains were cultured, grown and collected by following the same procedure as described above. Proteinase K solution (20 mg/mL; Goldbio, St. Louis, MO) was either added (0.5 mg/mL final concentration) to 400 µL aliquot of each sample or left unaltered in the control group. Samples were incubated at 37 ºC for 16 h, then spun down at 12,000 g for 1 min. The supernatant was removed, and cell pellets were washed twice with 1X PBS. Cells were incubated with the alkyne probe **9** (1.0 µM) at 37 ºC for 2 h, washed with 1X PBS twice, resuspended in 1X PBS with 0.02% TritonX-100, and lysed by bead-beating at 4 ºC. Then, 20 µL of lysed samples were labeled by incubating for 30 min at 37 ºC with 2.16 µL of freshly prepared click mix (0.5 µL of 50 mM CuSO_4_ in H_2_O, 1.16 µL of 100 mM BTTAA in DMSO, and 0.5 µL of N_3_-TAMRA in DMSO) and 1.16 µL of 300 mM sodium ascorbate solution in H_2_O. Clicked samples were denatured by boiling at 100 ºC for 5 min after addition of loading buffer (4X SDS, 20 µ). Prepared samples were cooled and analyzed by SDS-PAGE (12%) running at 120 V in an electrophoresis chamber under the ambient temperature. Protein concentrations for each sample were quantified by using the BCA assay kit, and 10 µg of total protein were loaded into each well. The gel fluorescence was scanned by using a GE Typhoon FLA 9000, followed by Coomassie staining.

**Plate-reader-based assay for measuring labeling of live *S. aureus***

WT and *fphE*::Tn *S. aureus* USA300 cells were cultured and grown shaking at 37˚C in 10 mL TSB. After overnight growth, samples were pelleted at 5,000 g for 10 min and washed twice with 20 mL 1X PBS pH 7.4. The washed samples were resuspended in 5 mL of 1X PBS. 100 nM fluorescent probe **10** was added in both the samples. Subsequently, each sample was aliquoted in 500 µL of multiple tubes and incubated at 37˚C. One tube from each WT and *fphE*::Tn *S. aureus* USA300 were taken out every 15 min and unbound probe was removed by washing twice by pelleting samples at 12,000 g for 2 min and resuspending in 500 µL 1X PBS. 50 µL of samples with 20 µL of 0.02% TritonX-100 in 1X PBS buffer were added in each well in a 384-well plate and fluorescence was measured using a Cytation 3 Multi-Mode Reader (λ_ex_ = 552 nm and λ_em_ = 578 nm).

**Confocal microscopy**

*S. aureus* USA300, *S. aureus fphE*::Tn, or GFP expressing *S. aureus* USA300 + pCM29, were grown shaking at 37˚C in 10 mL TSB. After overnight growth, samples were pelleted at 5,000 g for 10 min and washed twice with 1X PBS pH 7.4. After two washes, samples were resuspended in 5 mL of 1X PBS. 100 nM **10** was added to a 500 µL aliquot of each sample. Samples were incubated for 2 h at 37 ˚C and washed twice with 1X PBS buffer. Identical samples that were not labeled with **10** were incubated with Nile Red 10 µg/mL for 10 min in the dark and washed with 1X PBS buffer. The diluted pellet in 1X PBS buffer was immobilized on glass slides immediately prior to analysis. Confocal microscopy was performed using Zeiss Axio Observer Z1 LSM 700 (Carl Zeiss GmbH, Jena, Germany), and fluorescent images were taken in the same field of view as corresponding bright field images. Nile Red imaging was performed with excitation at 555 nm, power 5.0%, and gain 800. GFP imaging was performed with excitation at 488 nm, power 0.2%, and gain 700. TAMRA imaging was performed with excitation at 555 nm, power 0.2%, and gain 700. All fluorescence images were minimally processed using ImageJ software.

**Labeling of different bacterial strains**

Bacteria were grown shaking at 37 ˚C in 10 mL TSB for *S. aureus* USA300, *S. aureus* USA300 + pCM29, *S. aureus fphE*::Tn*, Staphylococcus epidermidis* ATCC 12228*, Pseudomonas aeruginosa* ATCC 27853, *Escherichia coli* BW25113, *Salmonella enterica* LT2, Brain Heart Infusion broth (Research Proucts International, Mt. Prospect, IL) for *Streptococcus pyogenes* ATCC 12384, *Listeria monocytogenes*, *Klebsiella aerogenes* ATCC 13048, or Yeast Extract Peptone Dextrose broth (Fisher Scientific, Fair Lawn, NJ) for *Candida albicans*. The GFP-expressing *S. aureus* USA300 + pCM29 strain was grown with the addition of 10 µg/mL chloramphenicol (Fluka Chemie GmBH, Buchs, Switzerland) to maintain the plasmid. After overnight growth, samples were pelleted at 3,000 rpm for 10 min and resuspended in 20 mL phosphate-buffered saline pH 7.4 (PBS; Corning, Manassas, VA). After two such washes, samples were resuspended in 4 mL of 1X PBS.

100 nM **10** was added to a 1 mL aliquot of each sample, and 100 nM fluorophosphonate-TAMRA (FP-TMR) was added to a second 1 mL aliquot of each sample. These aliquots were then incubated at 37˚C for 2 hours, washed, lysed, and prepared for SDS-PAGE as described above.

The OD_600_ of the remaining unlabeled samples was measured and additional 1X PBS added to samples as needed such that the final OD_600_ of all samples was the same. 100 µL of each bacterial strain was then added to a mixture. Three such mixtures were made: 1) the *S. aureus* strain used was USA300 + pCM29, 2) the *S. aureus* strain used was the *fphE*::Tn mutant, 3) 100 µL of PBS was used in place of *S. aureus*. 100 nM **10** was added, and samples incubated at 37˚C for 2 h. Unbound probe was removed by washing twice by pelleting samples at 12,000 g for 2 min and resuspending in 500 µL 1X PBS. After final wash, samples were resuspended in remaining supernatant after decanting. Identical mixtures that were not labeled with **10** were stained wile Nile Red 50 µg/mL for 10 min in the dark and washed as before. 3 µL of samples were placed on poly-L-lysine microscope slides (Polysciences Inc., Warrington, PA), covered with a cover slip, sealed on two sides with nail polish, and imaged on a confocal microscope (Zeiss Axio Observer Z1 LSM 700) at 63x with oil immersion. Fluorescent images were taken in the same field of view as corresponding brightfield images. Nile Red imaging was performed with excitation at 555 nm, power 5.0%, and gain 800. GFP imaging was performed with excitation at 488 nm, power 0.2%, and gain 700. TAMRA imaging was performed with excitation at 555 nm, power 0.2%, and gain 700.

**Quantitative analysis of microscopy images**

A multi-step methodology including image pre-processing, segmentation for cell image analysis was employed. The initial step involves reading the image and converting it to grayscale, followed by the application of Otsu thresholding for segmentation and generating a binary image. Otsu's method, an automatic thresholding technique, selects an optimal threshold by maximizing the variance between two classes of pixels in a grayscale image. To enhance segmentation, morphological operations are performed, including the removal of small objects, closing gaps in cell boundaries, and filling holes within cells. Subsequently, the watershed transform is applied to the binary image to effectively separate each cell. The watershed transform, a computer vision image segmentation technique, treats pixel intensities as elevations in a topographical landscape, allowing for the delineation of object boundaries based on the dynamics of water flowing into catchment basins. The resulting segmented cells are displayed by overlaying blue boundaries onto the original image. For the cell analysis, the major axis is calculated and plotted for each detected cell individually, visualizing it on the overlayed image.

**Mouse subcutaneous infection model**

All animal models were approved by the Stanford Institutional Animal Care and Use Committee. *S. aureus* USA300 culture was prepared by inoculating a single colony in TSB overnight at 37˚C. 100 µL of the overnight culture was added to 10 mL of TSB media and cultured until exponential growth phase (OD_600_ = 0.5). Cultures were then centrifuged, pellets washed three times with 1X PBS, and suspended to a concentration of 5 × 10^9^ CFU/mL.

Female BALB/c mice, 6–8 weeks old, were anesthetized by isoflurane inhalation. A small incision was made on left thigh, and 10 µL of bacterial culture (to yield inoculum of 5 × 10^7^ CFU) was injected subcutaneously at the incision site of the left thigh. An incision was then made on the right thigh, and 10 µL of sterile normal saline was injected subcutaneously at the incision site. All incisions were closed with 5-0 Prolene sutures in an interrupted fashion. JJ-OX-007 (**10**) was prepared in 30% PEG-400 and PBS for a final concentration of 100 mM. Mice were injected retro-orbitally with 100 µL of **10** (for total 10 nmol injection) 2 h after inoculation. Mice were serially imaged under anesthesia in an IVIS (LagoX from Spectral Instruments, Tucson, AZ) with excitation at 535 nm, medium binning, long pass filter at 610 nm, and up to 120 second exposure time. Imaging was performed at 2 h, 4 h, 6 h, and 8 h after injection of probe. For quantitative analysis, Aura software (Spectral Instruments) was used to compare fluorescence efficiency from entire infected left thigh was compared to fluorescence efficiency over the entire uninfected right thigh. Fluorescence signals were quantified as total efficiency, which is equivalent to the mean signal efficiency of the region of interest (ROI) × area of the ROI, where efficiency is defined as a ratio of detected emission radiance over LED excitation radiance.

**HEK293T cell labeling**

HEK293T cells (ATCC CRL-11268) were grown in Dulbecco's Modified Eagle Medium (DMEM; Gibco 11995065) supplemented with 10% fetal animal serum (FAS). Cells were grown in a humidified incubator at 37 ºC and 5% CO_2_ atmosphere. For intact cell labeling, HEK293T cells were seeded at 40% confluency in six-well dishes. After 24 h, the cell growth medium was removed and replaced with fresh medium containing either DMSO vehicle or **10** (100 nM). Following incubation for 2 h, media was removed, and each well was rinsed once with 1X PBS. Cells were harvested by scraping into pre-cooled 1.5-mL Eppendorf tubes. Cells were pelleted by centrifugation (5 min, 2,000 g, 4 ºC). Pellets were washed two times with ice cold 1X PBS at which point cell pellets were resuspended in lysis buffer (1% SDS in 1X PBS) and lysed via tip sonication (3 times, pulse ON 5 sec and pulse OFF 5 sec). Lysate was cleared by centrifugation (10 min, 10,000 g). The supernatant was moved to a fresh tube and protein concentration was determined by BCA assay kit. Gel samples were prepared at a concentration of 2 mg/mL with appropriate volumes of 1% SDS buffer and 4X reducing loading buffer and boiled for 5 min at 95 ºC. For each sample, 3.5 µL (7 µg) were separated on a 12% SDS-PAGE gel. Fluorescence was visualized using a GE Typhoon FLA 9000 followed by staining using Coomassie.

For lysate cell labeling, HEK293T cells grown in 10-mL dishes. The cell media was aspirated, and cells were washed with 1X PBS before being harvested by scraping into pre-cooled 15-mL Falcon tubes. Cells were pelleted by centrifugation (5 min, 2,000g, 4 ºC) and washed two times with ice cold 1X PBS. Cells were then resuspended in ice cold PBS and lysed via tip sonication (3 times, pulse ON 5 sec and pulse OFF 5 sec). Lysate was centrifuged (10 min, 10,000g) and the supernatant was moved to a fresh tube on ice. Protein concentration was determined by BCA assay. Labeling was conducted under the following three conditions: (1) HEK293T lysate (7 µg) with rFphE (30 ng), (2) HEK293T lysate (7 µg) alone, or (3) rFphE (30 ng) alone. For all three, **10** (100 nM) was added and proteins were incubated at 37 ºC for 2 hours. After which, gel samples were prepared by adding 4X reducing loading buffer and boiling for 5 minutes at 95 ºC. Each sample was separated on a 12% SDS-PAGE gel. Fluorescence was visualized using a GE Typhoon FLA 9000 followed by staining using Coomassie.

**3. Chemical Synthesis**

**General methods**

Unless noted otherwise, reagents and solvents were purchased from commercial suppliers and were used without further purification. All reactions were performed under argon atmosphere and stirring unless otherwise noted. Reaction flasks were dried overnight at 100 °C in an oven. Flash column chromatography was carried out using SiliaFlash® P60 (230–400 mesh, SiliCycle) with the indicated solvents of reagent-grade. Thin-layer chromatography (TLC) was performed using pre-coated 0.25 mm silica gel plates (Merck). ^1^H and ^13^C NMR spectra were recorded on BRUKER AVANCE III 500 spectrometer as solutions in the indicated solvents at room temperature (rt). Chemical shifts were quoted in parts per million (ppm, δ) downfield from tetramethylsilane (TMS) and were referenced to the deuterated solvent. ^1^H NMR data were reported in the order of chemical shift, multiplicity (s, singlet; brs, broad singlet; d, doublet; dd, doublet of doublets; t, triplet; td, triplet of doublets; q, quartet; qd, quartet of doublets; quint, quintet; m, multiplet and/or multiple resonance), number of protons, and coupling constant in hertz (Hz). High-resolution mass spectra were analyzed by LC-ESI/MS on a Waters Acquity UPLC system coupled to a Thermo Exploris 240 Orbitrap mass spectrometer. The following abbreviations for reagents and solvents are used: *N*,*N*-dimethylformamide (DMF), dimethyl sulfoxide (DMSO), *N*-methyl-2-pyrrolidone (NMP), petroleum ether (PE), tetrahydrofuran (THF).

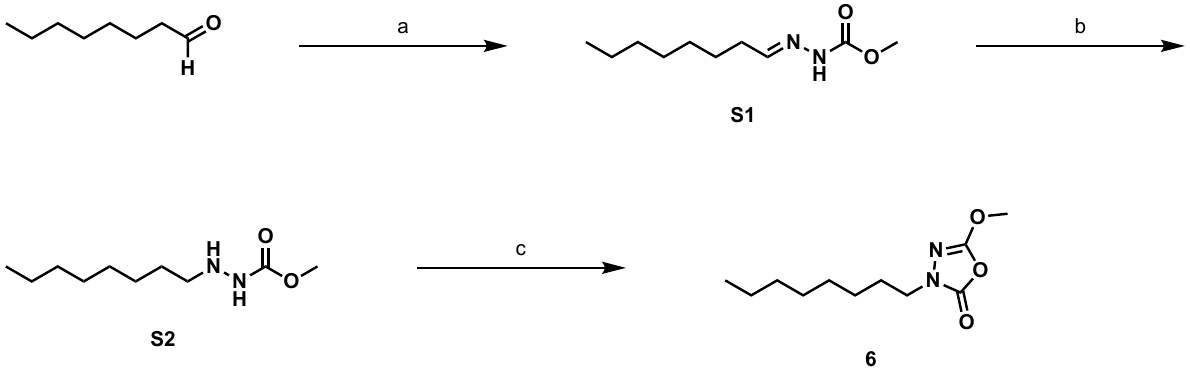

**Scheme S1**. Synthesis of JJ-OX-001 (**6**). Reagents and conditions: (a) methyl carbazate, AcOH, MeOH, reflux; (b) NaBH_3_CN, 3 N HCl/MeOH, MeOH, 0 ºC, 41% for 2 steps; (c) triphosgene, pyridine, CH_2_Cl_2_, 0 ºC to rt, 7%.

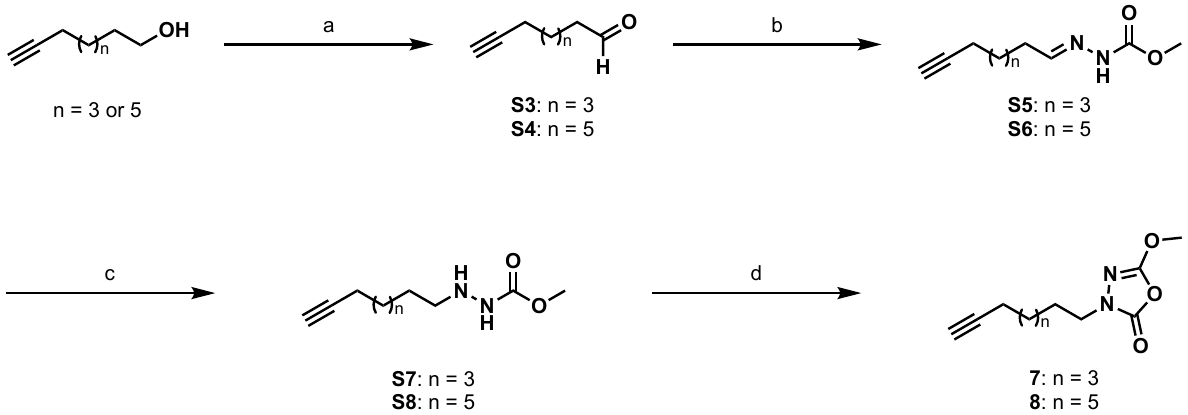

**Scheme S2**. Synthesis of JJ-OX-002 (**7**) and JJ-OX-003 (**8**). Reagent and conditions: (a) SO_3_·py, Et_3_N, DMSO, CH_2_Cl_2_, 0 ºC to rt, 75-82%; (b) methyl carbazate, AcOH, MeOH, reflux; (c) NaBH_3_CN, 3 N HCl/MeOH, MeOH, 0 ºC, 35-63% for 2 steps; (d) triphosgene, pyridine, CH_2_Cl_2_, 0 ºC to rt, 2-4%.

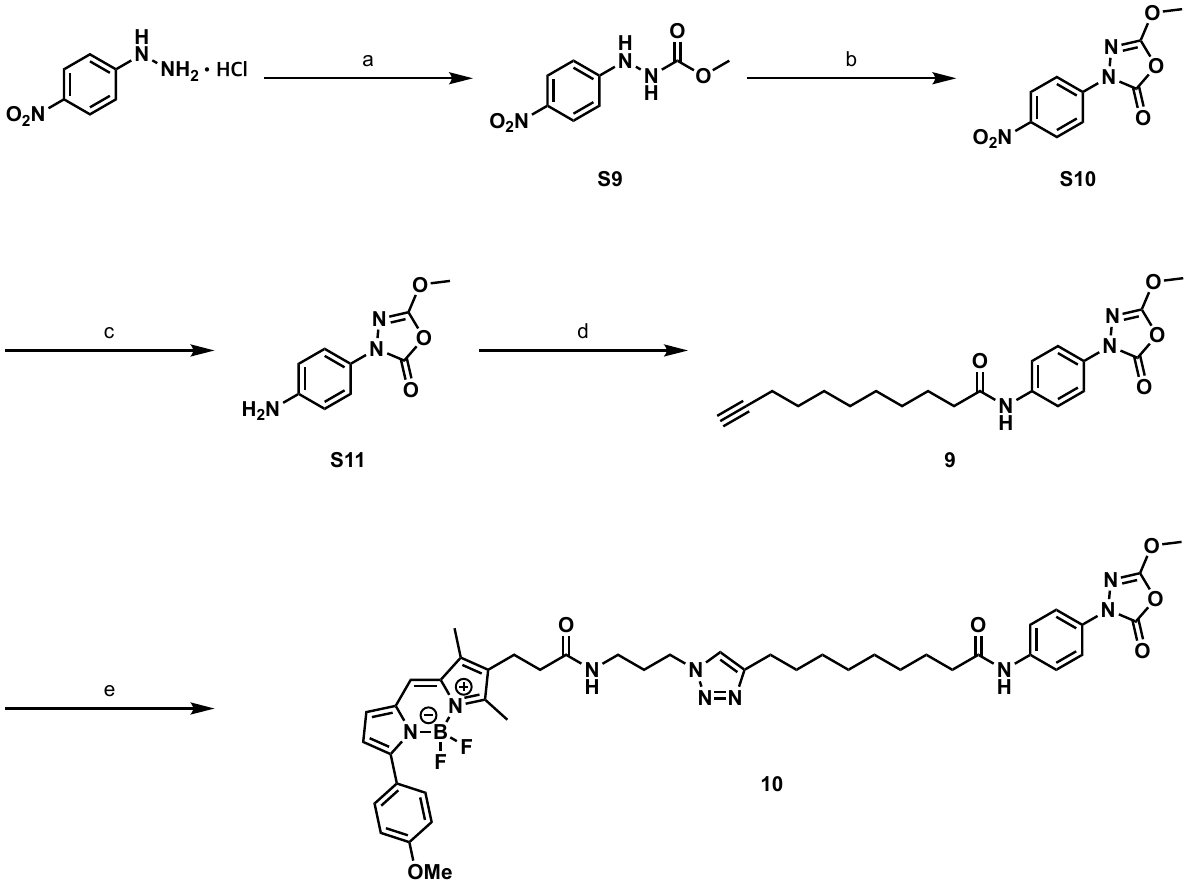

**Scheme S3**. Synthesis of JJ-OX-004 (**9**) and JJ-OX-007 (**10**). Reagent and conditions: (a) ClCOOMe, NMP, pyridine, 0 ºC; (b) triphosgene, pyridine, CH_2_Cl_2_, 0 ºC to rt, 17% for 2 steps; (c) H_2_, Pd/C, MeOH/MeOH, 61%; (d) 10-undecynoic acid, (COCl)_2_, Et_3_N, DMF, CH_2_Cl_2_, 87%; (e) BDP-TMR azide, CuSO_4_·5H_2_O, ascorbic acid, DMF, rt, 34%.

**Synthesis of JJ-OX-001 (6)**

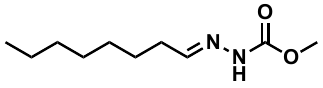

Methyl (*E*)-2-octylidenehydrazine-1-carboxylate (**S1**)

To a stirred solution of octanal (1.56 mL, 10.0 mmol) in MeOH (10.0 mL), methyl carbazate (901 mg, 10.0 mmol) and AcOH (57.2 µL, 1.00 mmol) were added at ambient temperature. After being refluxed for 12 h, the reaction mixture was cooled and then concentrated in vacuo. The residue was recrystallized from MeOH/PE to afford 1.39 g (69%) of **S1** as a white solid. Obtained **S1** was directly used for the next step.

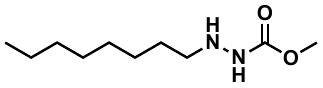

Methyl 2-octylhydrazine-1-carboxylate (**S2**)

To a stirred solution of **S1** (974 mg, 4.86 mmol) in MeOH (10.0 mL) was added NaBH_3_CN (764 mg, 12.2 mmol) and HCl (3 M in MeOH, 7.30 mL) at 0 ºC. After being stirred for 4 h at the same temperature, the reaction mixture was quenched with saturated NaHCO_3_ until basic pH and extracted with EtOAc. The organic layer was dried over MgSO_4_ and concentrated in vacuo. The resulting residue was purified by flash column chromatography on silica gel (EtOAc/*n*-hexane = 1:2) to afford 570 mg (59%) of **S2** as a colorless oil: ^1^H NMR (500 MHz, CDCl_3_) δ 3.71 (s, 3H), 2.84 (t, *J* = 7.3 Hz, 2H), 1.45 (quint, *J* = 7.3 Hz, 2H), 1.34 – 1.22 (m, 10H), 0.87 (t, *J* = 6.7 Hz, 3H); ^13^C NMR (126 MHz, CDCl_3_) δ 158.0, 52.5, 52.2, 31.9, 29.6, 29.4, 27.8, 27.2, 22.8, 14.2.

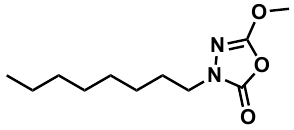

5-Methoxy-3-octyl-1,3,4-oxadiazol-2(3*H*)-one (**6**)

To a solution of carboxylate **S2** (100 mg, 0.494 mmol) in CH_2_Cl_2_ (9.90 mL), pyridine (119 µL, 1.48 mmol) was added. The resulting solution was cooled to 0 °C and followed by the addition of triphosgene (1.0 M in CH_2_Cl_2_, 494 µL) dropwise at 0 °C. The mixture was allowed to warm to ambient temperature. After being stirred for 16 h at the same temperature, the reaction mixture was quenched by saturated NH_4_Cl aqueous solution and then extracted with CH_2_Cl_2_. The organic layer was dried over MgSO_4_ and concentrated in vacuo. The residue was purified by flash column chromatography on silica gel (EtOAc/*n*-hexane = 1:9) to afford 7.5 mg (7%) of **6** as a colorless oil: ^1^H NMR (500 MHz, CDCl_3_) δ 3.98 (s, 3H), 3.61 (t, *J* = 7.2 Hz, 2H), 1.71 (quint, *J* = 7.3 Hz, 2H), 1.34 – 1.24 (m, 10H), 0.88 (t, *J* = 7.0 Hz, 3H); ^13^C NMR (126 MHz, CDCl_3_) δ 155.6, 151.6, 57.4, 45.9, 31.9, 29.3, 29.2, 28.1, 26.5, 22.8, 14.2; HRMS (ESI^+^) *m*/*z*: [M + H]^+^ calcd for C_11_H_21_N_2_O_3_ 229.1547; found 229.1544.

**Synthesis of JJ-OX-002 (7) and JJ-OX-003 (8)**

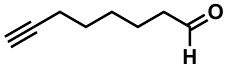

Oct-7-ynal (**S3**)

To a stirred solution of 7-octyn-1-ol (147 µL, 1.00 mmol) in CH_2_Cl_2_ (7.00 mL) was added Et_3_N (554 µL, 4.00 mmol) at 0 °C. To the reaction mixture, a solution of SO_3_·py (477 mg, 3.00 mmol) in DMSO (3.00 mL) was added at the same temperature. After being stirred for 3 h, the reaction flask was allowed to warm to ambient temperature and stirred another 1 h. Then, the reaction mixture was quenched by saturated NH_4_Cl aqueous solution, diluted with EtOAc, washed with H_2_O for 3 times, dried over MgSO_4_, and concentrated in vacuo. The residue was purified by flash column chromatography on silica gel (EtOAc/*n*-hexane = 1:8) to afford 102 mg (82%) of **S3** as a colorless oil: ^1^H NMR (500 MHz, CDCl_3_) δ 9.76 (t, *J* = 1.8 Hz, 1H), 2.43 (td, *J* = 7.3, 1.8 Hz, 2H), 2.19 (td, *J* = 7.0, 2.7 Hz, 2H), 1.93 (t, *J* = 2.6 Hz, 1H), 1.67 – 1.61 (m, 2H), 1.57 – 1.51 (m, 2H), 1.47 – 1.40 (m, 2H); ^13^C NMR (126 MHz, CDCl_3_) δ 202.6, 84.3, 68.6, 43.9, 28.3, 28.3, 21.7, 18.3. Spectroscopic data were consistent with literature reports.^19, 20^

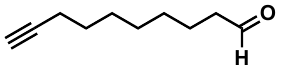

Dec-9-ynal (**S4**)

Dec-9-yn-1-ol (463 mg, 3.00 mmol) afforded 344 mg (75%) of **S4** as a colorless oil by following the same procedure as described above for compound **S3**. **S4** was purified by flash column chromatography on silica gel (EtOAc/*n*-hexane = 1:9): ^1^H NMR (500 MHz, CDCl_3_) δ 9.76 (t, *J* = 1.8 Hz, 1H), 2.42 (td, *J* = 7.3, 1.8 Hz, 2H), 2.18 (td, *J* = 7.0, 2.6 Hz, 2H), 1.94 (t, *J* = 2.6 Hz, 1H), 1.66 – 1.60 (m, 2H), 1.55 – 1.49 (m, 2H), 1.43 – 1.38 (m, 2H), 1.35 – 1.32 (m, 4H); ^13^C NMR (126 MHz, CDCl_3_) δ 203.0, 84.8, 68.3, 44.0, 29.2, 29.0, 28.6, 28.5, 22.1, 18.5. Spectroscopic data were consistent with literature reports.^19, 21^

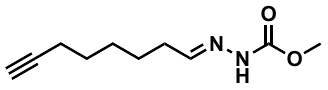

Methyl (*E*)-2-(oct-7-yn-1-ylidene)hydrazine-1-carboxylate (**S5**)

To a stirred solution of **S3** (69.6 mg, 0.560 mmol) in MeOH (1.00 mL), methyl carbazate (50.4 mg, 0.560 mmol) and AcOH (3.20 µL, 56.0 µmol) were added at ambient temperature. After being refluxed for 12 h, the reaction mixture was cooled and then concentrated in vacuo. The residue was recrystallized from MeOH/*n*-hexane to afford 93.9 mg (85%) of **S5** as a white solid. Obtained **S5** was directly used for the next step.

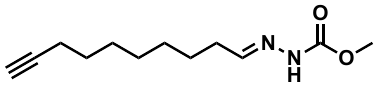

Methyl (*E*)-2-(dec-9-yn-1-ylidene)hydrazine-1-carboxylate (**S6**)

**S4** (344 mg, 2.26 mmol) afforded 342 mg (67%) of **S6** as a colorless oil by following the same procedures as described above for compound **S5**. **S6** was recrystallized from MeOH/ *n*-hexane. Obtained **S6** was directly used for the next step.

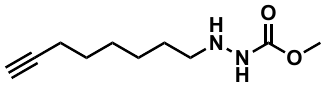

Methyl 2-(oct-7-yn-1-yl)hydrazine-1-carboxylate (**S7**)

To a stirred solution of **S5** (93.9 mg, 0.478 mmol) in MeOH (1.00 mL) was added NaBH_3_CN (75.4 mg, 1.20 mmol) and HCl (3 M in MeOH, 0.70 mL) at 0 ºC. After being stirred for 4 h at the same temperature, the reaction mixture was quenched with saturated NaHCO_3_ until basic pH and extracted with EtOAc. The organic layer was dried over MgSO_4_ and concentrated in vacuo. The resulting residue was purified by flash column chromatography on silica gel (EtOAc/*n*-hexane = 1:1) to afford 70.2 mg (74%) of **S7** as a colorless oil: ^1^H NMR (500 MHz, CDCl_3_) δ 3.71 (s, 3H), 2.86 (t, *J* = 7.3 Hz, 2H), 2.18 (td, *J* = 7.0, 2.6 Hz, 2H), 1.93 (t, *J* = 2.6 Hz, 1H), 1.55 – 1.32 (m, 8H); ^13^C NMR (126 MHz, CDCl_3_) δ 158.0, 84.7, 68.3, 52.6, 52.1, 28.7, 28.5, 27.6, 26.6, 18.5.

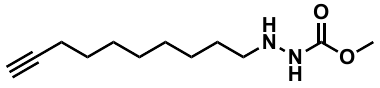

Methyl 2-(dec-9-yn-1-yl)hydrazine-1-carboxylate (**S8**)

**S6** (342 mg, 1.52 mmol) afforded 179 mg (52%) of **S8** as a colorless oil by following the same procedures as described above for compound **S7**. **S8** was purified by flash column chromatography on silica gel (EtOAc/*n*-hexane = 1:2): ^1^H NMR (500 MHz, CDCl_3_) δ 3.71 (s, 3H), 2.85 (t, *J* = 7.3 Hz, 2H), 2.17 (td, *J* = 7.1, 2.6 Hz, 2H), 1.93 (t, *J* = 2.6 Hz, 1H), 1.54 – 1.43 (m, 4H), 1.41 – 1.27 (m, 8H); ^13^C NMR (126 MHz, CDCl_3_) δ 157.9, 84.9, 68.2, 52.6, 52.1, 29.4, 29.1, 28.8, 28.6, 27.6, 27.1, 18.5.

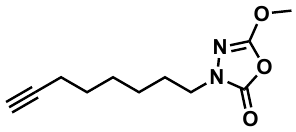

5-Methoxy-3-(oct-7-yn-1-yl)-1,3,4-oxadiazol-2(3*H*)-one (**7**)

To a solution of carboxylate **S7** (70.2 mg, 0.354 mmol) in CH_2_Cl_2_ (7.00 mL), pyridine (85.6 µL, 1.06 mmol) was added. The resulting solution was cooled to 0 °C and followed by the addition of triphosgene (1.0 M in CH_2_Cl_2_, 354 µL) dropwise at 0 °C. The mixture was allowed to warm to ambient temperature. After being stirred for 16 h at the same temperature, the reaction mixture was quenched by saturated NH_4_Cl aqueous solution and then extracted with CH_2_Cl_2_. The organic layer was dried over MgSO_4_ and concentrated in vacuo. The residue was purified by flash column chromatography on silica gel (EtOAc/*n*-hexane = 1:8) to afford 1.5 mg (2%) of **7** as a colorless oil: ^1^H NMR (500 MHz, CDCl_3_) δ 3.98 (s, 3H), 3.62 (t, *J* = 7.1 Hz, 2H), 2.19 (td, *J* = 7.0, 2.7 Hz, 2H), 1.94 (t, *J* = 2.6 Hz, 1H), 1.73 (q, *J* = 7.4 Hz, 2H), 1.56 – 1.51 (m, 2H), 1.48 – 1.42 (m, 2H), 1.39 – 1.32 (m, 2H); ^13^C NMR (126 MHz, CDCl_3_) δ 155.7, 151.6, 84.6, 68.5, 57.4, 45.8, 28.4, 28.3, 27.9, 26.0, 18.4; HRMS (ESI^+^) *m*/*z*: [M + H]^+^ calcd for C_11_H_17_N_2_O_3_ 225.1234; found 225.1232.

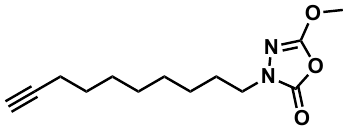

3-(Dec-9-yn-1-yl)-5-methoxy-1,3,4-oxadiazol-2(3*H*)-one (**8**)

**S8** (179 mg, 0.791 mmol) afforded 7.8 mg (4%) of **8** as a colorless oil by following the same procedures as described above for compound **7**. **8** was purified by flash column chromatography on silica gel (EtOAc/*n*-hexane = 1:8): ^1^H NMR (500 MHz, CDCl_3_) δ 3.98 (s, 3H), 3.61 (t, *J* = 7.1 Hz, 2H), 2.18 (td, *J* = 7.1, 2.7 Hz, 2H), 1.93 (t, *J* = 2.6 Hz, 1H), 1.73 – 1.68 (m, 2H), 1.55 – 1.49 (m, 2H), 1.42 – 1.30 (m, 8H); HRMS (ESI^+^) *m*/*z*: [M + H]^+^ calcd for C_13_H_21_N_2_O_3_ 253.1547; found 253.1546.

**Synthesis of JJ-OX-004 (9) and JJ-OX-007 (10)**

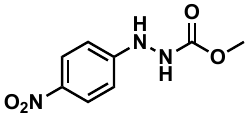

Methyl 2-(4-nitrophenyl)hydrazine-1-carboxylate (**S9**)

To a solution of 4-nitrophenylhydrazine hydrochloride (3.80 g, 20.0 mmol) in pyridine (50.0 mL), *N*-methyl-2-pyrrolidone (1.73 mL, 18.0 mmol) was added. The resulting solution was cooled to 0 °C and followed by the addition of methyl chloroformate (1.63 mL, 21.0 mmol) dropwise at 0 °C. After being stirred for 16 h at the same temperature, the reaction mixture was resuspended in EtOAc (100 mL), quenched with 1 M aqueous HCl, and then extracted with EtOAc. The combined organic layers were dried over MgSO_4_ and concentrated in vacuo. The residue was filtered and roughly washed with *n*-hexane. The resulting crude product was used for the next step without any further purification.

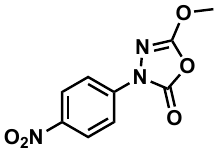

5-Methoxy-3-(4-nitrophenyl)-1,3,4-oxadiazol-2(3*H*)-one (**S10**)

To a solution of crude **S9** (4.09 g, 19.4 mmol) in CH_2_Cl_2_ (388 mL), pyridine (3.13 mL, 38.8 mmol) was added. The resulting solution was cooled to 0 °C and followed by the addition of triphosgene (19.4 mL of 1.0 M solution in CH_2_Cl_2_) dropwise at 0 °C. The mixture was allowed to warm to ambient temperature. After being stirred for 3 h at the same temperature, the reaction mixture was quenched by sat. NH_4_Cl aqueous solution and then extracted with CH_2_Cl_2_. The combined organic layers were dried over MgSO_4_ and concentrated in vacuo. The residue was purified by flash column chromatography on silica gel (EtOAc/CH_2_Cl_2_/*n*-hexane = 1:1:5) to afford 1.60 g (17% for two steps) of **S10** as a pale-yellow solid: ^1^H NMR (500 MHz, CDCl_3_) δ 8.32 – 8.28 (m, 2H), 8.02 – 7.99 (m, 2H), 4.16 (s, 3H); ^13^C NMR (126 MHz, CDCl_3_) δ 156.4, 148.0, 144.7, 141.2, 125.2, 117.7, 58.2. Spectroscopic data were consistent with literature reports.^2^

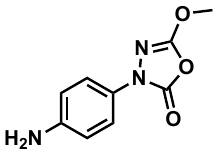

3-(4-Aminophenyl)-5-methoxy-1,3,4-oxadiazol-2(3*H*)-one (**S11**)

To a stirred solution of **S10** (1.60 g, 6.75 mmol) in CH_2_Cl_2_ (100 mL) and MeOH (100 mL) was added Pd/C (482 mg, 4.53 mmol). The reaction mixture was stirred for 3 h while hydrogen gas was bubbled through the mixture. The suspension was removed by celite filtration, and the filtrate was concentrated under the reduced pressure. The residue was diluted by EtOAc and washed with brine. The combined organic layers dried over MgSO_4_ and concentrated in vacuo. The resulting residue was purified by flash column chromatography on silica gel (EtOAc/*n*-hexane = 1:1) to afford 860 mg (61%) of **S11** as an orange solid: ^1^H NMR (500 MHz, CDCl_3_) δ 7.52 – 7.48 (m, 2H), 6.71 – 6.68 (m, 2H), 4.06 (s, 3H), 3.67 (brs, 2H); ^13^C NMR (126 MHz, CDCl_3_) δ 155.8, 148.7, 144.8, 127.6, 120.4, 115.3, 57.7. Spectroscopic data were consistent with literature reports.^2^

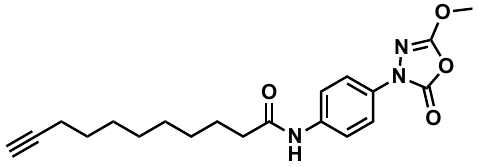

*N*-(4-(5-methoxy-2-oxo-1,3,4-oxadiazol-3(2*H*)-yl)phenyl)undec-10-ynamide (**9**)

To a stirred solution of undec-10-ynoic acid (17.3 mg, 95.1 µmol) in CH_2_Cl_2_ (2.00 mL), oxalyl chloride (24.5 µL, 286 µmol) and 2 drops of DMF were added at ambient temperature. After stirring for 1 h, the reaction mixture was concentrated under reduced pressure and then diluted with CH_2_Cl_2_ (1.50 mL). To the resulting solution, a solution of aniline **S11** (19.7 mg, 95.1 µmol) in CH_2_Cl_2_ (400 µL) and Et_3_N (8.4 µL, 105 µmol) were added at 0 °C. After stirring for 2 h, the reaction was quenched with H_2_O and then extracted with CH_2_Cl_2_. The organic layer was dried over MgSO_4_ and concentrated in vacuo. The resulting residue was purified by flash column chromatography on silica gel (EtOAc/*n*-hexane = 1:2) to afford 30.9 mg (87%) of **9** as a white solid: ^1^H NMR (500 MHz, CDCl_3_) δ 7.73 – 7.71 (m, 2H), 7.59 – 7.57 (m, 2H), 7.32 (s, 1H), 4.10 (s, 3H), 2.38 – 2.33 (m, 2H), 2.17 (td, *J* = 7.1, 2.7 Hz, 2H), 1.93 (t, *J* = 2.6 Hz, 1H), 1.72 (q, *J* = 7.4 Hz, 2H), 1.56 – 1.47 (m, 2H), 1.42 – 1.29 (m, 8H); ^13^C NMR (126 MHz, CDCl_3_) δ 171.6, 156.0, 148.4, 135.7, 132.3, 120.5, 118.8, 84.9, 68.3, 57.9, 37.8, 29.3, 29.3, 29.0, 28.8, 28.6, 25.6, 18.5; HRMS (ESI^+^) *m*/*z*: [M + H]^+^ calcd for C_20_H_26_N_3_O_4_ 372.1918; found 372.1917.

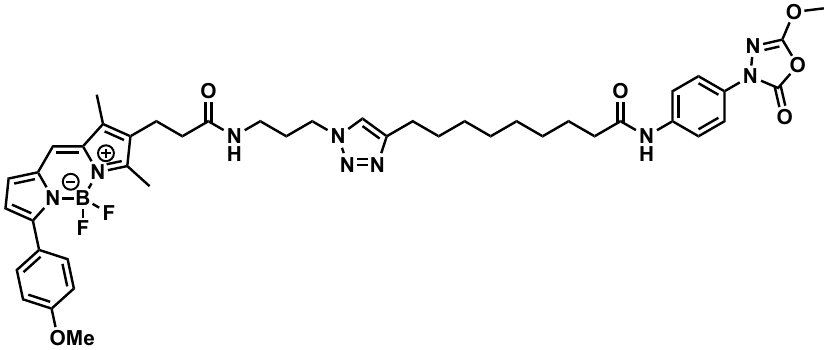

9-(1-(3-(3-(5,5-Difluoro-7-(4-methoxyphenyl)-1,3-dimethyl-5*H*-4λ^4^,5λ^4^-dipyrrolo[1,2-*c*:2',1'-*f*][1,3,2]diazaborinin-2-yl)propanamido)propyl)-1*H*-1,2,3-triazol-4-yl)-*N*-(4-(5-methoxy-2-oxo-1,3,4-oxadiazol-3(2*H*)-yl)phenyl)nonanamide (**10**)

To a stirred solution of alkyne **9** (4.7 mg, 12.6 µmol) in DMF (900 µL) was added BDP-TMR azide (5.0 mg, 10.4 µmol) at ambient temperature. To a stirred reaction mixture, 100 mM CuSO_4_·5H_2_O solution in DMF (25 µL, 2.5 µmol) and 200 mM ascorbic acid solution in DMF (62.5 µL, 12.5 µmol) were added. After stirring at the same temperature for 16 h, the mixture was diluted with H_2_O and extracted 3 times with EtOAc. The combined organic layer was dried over Na_2_SO_4_ and concentrated in vacuo. The residue was purified by flash column chromatography on silica gel (H_2_O/CH_3_CN/*n*-hexane = 8:1:1) to afford 3.7 mg (42%) of **10** as a magenta solid: ^1^H NMR (500 MHz, CDCl_3_) δ 8.44 (brs, 1H), 7.85 (d, *J* = 8.8 Hz, 2H), 7.71 – 7.62 (m, 5H), 7.06 (s, 1H), 6.99 – 6.94 (m, 3H), 6.54 (d, *J* = 4.1 Hz, 1H), 4.24 – 4.19 (m, 2H), 4.08 (s, 3H), 3.85 (s, 3H), 3.20 – 3.19 (m, 2H), 2.77 – 2.67 (m, 4H), 2.53 (s, 3H), 2.41 – 2.36 (m, 4H), 2.22 (s, 3H), 2.04 – 2.00 (m, 2H), 1.71 – 1.62 (m, 4H), 1.32 – 1.26 (m, 8H); HRMS (ESI^+^) *m*/*z*: [M + H]^+^ calcd for C_44_H_53_BF_2_N_9_O_6_ 852.4174; found 852.4177.

**4. LC traces of synthesized probes 7–10**

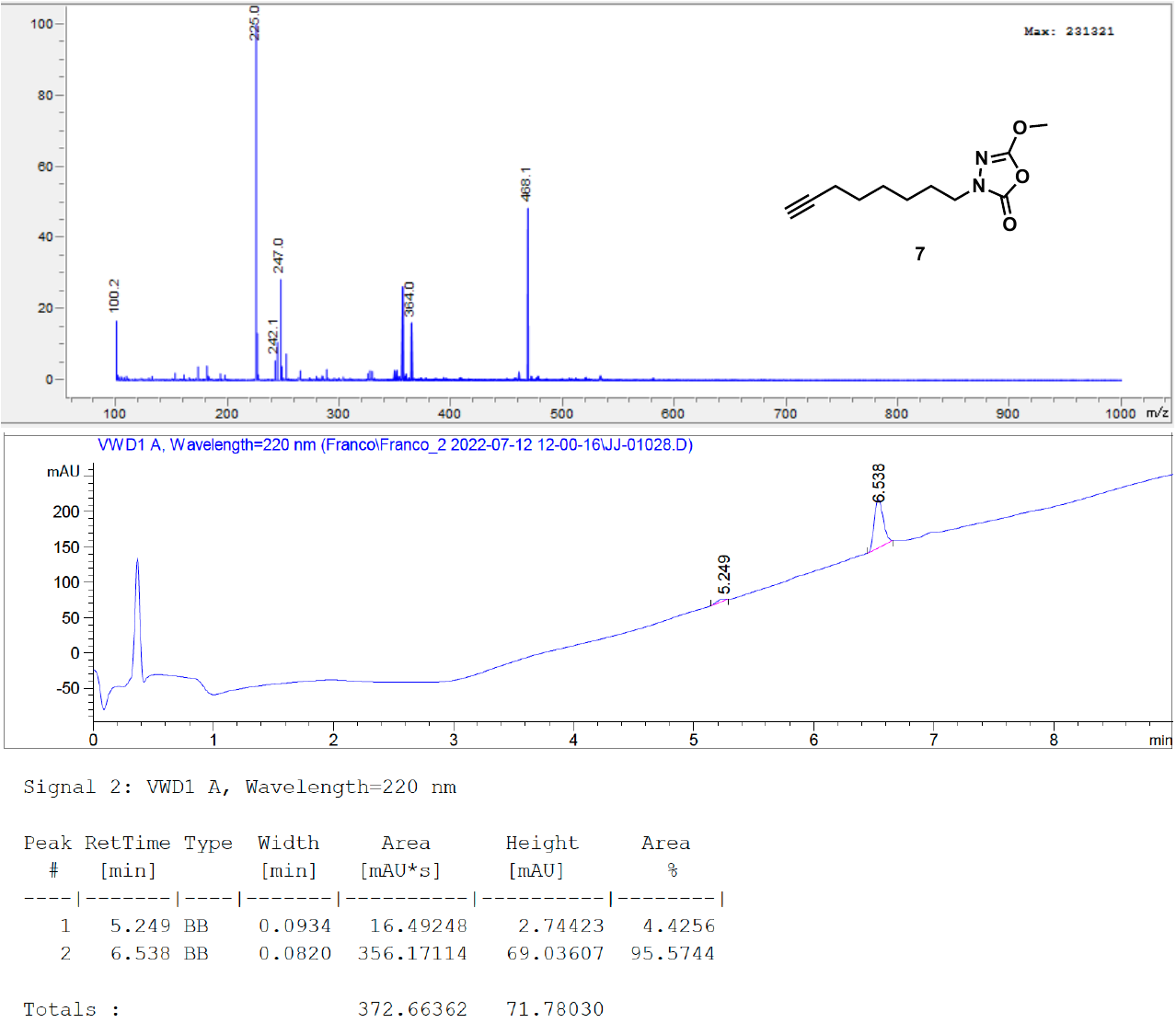

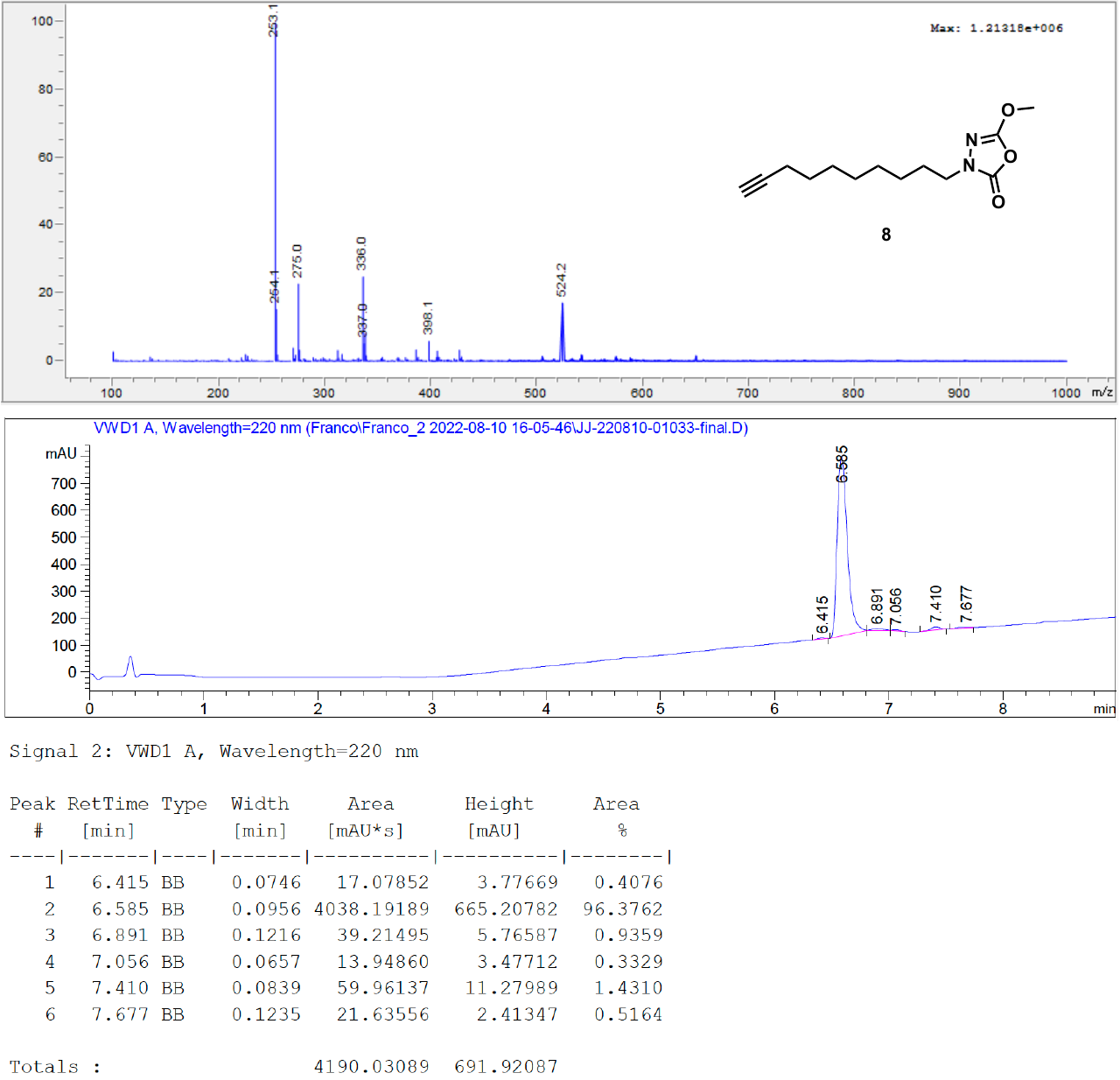

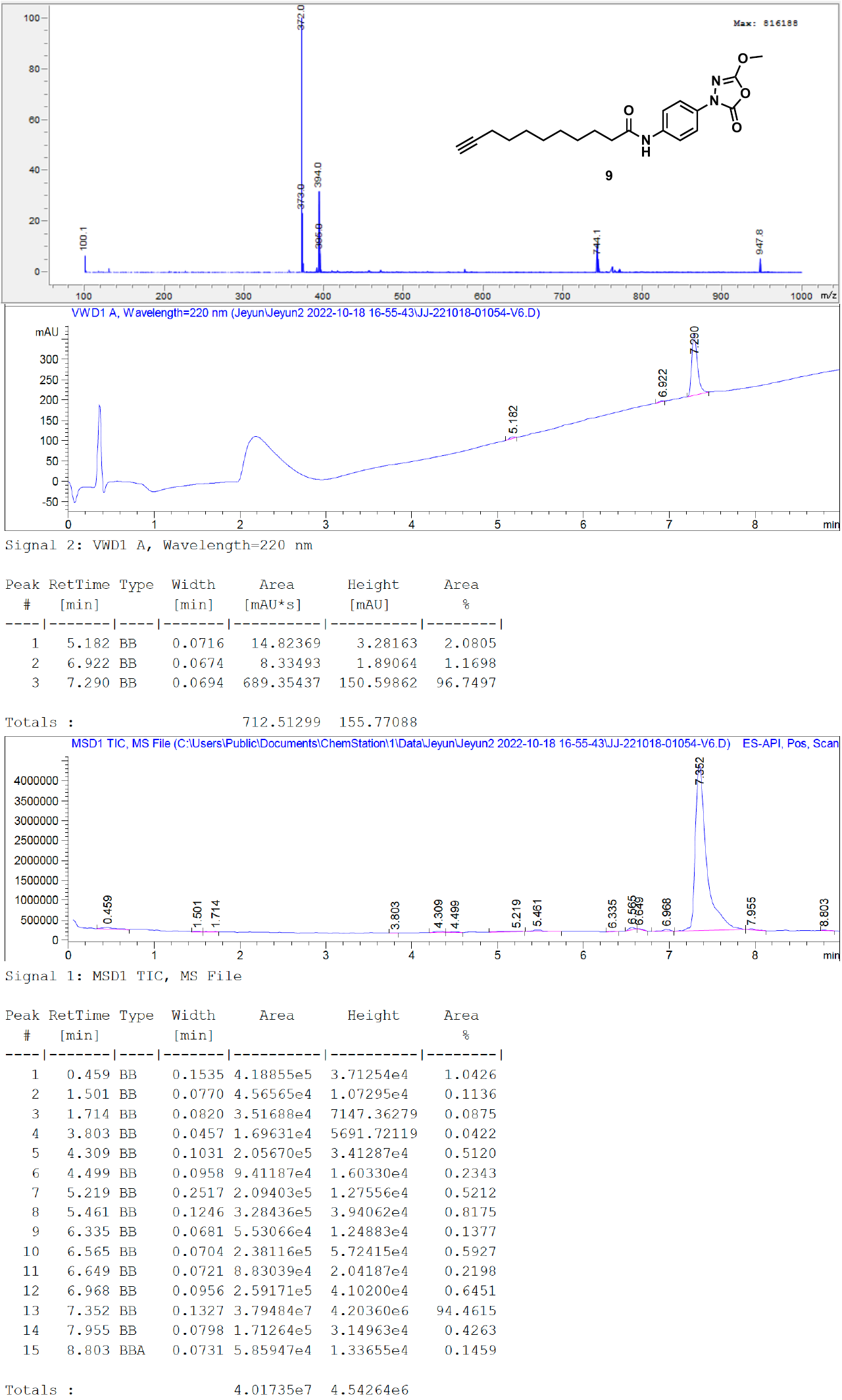

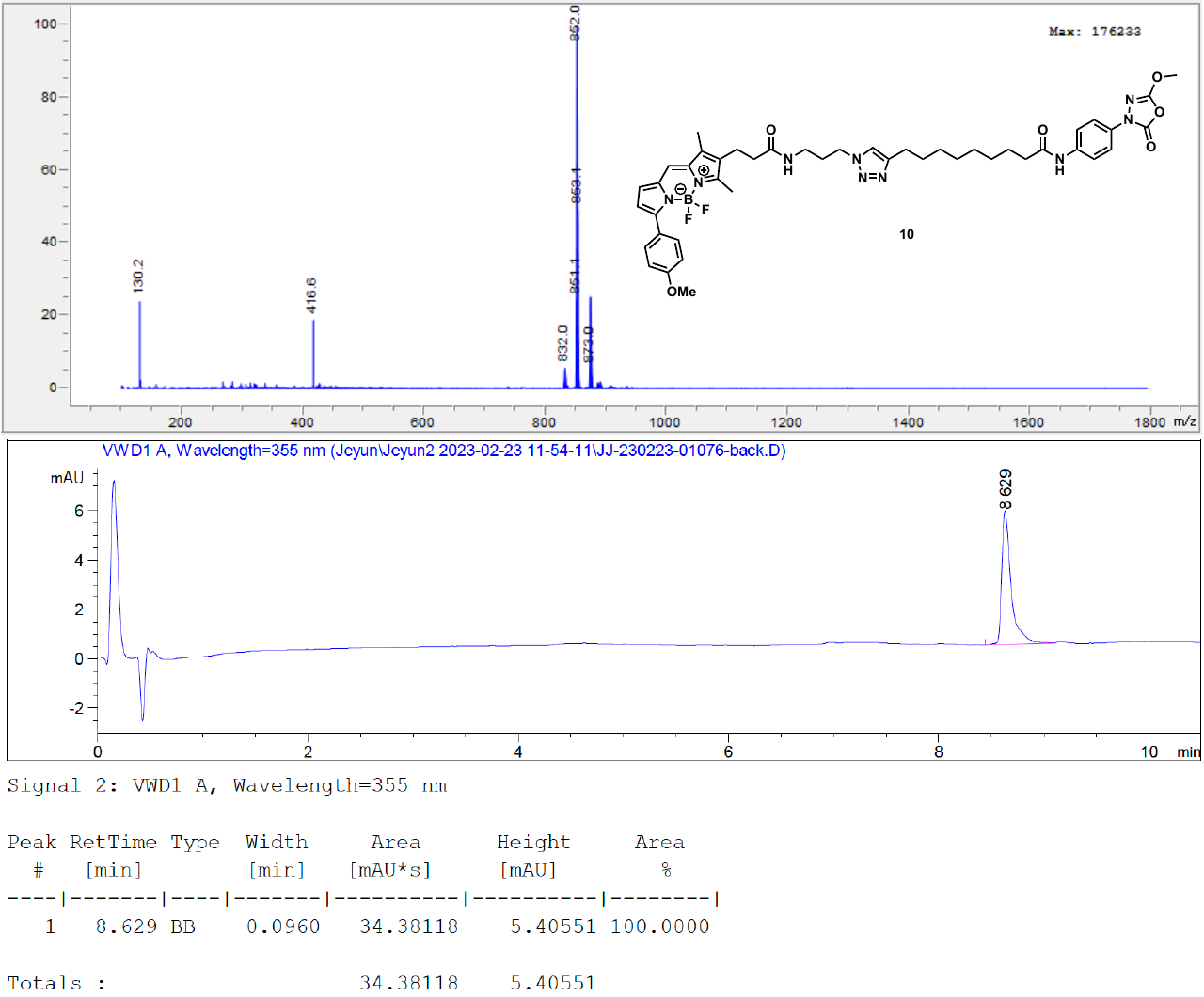

**5. ^1^H NMR Spectra**

• **^1^H NMR spectrum of compound S2 (CDCl_3_, 500 MHz)**
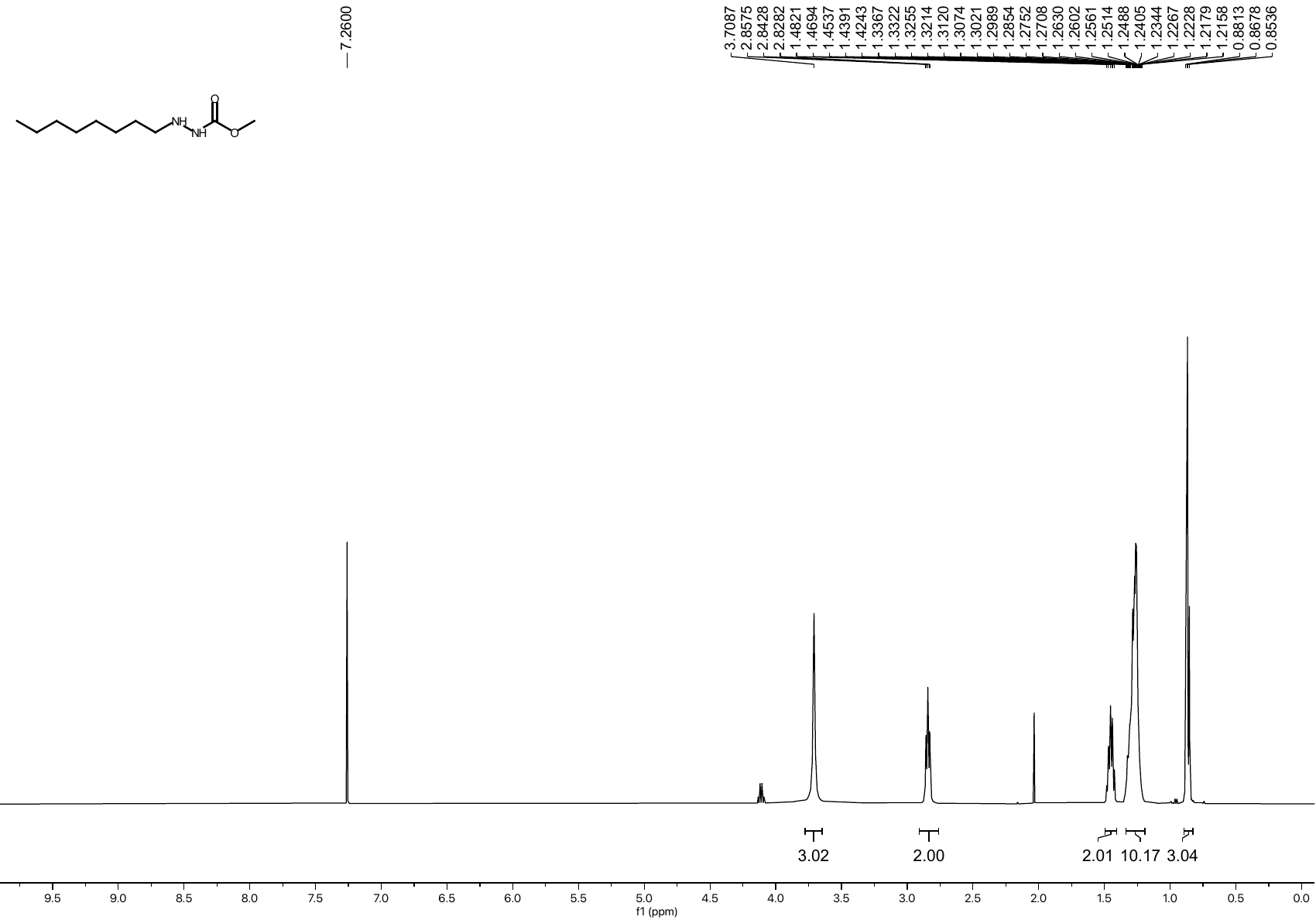

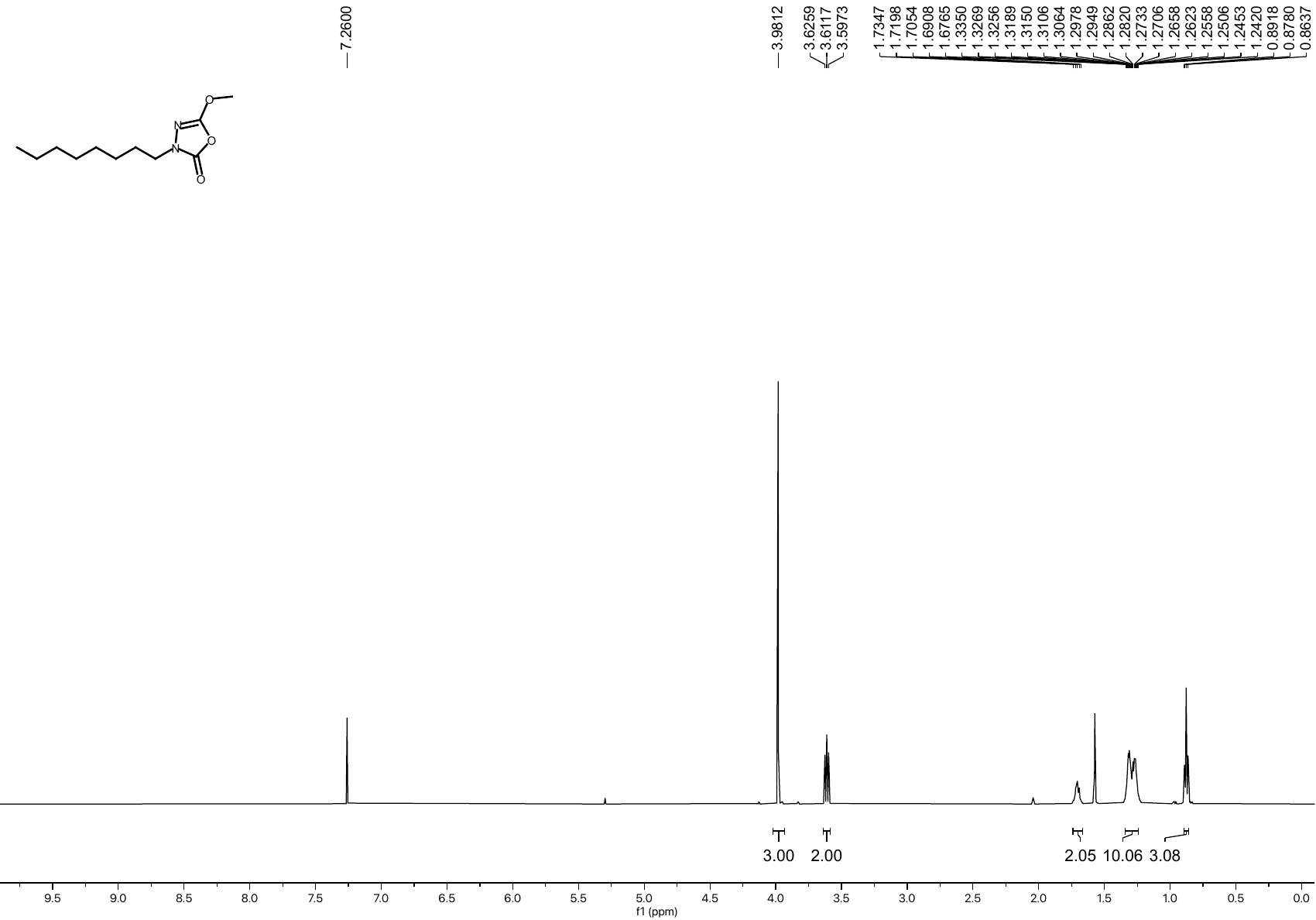
• **^1^H NMR spectrum of compound 6 (CDCl_3_, 500 MHz)**

• **^1^H NMR spectrum of compound S3 (CDCl_3_, 500 MHz)**
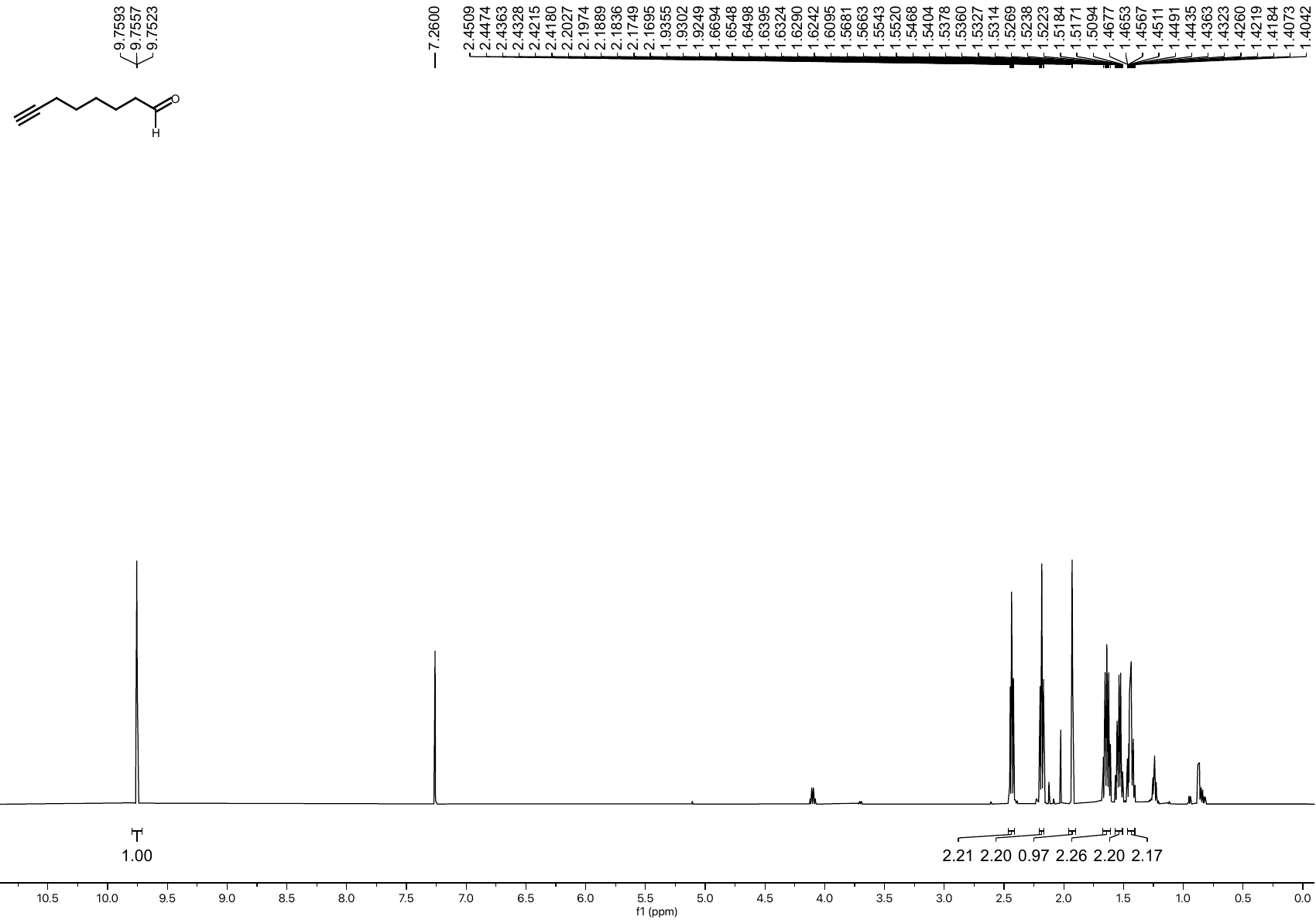

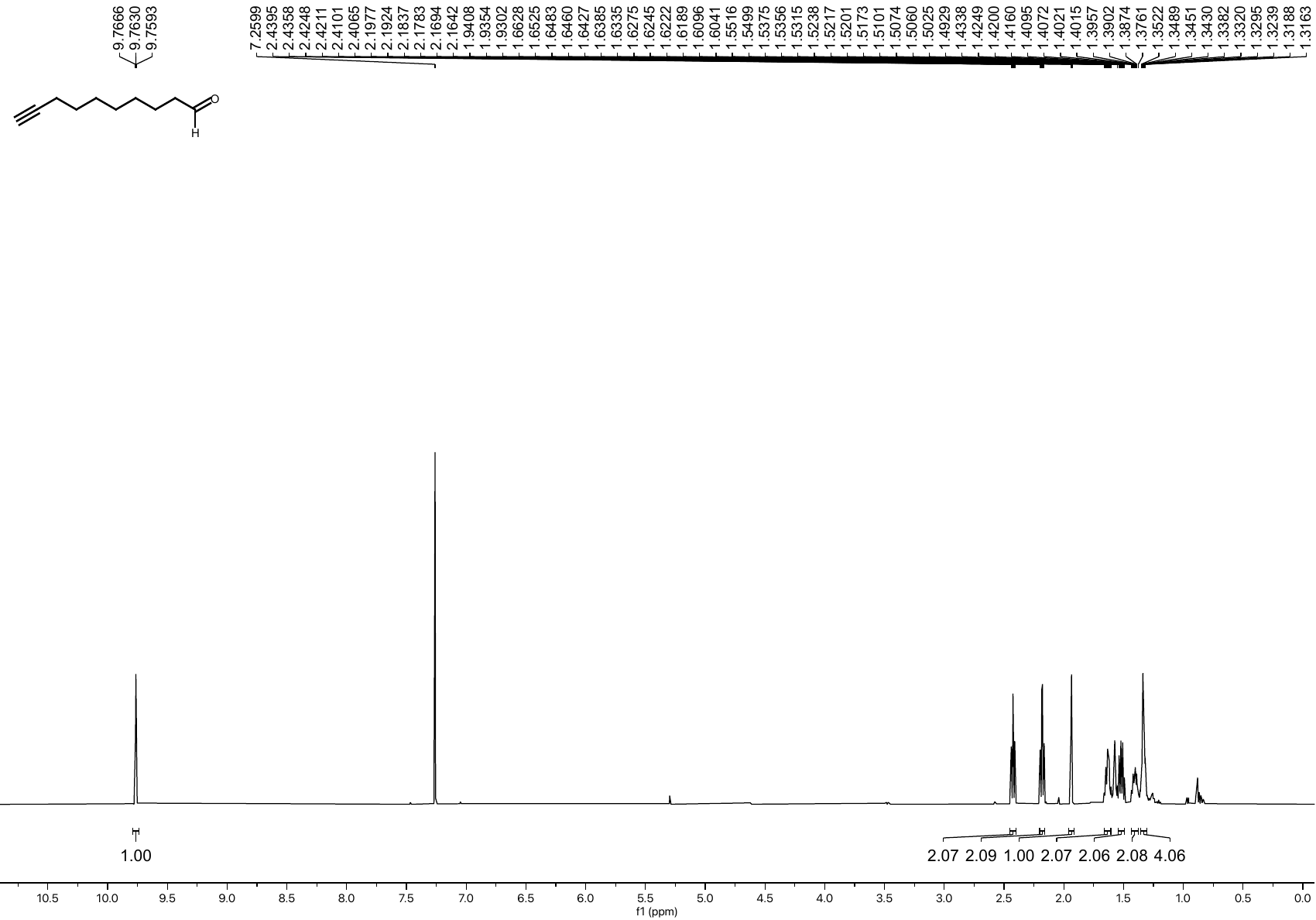
• **^1^H NMR spectrum of compound S4 (CDCl_3_, 500 MHz)**

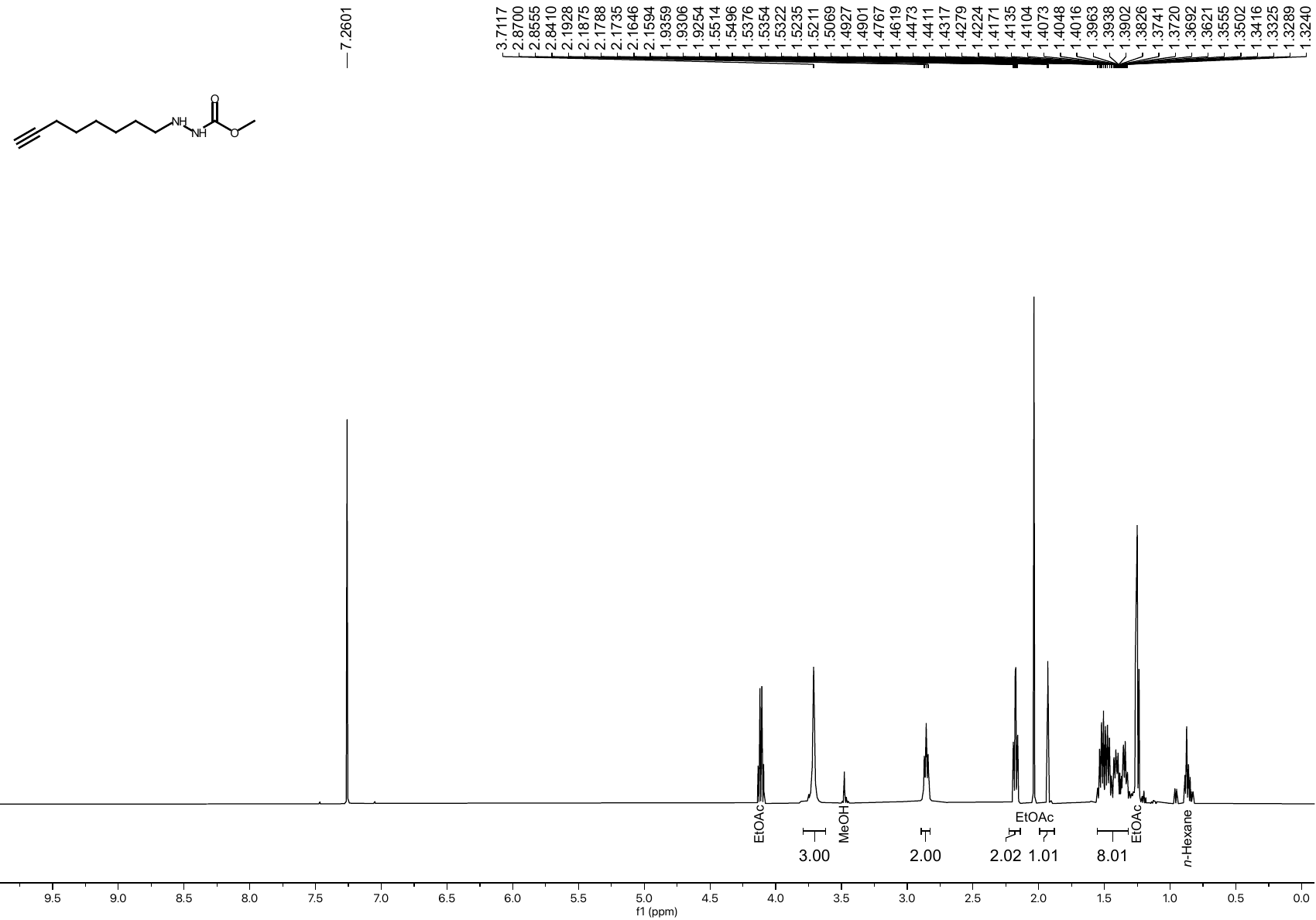
• **^1^H NMR spectrum of compound S7 (CDCl_3_, 500 MHz)**

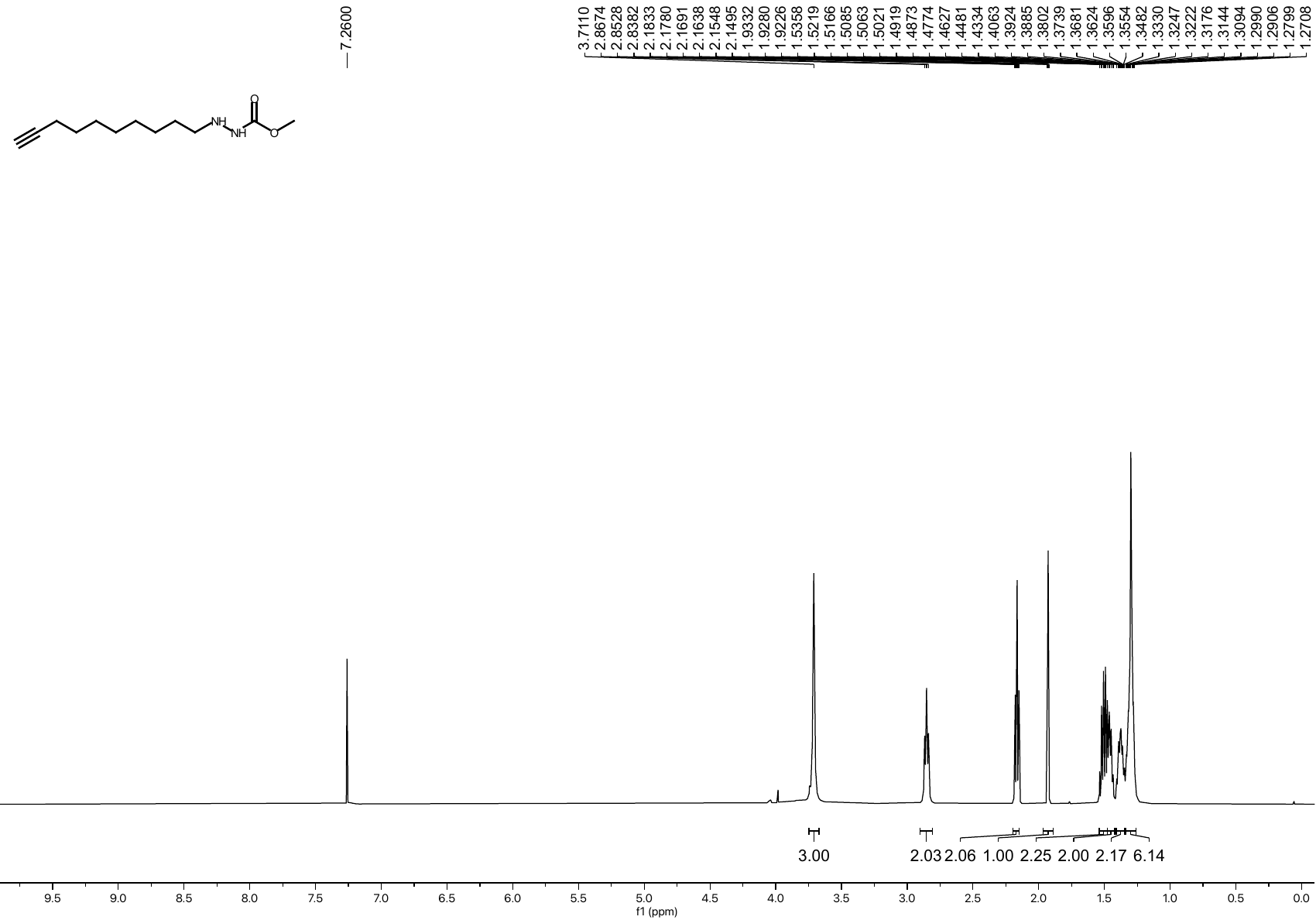
• **^1^H NMR spectrum of compound S8 (CDCl_3_, 500 MHz)**

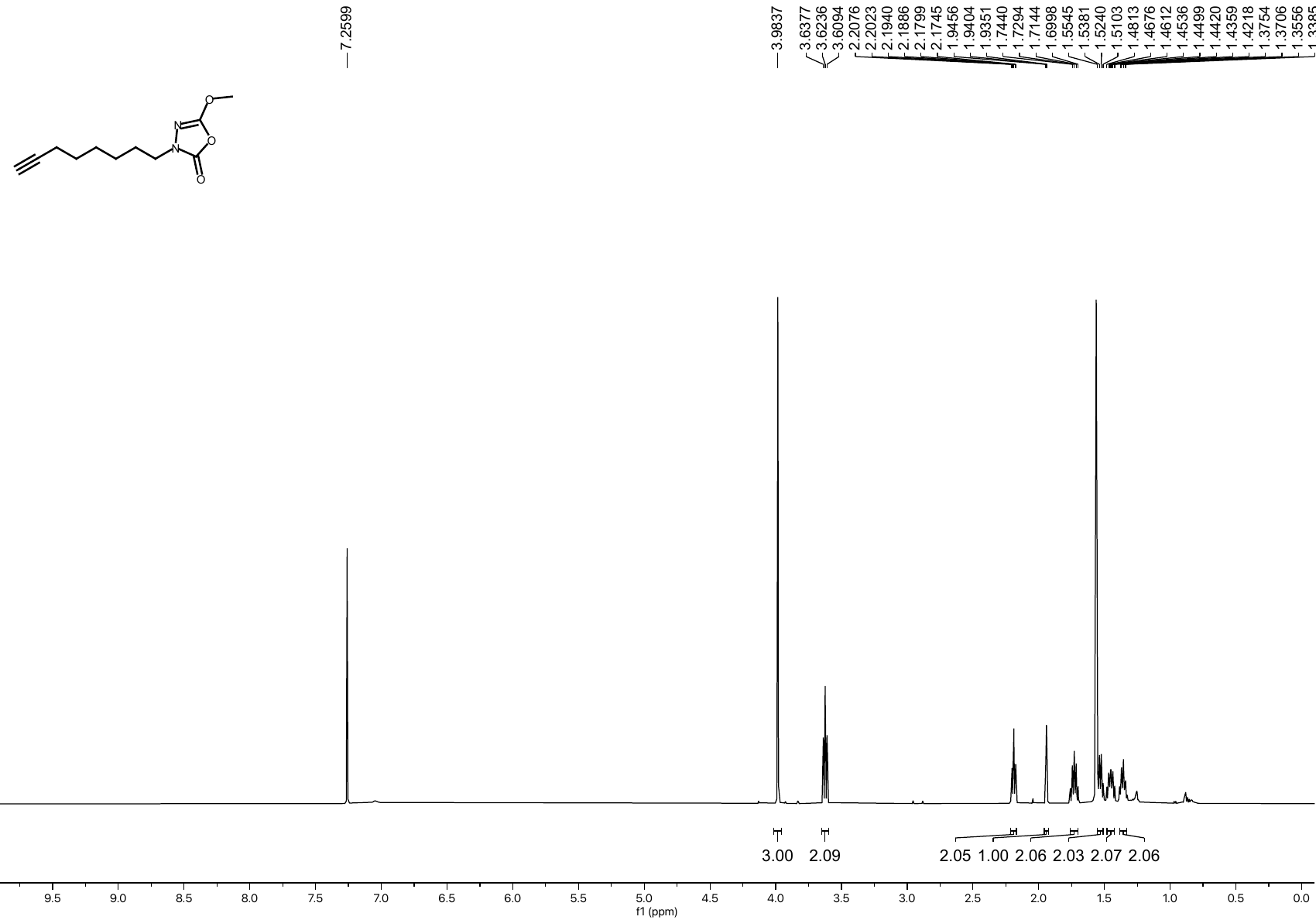
• **^1^H NMR spectrum of compound 7 (CDCl_3_, 500 MHz)**

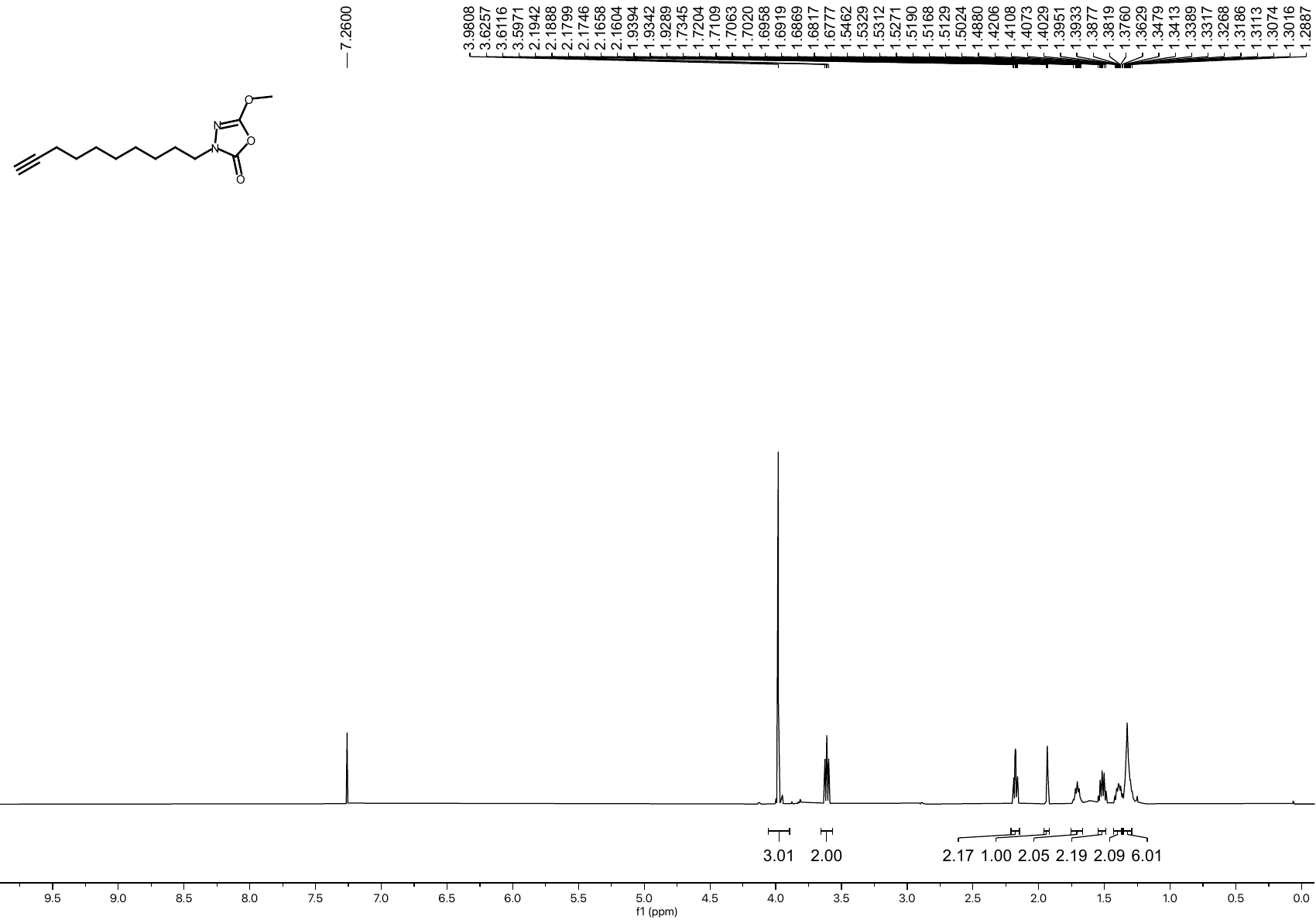
• **^1^H NMR spectrum of compound 8 (CDCl_3_, 500 MHz)**

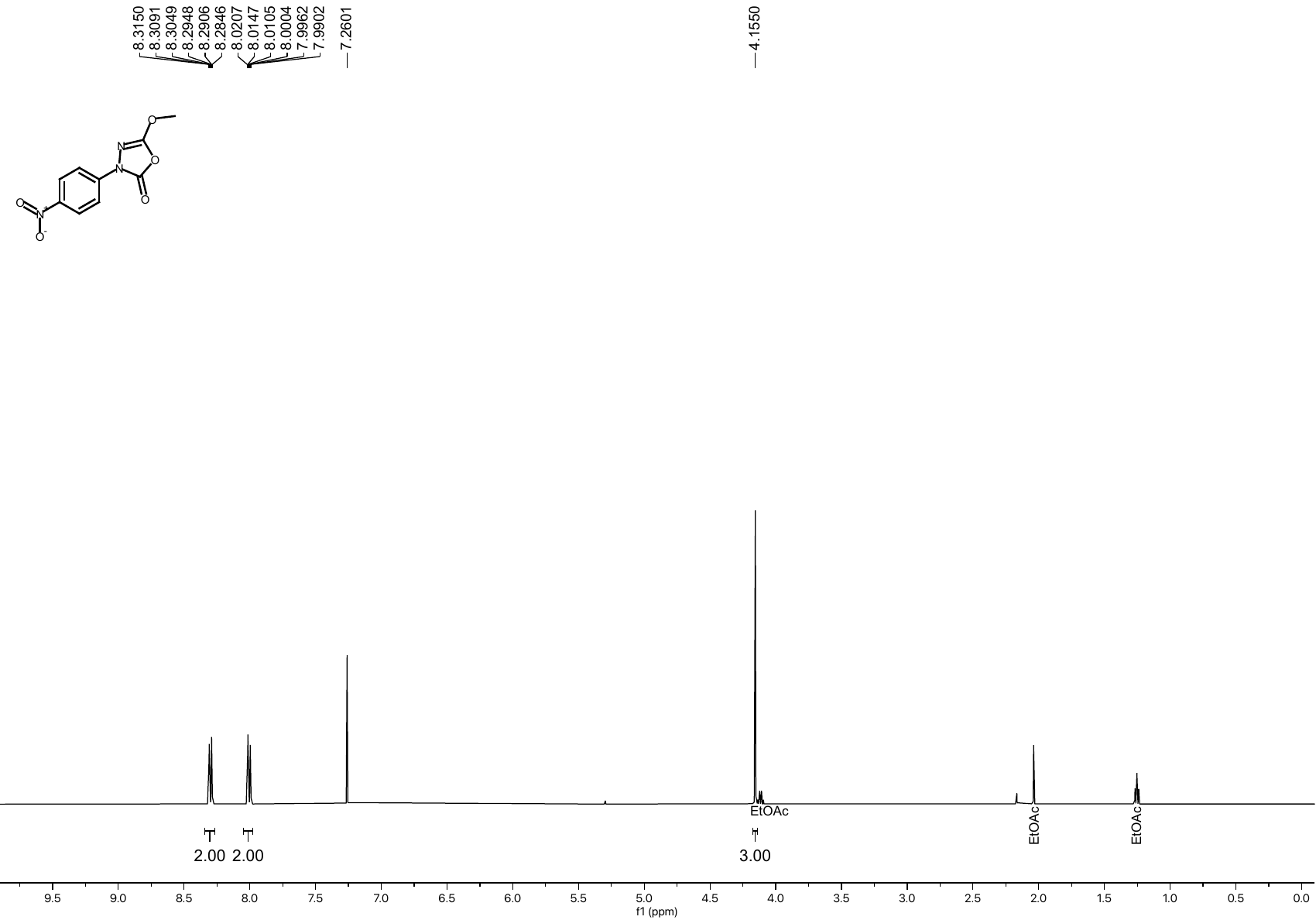
• **^1^H NMR spectrum of compound S10 (CDCl_3_, 500 MHz)**

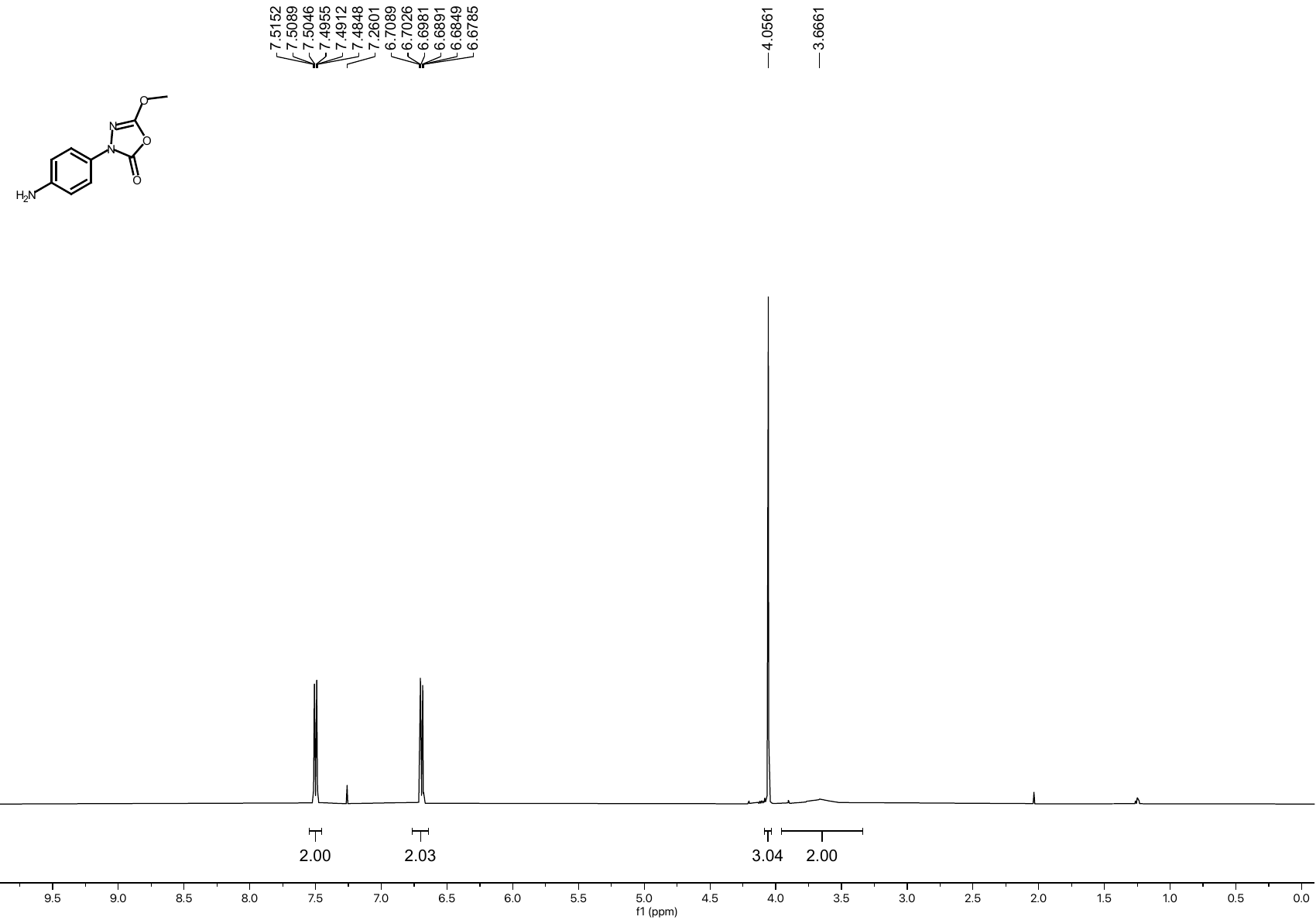
• **^1^H NMR spectrum of compound S11 (CDCl_3_, 500 MHz)**

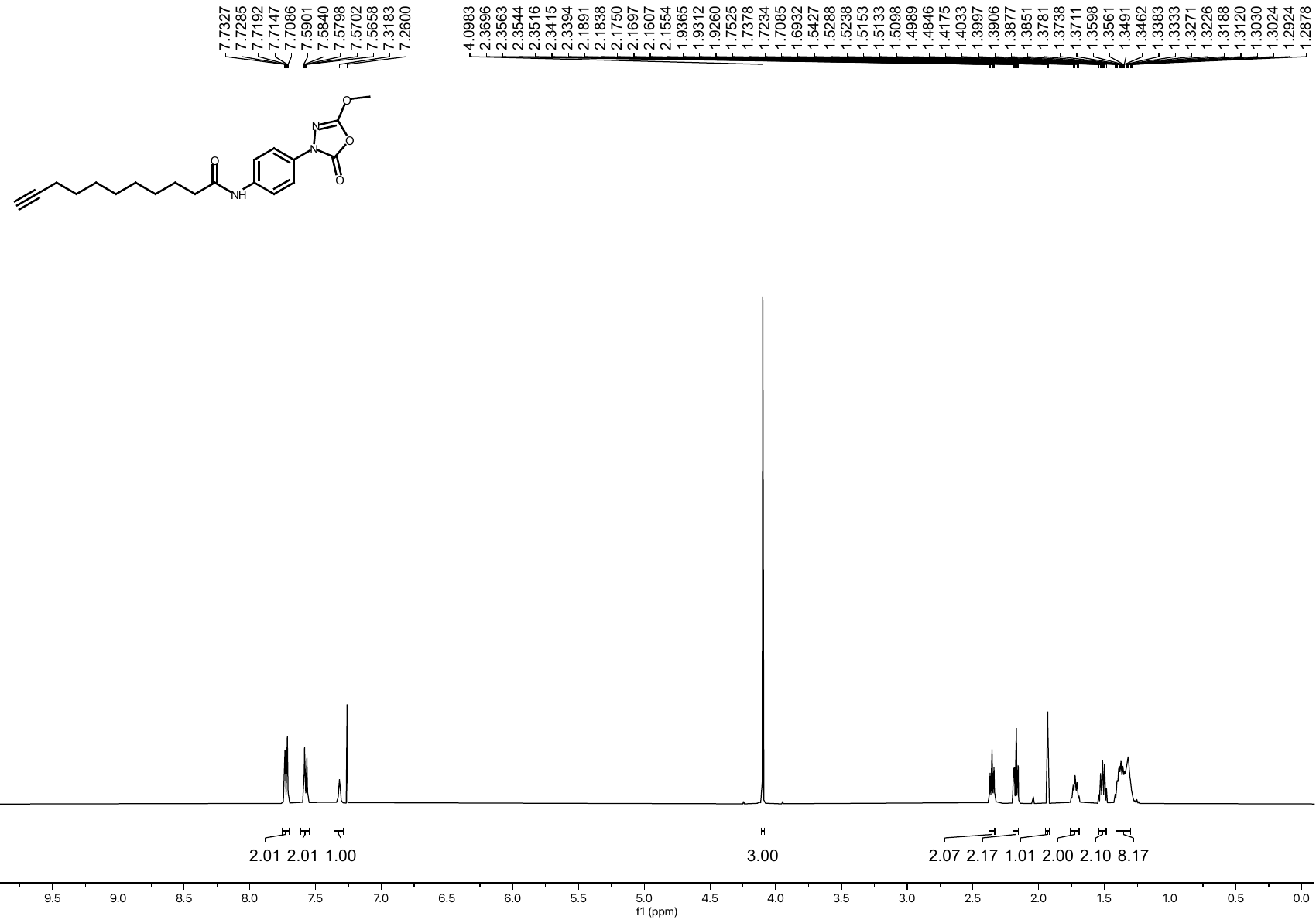
• **^1^H NMR spectrum of compound 9 (CDCl_3_, 500 MHz)**

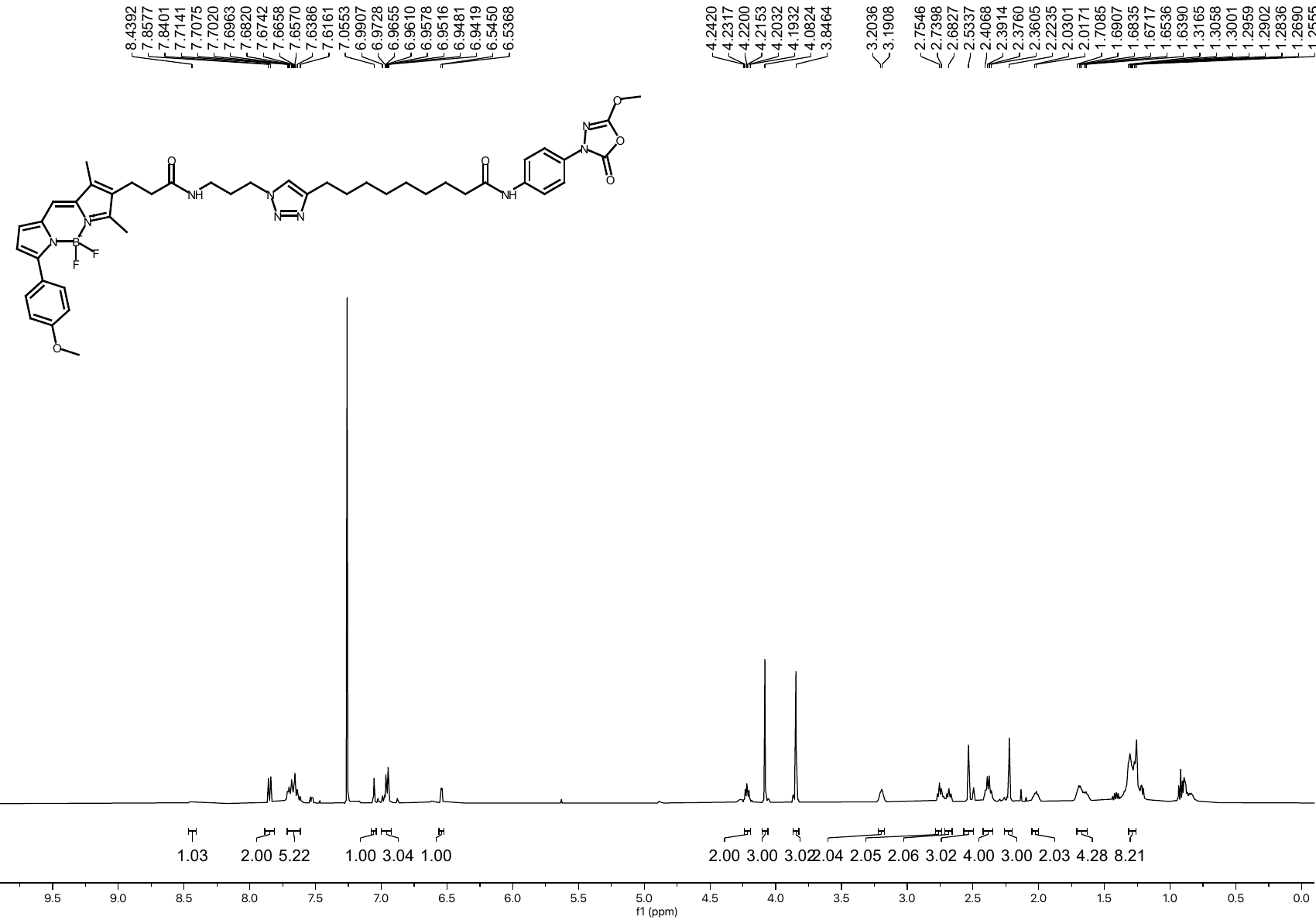
• **^1^H NMR spectrum of compound 10 (CDCl_3_, 500 MHz)**

**6. ^13^C NMR Spectra**

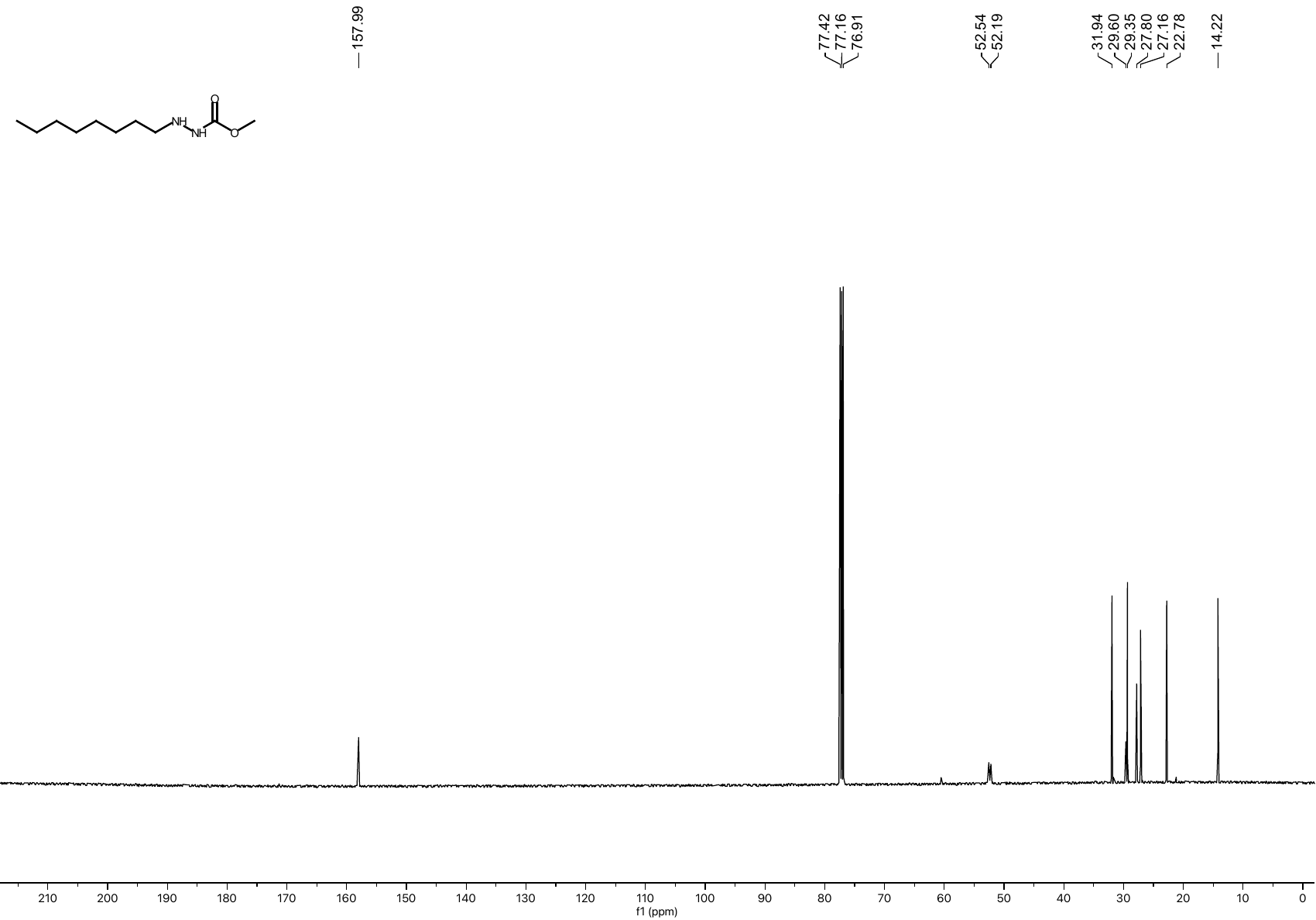
• **^13^C NMR spectrum of compound S2 (CDCl_3_, 126 MHz)**

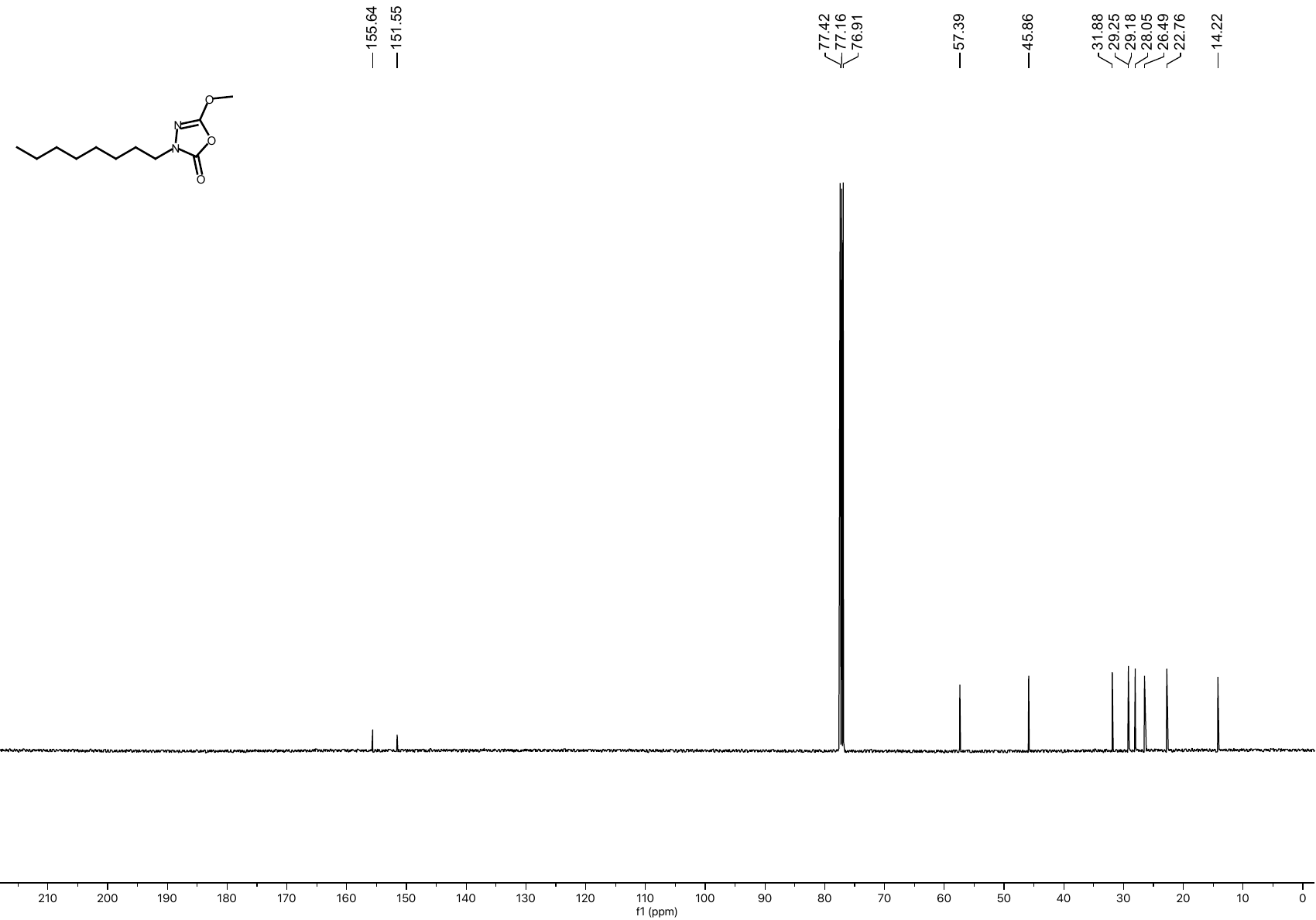
• **^13^C NMR spectrum of compound 6 (CDCl_3_, 126 MHz)**

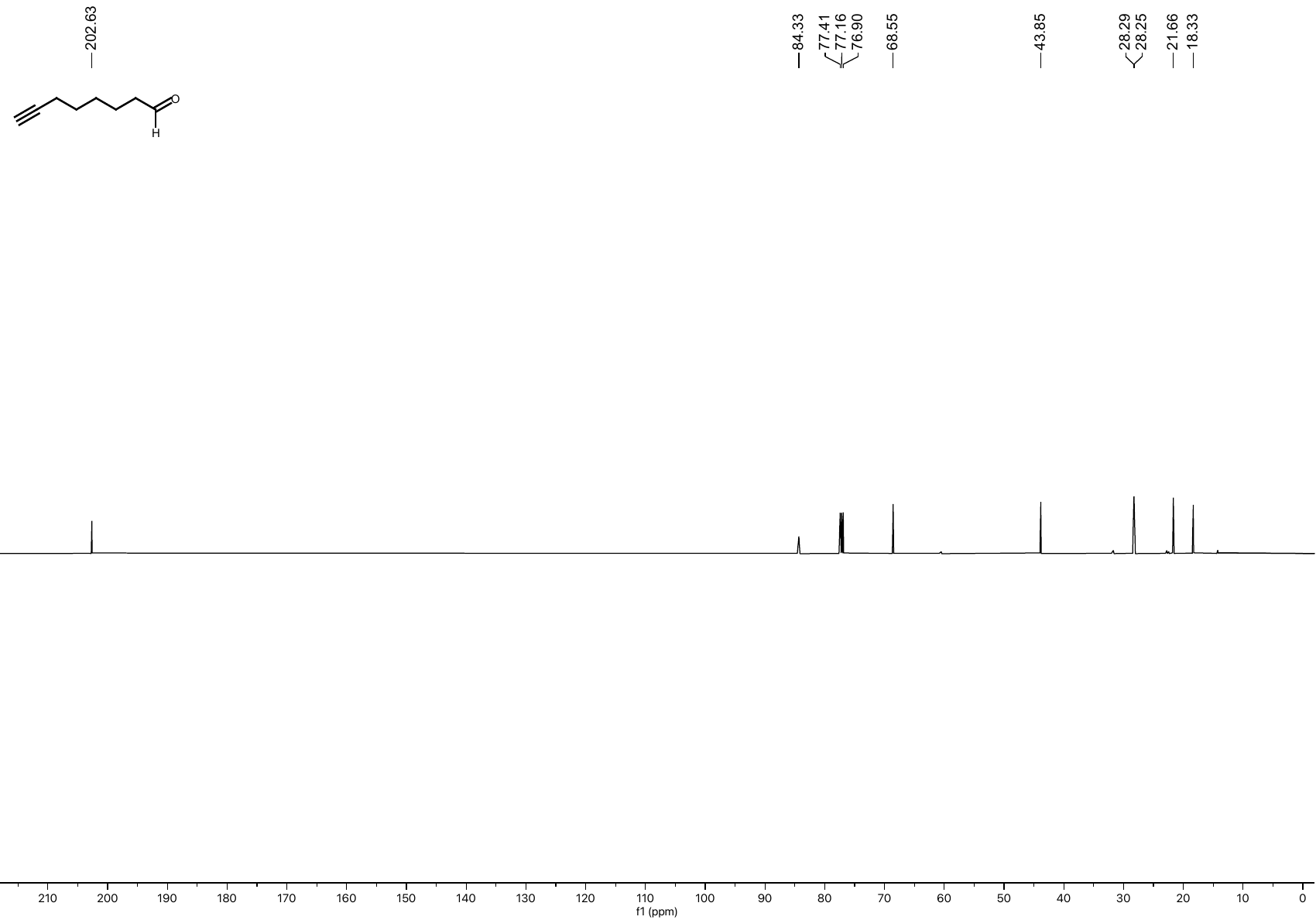
• **^13^C NMR spectrum of compound S3 (CDCl_3_, 126 MHz)**

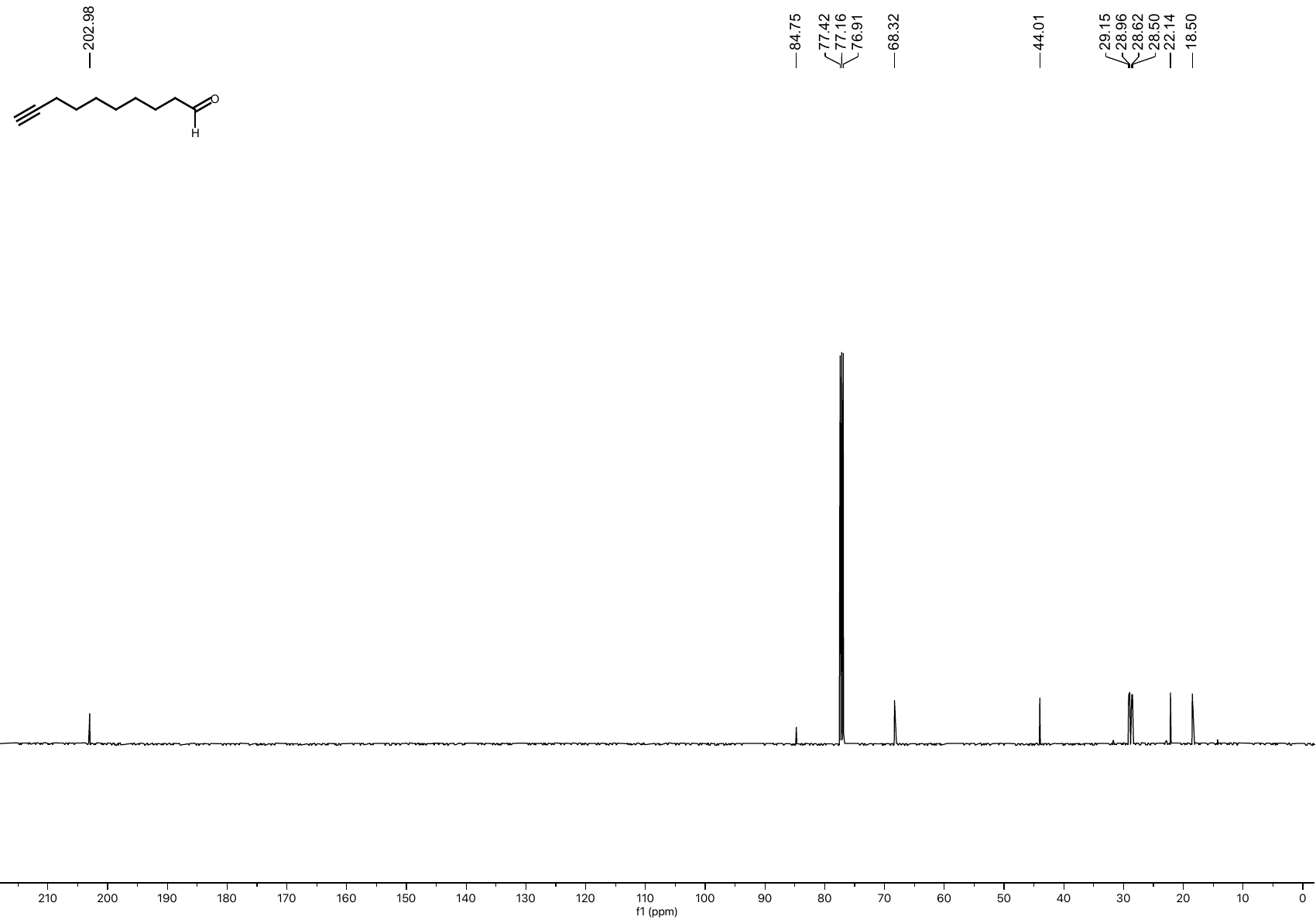
• **^13^C NMR spectrum of compound S4 (CDCl_3_, 126 MHz)**

• **^13^C NMR spectrum of compound S7 (CDCl_3_, 126 MHz)**
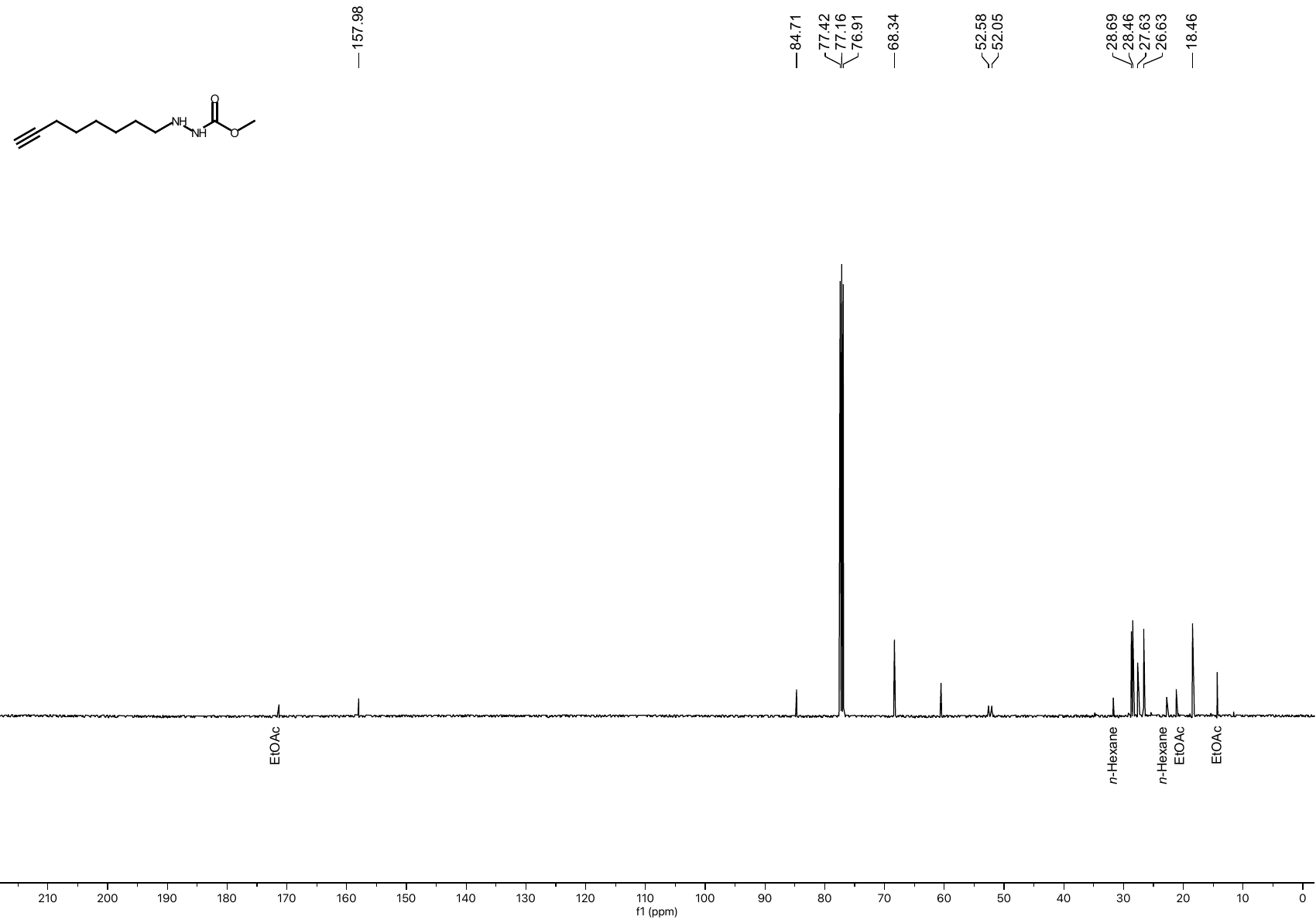

• **^13^C NMR spectrum of compound S8 (CDCl_3_, 126 MHz)**
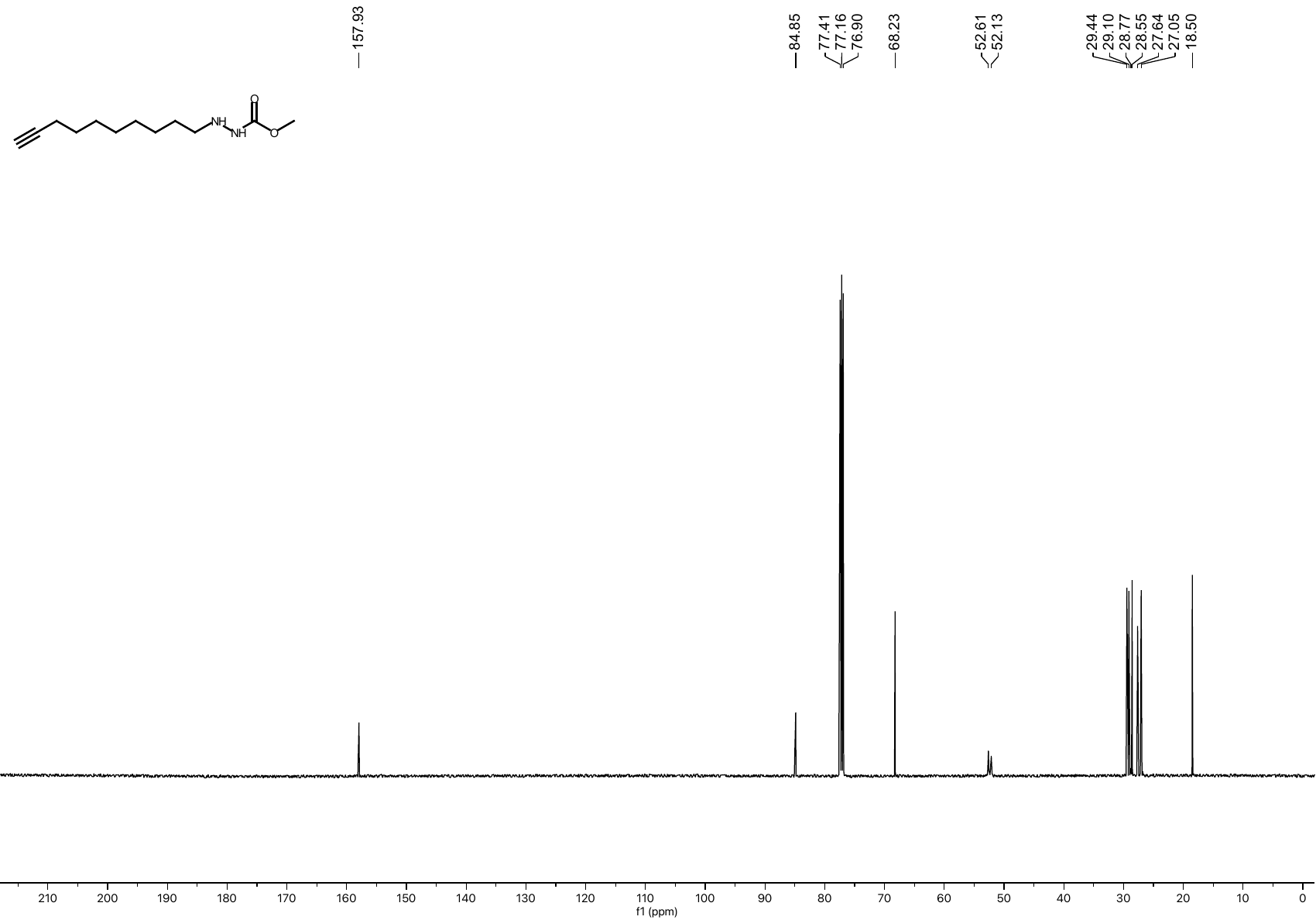

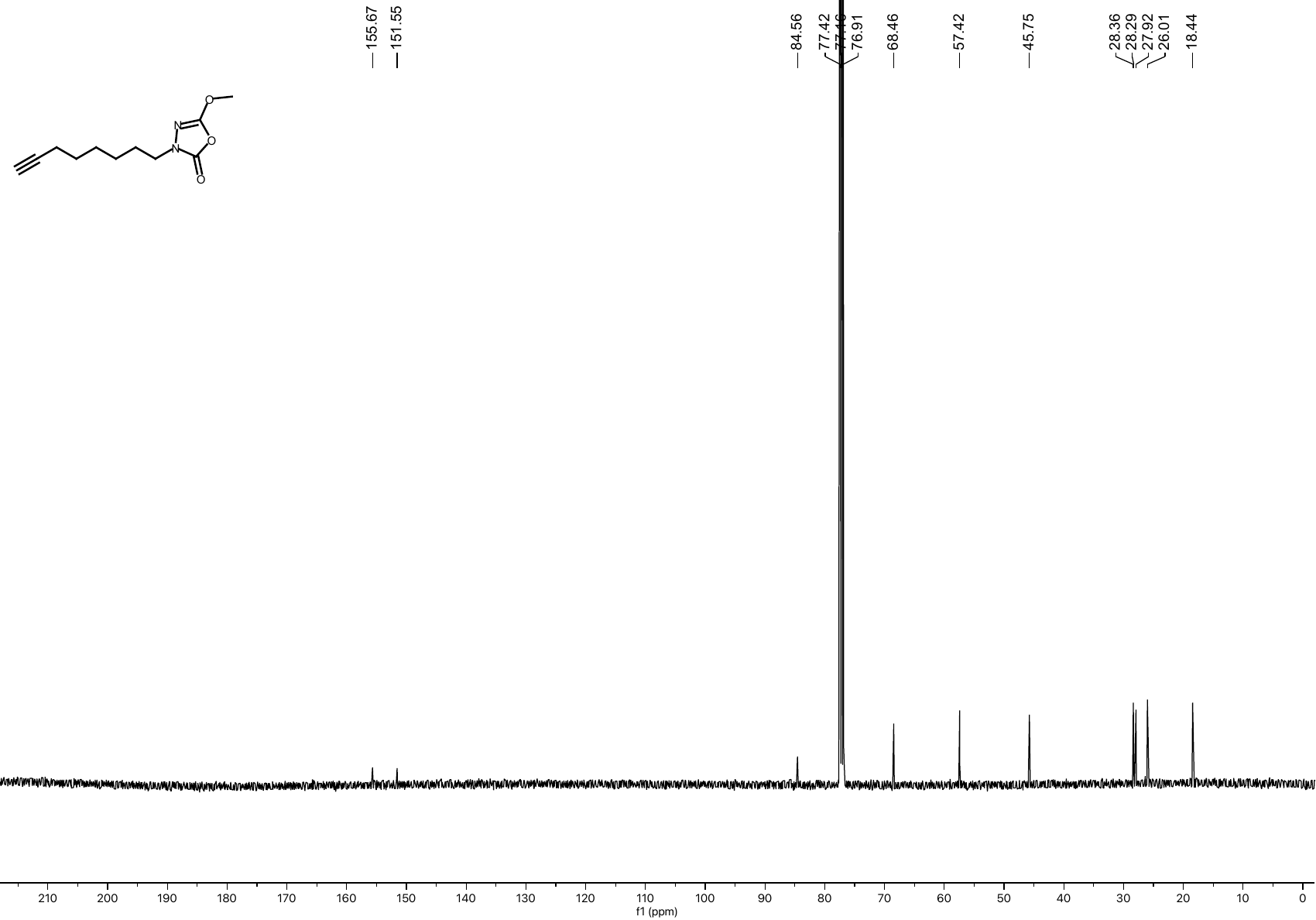
• **^13^C NMR spectrum of compound 7 (CDCl_3_, 126 MHz)**

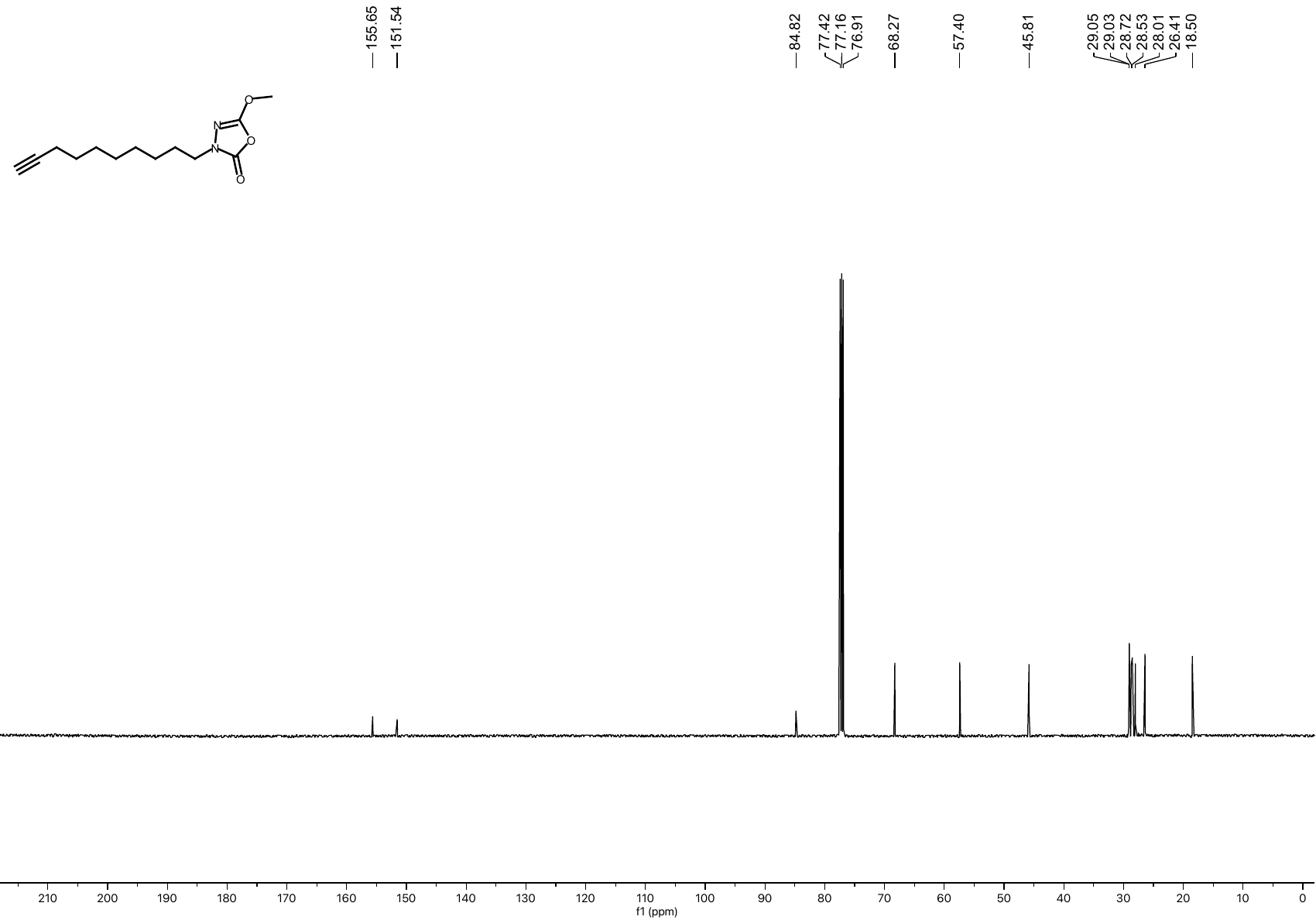
• **^13^C NMR spectrum of compound 8 (CDCl_3_, 126 MHz)**

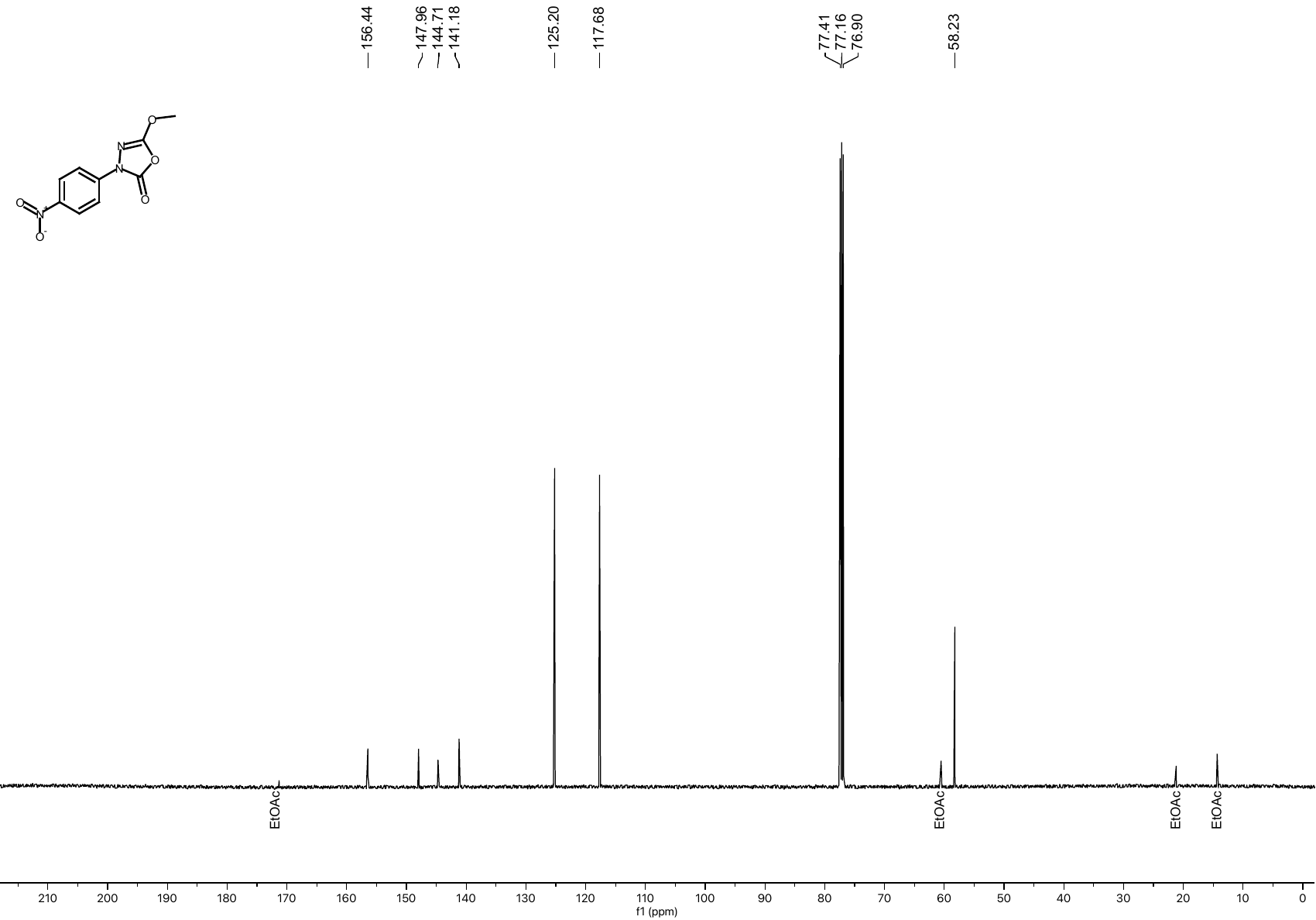
• **^13^C NMR spectrum of compound S10 (CDCl_3_, 126 MHz)**

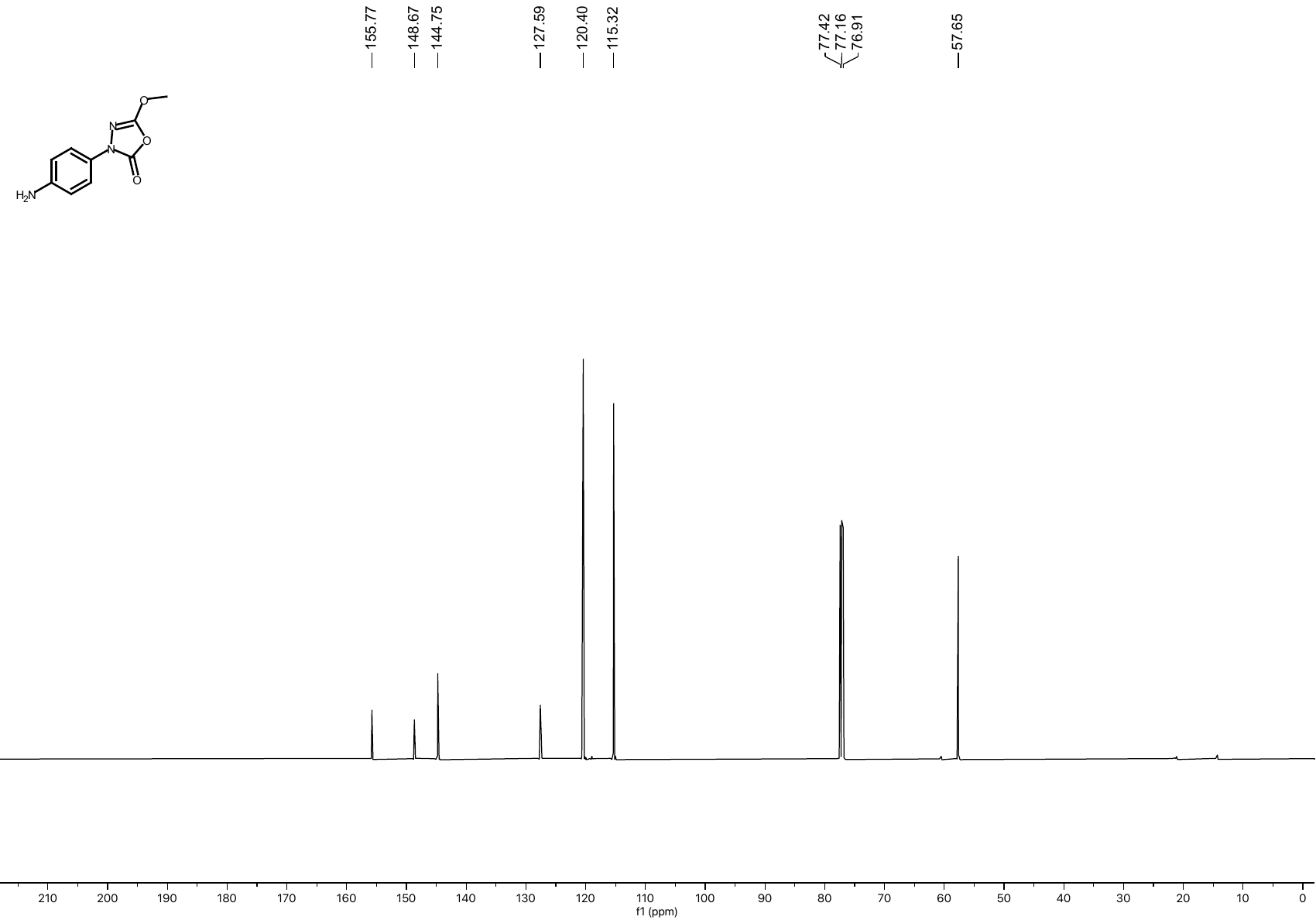
• **^13^C NMR spectrum of compound S10 (CDCl_3_, 126 MHz)**

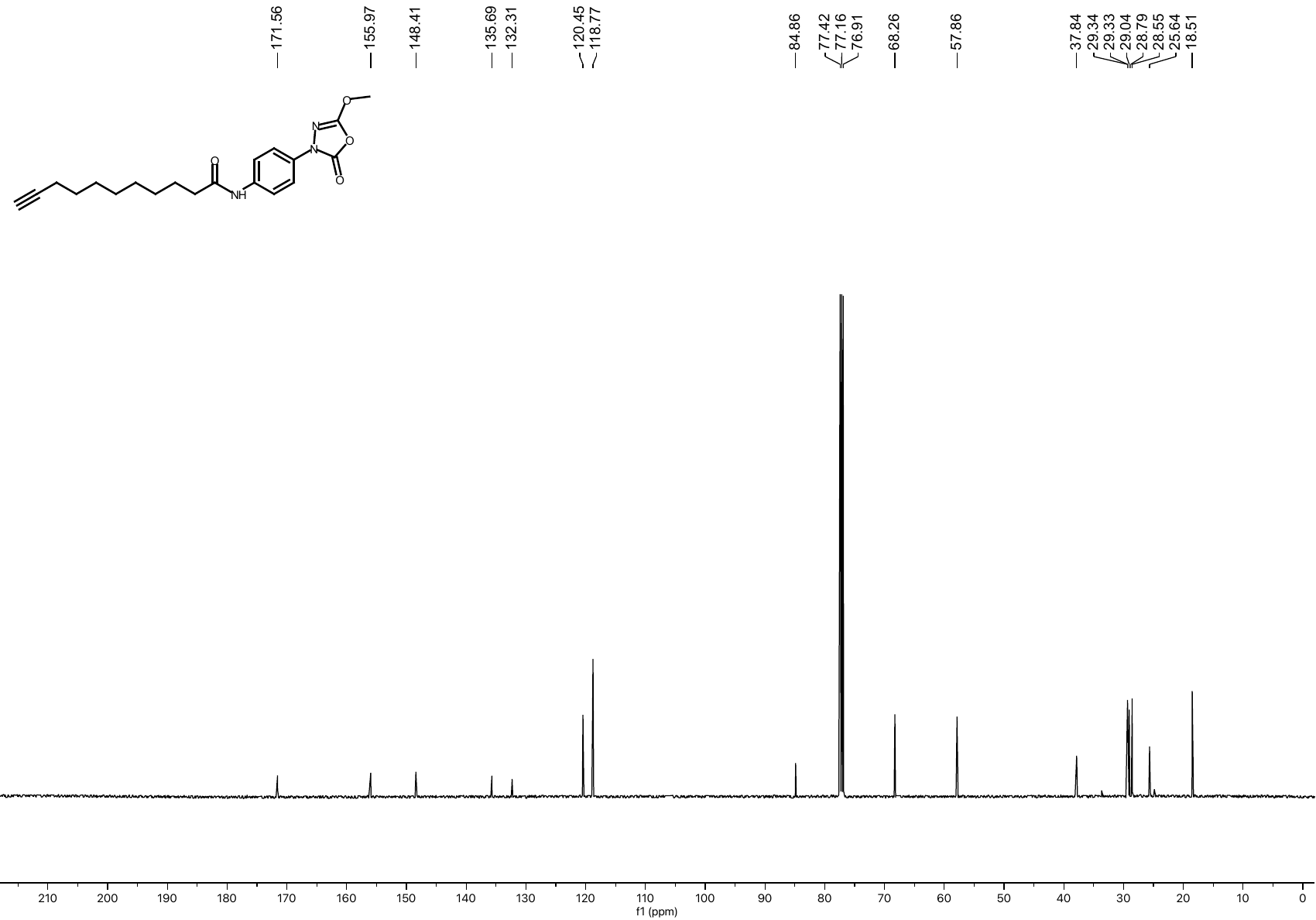
• **^13^C NMR spectrum of compound 9 (CDCl_3_, 126 MHz)**
